# Omitted familial extrinsic risk inflates inferred intrinsic lifespan heritability

**DOI:** 10.64898/2026.04.02.716222

**Authors:** Sergey A. Kornilov

## Abstract

Shenhar et al. (2026) report 50% “intrinsic” lifespan heritability after calibrating a one-component correlated-frailty survival model to Scandinavian twin lifespans. Their framework is mathematically coherent, but the intrinsic component is not identified if heritable, mortality-relevant extrinsic susceptibility is omitted at calibration. We show that one-component calibration absorbs omitted familial extrinsic structure into the intrinsic frailty scale parameter *σ*_*θ*_, and that this variance absorption is visible through separate diagnostics **(1) Variance absorption**. Under misspecification, *σ*_*θ*_ is inflated by +22.1% (95% CI: 21.5–22.7%), corresponding to +49% inflation in 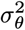. Falconer *h*^2^ is downstream of calibration and inherits a +9.2 pp bias (95% CI: 8.7–9.7). The *σ*_*θ*_ inflation is model-general: +22% (GM), +18% (MGG), +14% (SR); any dependence summary that is strictly increasing in *σ*_*θ*_ inherits this inflation, so Falconer *h*^2^ is one affected downstream quantity among many (Corollary B3). **(2) Structural fingerprint**. In the joint twin survival surface *S*(*t*_1_, *t*_2_), misspecification produces systematic dependence errors (ISE 48× that of the recovery model). Conditional twin dependence is inflated at all ages, peaking at age 80 (Δ*r* = 0.048). **(3) Specificity**. The bias requires an omitted component that is both heritable and mortality-relevant. Three negative controls, a boundary check (*ρ* = 0), and a two-component recovery refit (*σ*_*θ*_ restored to within −3.2%) establish specificity. ACE decomposition yields *C* ≈ 0 throughout: the omitted extrinsic component loads onto *A* (because it is shared 1.0/0.5 in MZ/DZ), so switching summary statistics does not restore identification. **(4) Sensitivity and falsifiability**. Over an empirically anchored regime (*σ*_*γ*_ ∈ [0.30, 0.65], *ρ* ∈ [0.20, 0.50]), Falconer bias ranges from +2.8 to +18.9 pp (mean 9 pp). If *ρ* is sufficiently negative, the bias reverses sign in all three model families (Corollary B4). A full-likelihood robustness check shows that this upward pull is partly structural and partly estimator-specific: in the same misspecified one-component model, ML still inflates *σ*_*θ*_ (+3%), whereas matching only *r*_MZ_ inflates it much more (+21%). These results do not resolve true intrinsic heritability but establish that Shenhar’s 50% estimate carries a structured, model-general upward bias originating in the fitted latent variance *σ*_*θ*_.

## 1 Introduction

### 1.1 The intrinsic heritability estimate

Shenhar et al. (2026) report that the heritability of human lifespan rises from the commonly cited 20–25% to approximately 50% after correcting for extrinsic mortality — deaths from infections, accidents, and environmental hazards. Their method fits parametric mortality models to historical twin cohorts, calibrates genetic variance to reproduce observed twin correlations, then extrapolates to a counterfactual world where the extrinsic hazard *m*_ex_ is set to zero. The implied correction — from observed *h*^2^ ≈ 0.20 to intrinsic *h*^2^ ≈ 0.50 — is large, and any systematic bias in the calibration step propagates directly into the headline estimate.

### 1.2 Two distinct threats

Hamilton (2026) pointed out that infection mortality is heritable and that removing extrinsic mortality introduces selection bias. His simulation demonstrates that conditioning on survival from genetically-patterned extrinsic death inflates Falconer *h*^2^. The biological argument — that susceptibility to severe infection is substantially heritable — is well-supported by the Sørensen et al. (1988) adoption study (RR = 5.81, 95% CI: 2.47–13.7, for infection death) and Danish twin studies estimating 40% heritability of infectious disease mortality (Obel et al., 2010).

However, Hamilton’s simulation does not replicate Shenhar’s procedure. It removes twin pairs where either member died extrinsically (concordant-survivor conditioning), whereas Shenhar does not apply concordant-survivor restriction — they parametrically set *m*_ex_ = 0 in the fitted model. These are different operations and can induce bias through distinct mechanisms (which may coexist):

- **Collider bias** (Hamilton): conditioning on survival creates a genetically non-representative sample.
- **Omitted-variable bias** (this paper): a misspecified model absorbs extrinsic genetic variance into the intrinsic parameter at the calibration step.

### 1.3 Our contribution

Building on Hamilton’s biological insight, and intervening within the correlated-frailty tradition for twin survival analysis (Wienke et al., 2001; 2003; Yashin et al., 1999; Yashin & Iachine, 1995), we identify the bias mechanism specific to Shenhar’s one-component calibration: omitted heritable extrinsic susceptibility is absorbed into the fitted intrinsic frailty scale *σ*_*θ*_, which then propagates to any downstream heritability summary (including Falconer *h*^2^).

We characterize this mechanism through three tiers of evidence. First, we establish the core argument: direct estimation of *σ*_*θ*_ inflation (with a two-component bias decomposition, recovery analysis, and a predictive heuristic; *R*^2^ = 0.988), and independence from concordant-survivor conditioning. Second, we identify structural fingerprints: misfit in the bivariate twin survival surface, age-specific conditional correlations, and failure of alternative summaries (ACE structural equation model) to restore identification. Third, we delineate scope, specificity, and falsifiability via negative controls, dose–response to extrinsic mortality, consistency across three mortality model families (GM, MGG, SR), anchored quantification over an externally informed sensitivity regime deliberately capped below bridge-implied levels, and a sign-reversal prediction under negative genetic correlation.

## 2 Conceptual framework

### 2.1 Shenhar’s estimand and procedure

Shenhar et al. model individual mortality as:

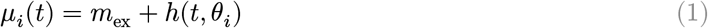

where *m*_ex_ is a cohort-level constant (identical for all individuals) and *θ*_*i*_ are individual intrinsic aging parameters drawn from a Gaussian distribution with standard deviation *σ*_*θ*_. For MZ twins, both members share the same *θ*; for DZ twins, their *θ* values are correlated at *r* = 0.5.

The procedure has two steps:

1. **Calibration:** adjust *σ*_*θ*_ so that simulated MZ twin lifespan correlations match those observed in historical twin data (at historical *m*_ex_).
2. **Extrapolation:** set *m*_ex_ = 0, keep the calibrated *σ*_*θ*_, and recompute Falconer *h*^2^ = 2(*r*_MZ_ − *r*_DZ_).

Under their assumptions — that *m*_ex_ is truly a non-genetic population constant — this extrapolation is mathematically valid.

### 2.2 Omitted heritable extrinsic susceptibility

Suppose the true data-generating process is:

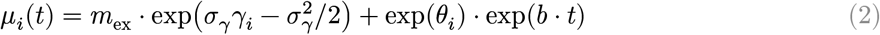

where *γ*_*i*_ ∼ 𝒩(0, 1) is an individual’s genetic susceptibility to extrinsic mortality (the centering term 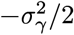 ensures 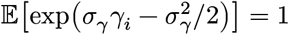, so that 𝔼[*c*_*i*_] = *m*_ex_ for all *σ*_*γ*_) and 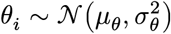 is the intrinsic aging parameter. MZ twins share the same *γ* (and *θ*); DZ twins’ values are correlated at 0.5. The within-individual correlation Corr(*θ*_*i*_, *γ*_*i*_) = *ρ* represents pleiotropy — shared genetic pathways between immune function and intrinsic aging.

Shenhar’s model cannot represent individual extrinsic heterogeneity — in the classic frailty sense of unobserved risk variation shaping mortality dynamics (Vaupel et al., 1979). When the calibration step matches *σ*_*θ*_ to observed *r*_MZ_, it must account for twin concordance contributed by **both** shared intrinsic genetics (through *θ*) **and** shared extrinsic genetics (through *γ*). Since the model has only one source of genetic variation (*σ*_*θ*_), it absorbs both. The calibrated *σ*_*θ*_ is inflated.

Because *θ* and *γ* enter different terms of the hazard, the relationship between fitted and true variances is model-dependent. As a first-order heuristic:

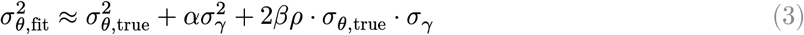

where *α* and *β* are model-dependent coefficients. The *α* = *β* = 1 simplification provides a reasonable first-order approximation (mean absolute residual 8.6%; see §4.3 below). A formal delta-method derivation establishing conditions for inflation is given in Section B (Proposition B1). (Note: Appendix B reparameterizes *θ*_*i*_ = *σ*_*θ*_(*G*_*i*_ + *E*_*i*_) with unit-variance additive-genetic (*G*) and environmental (*E*) components, yielding 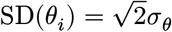; throughout the main text, *σ*_*θ*_ denotes total standard deviation, while the appendix uses *σ*_*θ*_ as a per-component scale factor for derivational convenience. The mapping is 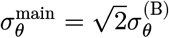. The latent components *G, E* in the proof are distinct from the empirical variance-absorption coefficients *α, β* in Equation (3).)

This is fundamentally a decomposition problem: the intrinsic–extrinsic partition is not identified under the restricted single-component model class (Hougaard, 1995). The calibration step solves 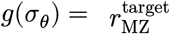 for a functional *g* that maps the dispersion parameter to the predicted MZ twin correlation. Under misspecification, this moment equation converges to a *pseudo-true* parameter 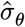 in the sense of (White, 1982) — the best approximation to the true joint MZ/DZ survival structure available within the restricted model class. The key point is that 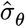 under the misspecified model is not a structural parameter invariant to interventions on *m*_ex_: it is a calibration parameter that absorbs *m*_ex_-driven heterogeneity, so extrapolating to *m*_ex_ = 0 using this inflated value attributes omitted extrinsic structure to intrinsic aging.

Positive pleiotropy (*ρ* > 0) amplifies the inflation and ensures it is robustly upward, because the omitted extrinsic component pushes twin concordance in the same direction as intrinsic frailty. Even at *ρ* = 0, heritable extrinsic frailty alone produces a small but positive bias (§4.8), because the calibration absorbs extrinsic concordance regardless of its genetic correlation with intrinsic aging. The exact magnitude depends on the fitting criterion, but the mechanism is structural (see §3.3 and §4.3 for simulation evidence; Section B for a formal derivation).

To ensure the algebraic integrity of our core claims, the deductive chain from the first-order moment expressions to the final inflation and sign-reversal inequalities (Proposition B1, B2, and Corollary B4) has been fully formalized and mechanically verified using the Lean 4 interactive theorem prover (see Section C). It is important to note the precise epistemic boundary of this formalization: while the algebraic manipulation of the moments is machine-checked to be logically airtight (compiled with zero `sorry`s or unproven assertions), the underlying delta-method linearization and the probability space generating the moment expressions themselves are treated as axiomatized definitions. This verification guarantees that, given the standard asymptotic approximations of survival analysis, the omitted-variable inflation we describe is an inescapable mathematical consequence, free from any hidden algebraic or sign errors.

### 2.3 Distinction from Hamilton’s mechanism

Hamilton demonstrates collider bias from concordant-survivor conditioning — removing extrinsically-dead pairs creates a genetically non-representative sample. Shenhar’s procedure does not do this; our bias enters at calibration and persists with cutoff = 0 (§4.4). The two mechanisms may coexist but are analytically distinct.

## 3 Methods

### 3.1 Simulation overview

We construct a simulation that replicates Shenhar’s calibrate-then-extrapolate procedure under two data-generating processes: one where their assumptions hold, and one where they are violated by heritable extrinsic susceptibility. All results use 50,000 twin pairs per zygosity. We use Gompertz–Makeham as the reference framework because it is the simplest closed-form mortality model that cleanly separates intrinsic Gompertz aging from Makeham extrinsic mortality, making it the most transparent baseline for the oracle, decomposition, and controls before testing generality in Shenhar’s MGG and SR models (§4.9). Lifespans are simulated by inverting the cumulative hazard via the Lambert W function in closed form.

### 3.2 Arms and controls

Our experimental design follows a structured hierarchy: a **reference oracle, primary conditions** that test the central hypothesis against its correctly specified baseline, **negative controls** that isolate necessary conditions, a **recovery analysis** that confirms recoverability, **diagnostic checks**, and **sweeps** that probe boundary behavior. Table 2 summarizes all conditions; the subsections below describe each in detail.

**Table 1.**
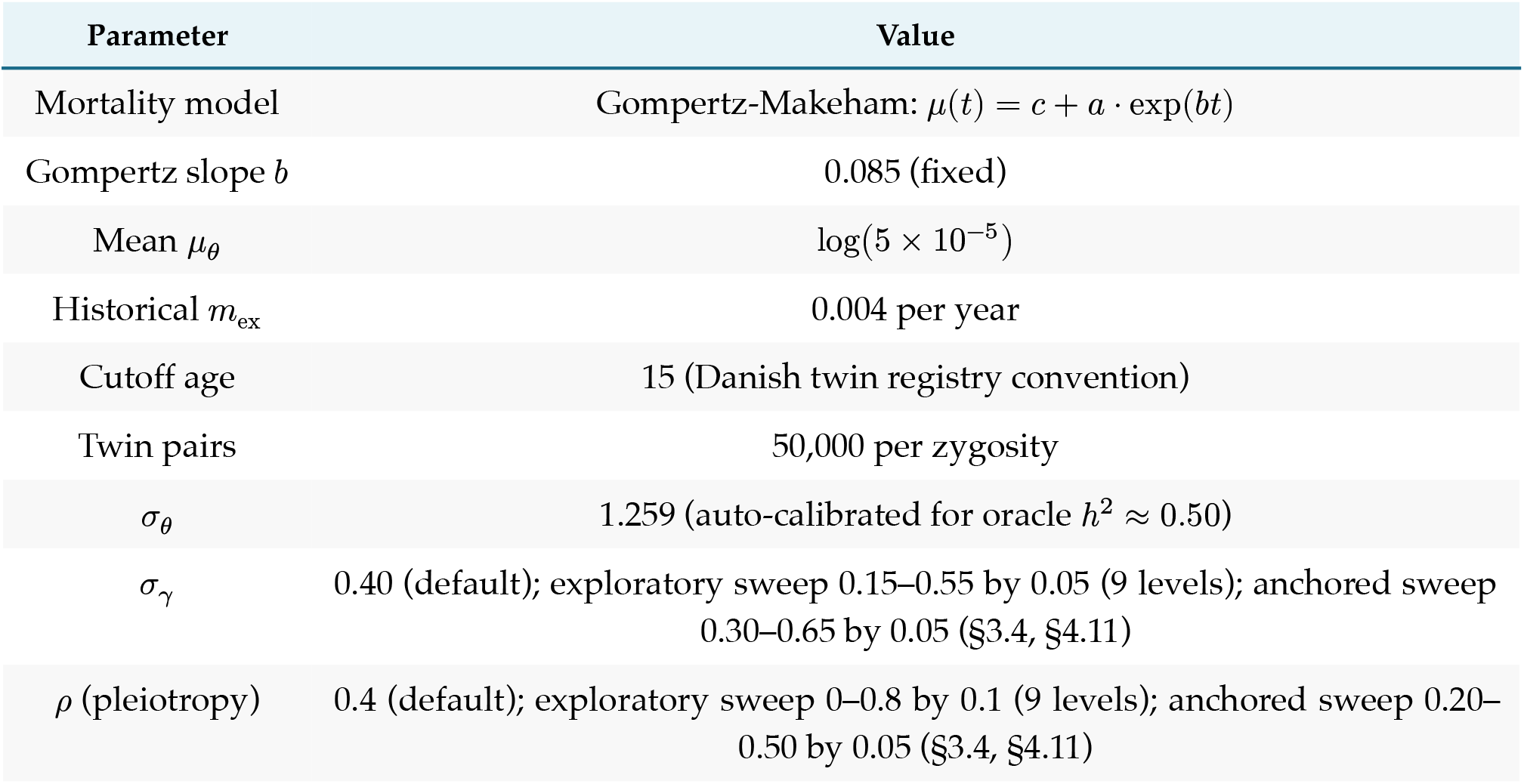
Simulation parameters matching Danish cohorts born 1870–1900.

| Parameter | Value |
| --- | --- |
| Mortality model | Gompertz-Makeham: $\mu(t) = c + a \cdot \exp(bt)$ |
| Gompertz slope $b$ | 0.085 (fixed) |
| Mean $\mu_\theta$ | $\log(5 \times 10^{-5})$ |
| Historical $m_{\text{ex}}$ | 0.004 per year |
| Cutoff age | 15 (Danish twin registry convention) |
| Twin pairs | 50,000 per zygosity |
| $\sigma_\theta$ | 1.259 (auto-calibrated for oracle $h^2 \approx 0.50$) |
| $\sigma_\gamma$ | 0.40 (default); exploratory sweep 0.15–0.55 by 0.05 (9 levels); anchored sweep 0.30–0.65 by 0.05 (§3.4, §4.11) |
| $\rho$ (pleiotropy) | 0.4 (default); exploratory sweep 0–0.8 by 0.1 (9 levels); anchored sweep 0.20–0.50 by 0.05 (§3.4, §4.11) |

**Table 2.** Full simulation design. All conditions use 50,000 twin pairs per zygosity. *σ*_*γ*_, *ρ*, and heritability refer to the **data-generating process**; the fitted model is always Shenhar’s one-component specification except for Recovery. Misspecified row is highlighted.

| Condition | $\sigma_\gamma$ | $\rho$ | $m_{\text{ex}}$ | Heritable? | Tests |
| --- | --- | --- | --- | --- | --- |
| Oracle | 0 | — | 0 | — | Ground-truth intrinsic $h^2$ |
| Baseline: correctly specified | 0 | — | 0.004 | — | Pipeline consistency under correct model |
| Misspecified: omitted familial extrinsic | 0.40 | 0.4 | 0.004 | Yes | Central hypothesis — variance absorption |
| Recovery: two-component refit | 0.40 | 0.4 | 0.004 | Yes | Positive control — model recovers $\sigma_\theta$? |
| Check: no survivor cutoff | 0.40 | 0.4 | 0.004 | Yes | Boundary — requires survival conditioning? |
| Check: pleiotropy only | 0.40 | 0 | 0.004 | Yes | Isolates $\rho = 0$ from $\rho > 0$ effect |
| Control: nonfamilial extrinsic | 0.40 | 0 | 0.004 | No | Specificity — requires genetic concordance? |
| Control: irrelevant trait ($\zeta$) | — | $\rho_\zeta = 0.4$ | 0.004 | Yes | Specificity — trait must enter hazard? |
| Control: vanishing extrinsic | 0.40 | 0.4 | 0.0002 | Yes | Boundary — requires extrinsic signal? |
| Comparator: Hamilton | 0.40 | 0.4 | 0.004 | Yes | Collider bias comparison (different procedure) |

#### 3.2.1 Reference

##### Oracle

The oracle runs the pure intrinsic DGP (*σ*_*γ*_ = 0, *m*_ex_ = 0) using the true *σ*_*θ*_ = 1.259, calibrated to yield *h*^2^ ≈ 0.50. No calibrate-then-extrapolate pipeline is applied — the oracle evaluates Falconer *h*^2^ directly from intrinsic-only twin correlations, providing the ground-truth benchmark (*h*^2^ = 0.51) against which all other conditions are measured.

#### 3.2.2 Primary conditions

##### Baseline — Correctly specified

Data are generated under Shenhar’s own assumptions: extrinsic mortality is a population-level constant (*c*_*i*_ = *m*_ex_ = 0.004 for all individuals), with no individual heterogeneity (*σ*_*γ*_ = 0). The model is **correctly specified**. We calibrate *σ*_*θ*_ to match the simulated *r*_MZ_ via 40-iteration stochastic bisection, then extrapolate to *m*_ex_ = 0. Any deviation from the oracle reflects the estimand mismatch between “intrinsic” (no extrinsic mortality present) and “extrapolated” (extrinsic mortality present during calibration, then removed). This condition validates that the calibrate-then-extrapolate logic is internally consistent.

##### Misspecified — Omitted familial extrinsic (main result)

Data are generated under the two-component DGP (Equation (2)): extrinsic mortality is individually heterogeneous (*σ*_*γ*_ = 0.40, *ρ* = 0.4), heritable, and assigned at the individual level. Shenhar’s **one-component** model — which lacks any *γ* parameter — is then fit by the same calibration pipeline as the Baseline. This is the central test: does omitting heritable extrinsic susceptibility from the model inflate the intrinsic estimate?

#### 3.2.3 Negative controls

##### Control — Nonfamilial extrinsic

Data include individual extrinsic heterogeneity (*σ*_*γ*_ = 0.40), but *γ* values are drawn **independently** for each twin rather than shared within pairs (`gamma_heritable = FALSE`). The extrinsic component adds individual mortality variance but no **genetic** concordance. If the calibration bias requires within-pair correlation in extrinsic susceptibility, this control should show no inflation.

##### Control — Irrelevant trait (*ζ*)

heritable latent trait *ζ* is genetically correlated with *θ* at the same level as *γ* in the Misspecified condition, but *ζ* does not enter the hazard function — it has no effect on mortality. This tests whether shared genetic architecture **per se** is sufficient to bias calibration, or whether the correlated trait must affect mortality through the hazard.

##### Control — Vanishing extrinsic

Data include heritable extrinsic susceptibility (*σ*_*γ*_ = 0.40, *ρ* = 0.4) — the same two-component DGP as the Misspecified condition — but with negligible extrinsic mortality magnitude (*m*_ex_ = 0.0002, one-twentieth of historical levels). This tests the boundary condition: when the extrinsic hazard channel carries negligible mortality signal, the bias should vanish regardless of how strongly extrinsic susceptibility is heritable.

##### Check — Pleiotropy only (*ρ* = 0)

Same DGP as the Misspecified condition except *ρ* = 0: extrinsic susceptibility is heritable and individually heterogeneous, but genetically **independent** of intrinsic frailty. This isolates the contribution of pure extrinsic concordance from the amplifying effect of pleiotropy.

#### 3.2.4 Recovery analysis

##### Recovery — Two-component refit

Data are generated under the full two-component DGP (same as the Misspecified condition), but calibration uses a model that **includes** the extrinsic frailty component: (*σ*_*γ*_, *ρ*) are held at their true DGP values, and only *σ*_*θ*_ is estimated via bisection on *r*_MZ_. If the bias is genuinely caused by omitting *γ* from the model, then including it should recover the true *σ*_*θ*_ and eliminate the inflation component, leaving only the attenuation that arises under correct specification. This is the critical positive control — the mirror image of the Misspecified condition.

#### 3.2.5 Diagnostic checks and sweeps

Several additional analyses probe boundary behavior and dose-response (full results in §4):

- **Check — No survivor cutoff:** Repeats the Misspecified condition with no age cutoff — all twin pairs included, including infant deaths. Tests whether the bias requires concordant-survivor conditioning.
- **Dose-response:** Sweeps *m*_ex_ from 0 to 0.012 (3× historical) with all other parameters at Misspecified values.
- **Negative** *ρ* **sweep:** Sweeps the genetic correlation *ρ* from −0.60 to +0.80, testing the prediction that negative *ρ* reverses the bias sign.
- *m*_ex_ **split:** Decomposes *m*_ex_ into heritable and non-heritable fractions, sweeping the heritable share from 0 to 100%.

### 3.2.6 Comparator

#### Comparator — Hamilton

Hamilton’s procedure removes twin pairs where either member died extrinsically before a threshold age (concordant-survivor restriction). We implement this as a separate condition for comparison: data follow the same two-component DGP as the Misspecified condition, but instead of calibrating a parametric model, we compute Falconer *h*^2^ directly from the surviving concordant subsample. This represents a different analytical procedure and a different bias mechanism (collider bias from sample restriction rather than variance absorption from calibration).

### 3.3 Calibration and estimands

The primary calibration uses stochastic bisection on Monte Carlo estimates of *r*_MZ_ (40 iterations). This matches Shenhar et al.‘s procedure: their supplement explicitly states that “parameter heterogeneity was adjusted to match MZ twin lifespan correlations by varying the degree of parameter variation” — confirming single-target *r*_MZ_ calibration. They also report Falconer *h*^2^ = 2(*r*_MZ_ − *r*_DZ_), making our summary statistic directly comparable to their published estimates. Falconer *h*^2^ summarizes twin concordance under classical ACE assumptions and does not, without further conditions, identify a unique causal variance ratio (it is sensitive to assortative mating, epistasis, and gene–environment correlation); we adopt it here strictly for comparability with Shenhar et al.’s reported estimates.

A natural concern is whether calibrating to *r*_MZ_ alone understates the information available — perhaps a joint (*r*_MZ_, *r*_DZ_) calibration would behave differently. We tested this directly: under the misspecified DGP, the model fitted to *r*_MZ_ also predicts *r*_DZ_ with near-zero discrepancy (0.0002). A joint calibration targeting both correlations yields *σ*_fit_ = 1.552 and bias = +9.3 pp — slightly larger than the *r*_MZ_-only result (*σ*_fit_ = 1.537, bias = +7.6 pp), consistent with the additional *r*_DZ_ target pulling *σ*_*θ*_ further upward to accommodate genetic concordance from both zygosity groups simultaneously. This shows that adding *r*_DZ_ information does not resolve the absorption — and if anything slightly amplifies it: the misspecified model’s single genetic parameter must account for all genetic concordance regardless of whether one or two correlation targets are used. Full-likelihood fitting can improve diagnostics (e.g., posterior predictive checks on bivariate survival surfaces), but it cannot recover a decomposition that the model does not parameterize; closely related sensitivity of heterogeneity estimates to frailty-model misspecification is documented in shared frailty survival models by (Gasparini et al., 2019). Under an underparameterized one-component model, maximum likelihood will still select the parameter value that best approximates the true two-component data within the restricted model class — i.e., a pseudo-true *σ*_*θ*_ that absorbs part of the omitted heritable extrinsic component. Indeed, starting with the correlated frailty framework introduced by Yashin & Iachine (1995), maximum-likelihood correlated gamma-frailty models have been applied to Scandinavian twin cohorts; Wienke et al. (2003) estimate frailty heritability ≈ 0.58 in the same Danish cohorts, confirming that the phenomenon is not an artifact of our specific calibration criterion.

### 3.4 Anchoring *σ*_*γ*_ and *ρ* to independent evidence

#### Anchor for *σ*_*γ*_

Obel et al. (2010) report liability-scale heritability 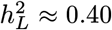 (95% CI: 0.12–0.50) for infection mortality in 44,005 Danish twin pairs. This is consistent with the substantial familial aggregation reported by Sørensen et al. (1988 RR = 5.81 for adoptee infection death) and extended by Petersen et al. (2010), who show genetic influences on both infection incidence and case fatality in Danish adoptees. As an order-of-magnitude anchor, 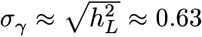 at 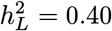. This mapping is approximate — liability-scale and log-hazard dispersion are not on identical scales — and we evaluate robustness to this translation by sweeping *σ*_*γ*_ ∈ {0.30, 0.40, 0.55, 0.63}.

Corroborating evidence comes from correlated frailty models fit directly to Scandinavian twin survival data: Wienke et al. and Yashin et al. estimate frailty heritabilities of ≈ 0.58 (Danish twins) and comparable magnitudes (Swedish twins), respectively (see also §3.3). These estimates reflect total unobserved frailty, not an extrinsic channel per se, but they confirm that substantial familial heterogeneity in mortality susceptibility exists in the same cohort context — supporting the empirical plausibility of our sensitivity range.

##### Anchor for *ρ*

Qiu (2025) report genome-wide genetic correlations between COVID-19 severity and longevity of *r*_*g*_ ≈ −0.34 to −0.35, corroborated by Mendelian randomization (Ying et al., 2021 genetically-proxied 10-year lifespan increase reduces COVID-19 infection risk, OR = 0.31). The shared architecture is driven by conserved inflammatory pathways (IL-6/IL-6R, TNF-*α*, NF-*κ*B, FOXO3, APOE, TLR) with documented pleiotropic effects on immune function and cellular senescence (Franceschi et al., 2000; Rosa et al., 2019). Because our *θ* parameterizes log-hazard (higher *θ* increases mortality, decreasing longevity), a negative genetic correlation between **longevity** and infection severity corresponds to a **positive** correlation between the two mortality-hazard components — i.e., Corr(severity, *θ*) = −Corr(severity, longevity), hence *ρ* > 0 in our notation. We adopt *ρ* ∈ [0.20, 0.50]. COVID-19 represents a single novel pathogen and serves as an existence proof that genetic correlation between infection susceptibility and aging pathways exists — not as a magnitude anchor for the preantibiotic Danish cohorts, whose mortality was dominated by tuberculosis, pneumonia, diphtheria, and enteric infections with different immunological mechanisms. Stronger evidence comes from GWAS: Schurz et al. (2024) estimate the SNP-heritability of TB susceptibility at 26.3% (95% CI 23.7–29.0%) with the only genome-wide significant signal in the HLA class II region (HLA-DQA1/DRB1). Independently, Joshi et al. (2017) identify HLA-DQA1/DRB1 as genome-wide significant for parental lifespan — a proxy longevity phenotype — in 606,059 individuals. The convergence of signals in the same HLA class II region across the two traits is consistent with shared immunogenetic architecture linking infection susceptibility to aging-related mortality, though formal colocalization has not been performed. In Nordic populations specifically, HLA-DQB1*05:01 is protective for TB (OR = 0.81; (Tervi et al., 2023)), and the Sørensen adoption study found RR = 5.81 (95% CI: 2.47–13.7) for infection death among biological relatives — concentrated in case-fatality rather than incidence, with evidence for non-additive genetic effects (Petersen et al., 2010; Sørensen et al., 1988). The shared inflammatory architecture (IL-6/IL-6R, TNF-*α*, NF-*κ*B, FOXO3, APOE, TLR) connects immune function to cellular senescence across pathogen classes (Franceschi et al., 2000; Rosa et al., 2019). No direct LD-score regression estimate between pre-antibiotic infection mortality and longevity exists; our *ρ* range is informed by the convergence of these indirect lines of evidence, not calibrated to any single estimate. The true genetic correlation between **general** historical extrinsic susceptibility and intrinsic aging may differ from the COVID-19-specific estimate in either direction.

##### Bridging *σ*_*γ*_ to liability-scale 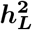

The mapping 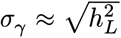 is heuristic — liability-scale heritability depends on prevalence and the threshold model, while *σ*_*γ*_ parameterizes a continuous log-hazard frailty. To ground the translation, we run the full competing-risks DGP at each value of *σ*_*γ*_, generate MZ and DZ twin pairs, create a binary indicator for whether each twin died of infection, compute tetrachoric correlations from the resulting 2 × 2 tables, and derive Falconer’s 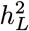 (Figure 12). Under our assumed hazard structure, matching Obel et al.‘s 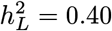 requires *σ*_*γ*_ ≈ 1.47 — more than double our upper bound of 0.65. We treat this bridge as directional evidence — confirming that published infection-mortality heritability implies substantial extrinsic heterogeneity, and that our sensitivity range sits well below the empirically implied parameterization — but not as a calibration target for *σ*_*γ*_. The bridge inherits the DGP’s simplifications (single competing risk, stylized hazard structure, liability-threshold mapping), so the specific *σ*_*γ*_ value it implies should not be taken at face value. What it does establish is the sign and relevance of the effect: extrinsic heterogeneity of the magnitude implied by published epidemiological data would produce biases at least as large as those we report.

##### Externally informed restricted sensitivity regime

*σ*_*γ*_ ∈ [0.30, 0.65], *ρ* ∈ [0.20, 0.50]. We cap *σ*_*γ*_ at 0.65 — well below the bridge-implied value of ≈ 1.47 — so that the reported biases reflect what the stylized DGP produces under cautious assumptions. We do not treat the bridge as a calibration target. Under 10-seed Monte Carlo averaging, the bridge maps Obel’s central estimate 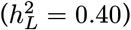 to *σ*_*γ*_ ≈ 1.35 (95% CI: 1.35 ± 0.04); the lower bound of Obel’s CI 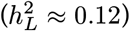 maps to *σ*_*γ*_ ≈ 0.55, which falls within the anchored range. Thus our sensitivity ceiling sits at or below the empirically implied parameterization across the full Obel CI, and well below the point estimate. An extended stress test (§4.11, Figure 12 panel B) confirms that bias continues to increase monotonically beyond this ceiling; the formal proof’s equal-noise approximation (Proposition B1) also becomes quantitatively unreliable in that regime. The correlated-frailty literature (Wienke et al., 2003; Yashin et al., 1999) independently supports the existence of substantial familial heterogeneity in these cohorts, reinforcing the qualitative conclusion that the sensitivity range sits below empirically motivated levels.

### 3.5 Bivariate survival diagnostics

To complement univariate fits and scalar concordance summaries, we assessed model adequacy using the full *bivariate survival surface*

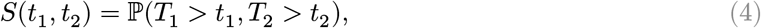

evaluated for twin lifespans (*T*_1_, *T*_2_). This surface encodes dependence across the entire age range: for fixed *t*_2_, the function *t*_1_ ↦ *S*(*t*_1_, *t*_2_) is the survival curve of twin 1 among pairs for which twin 2 survives past *t*_2_. Under independence, *S*(*t*_1_, *t*_2_) = *S*_1_(*t*_1_)*S*_2_(*t*_2_); positive dependence (in the usual positive quadrant dependence sense) manifests as *S*(*t*_1_, *t*_2_) > *S*_1_(*t*_1_)*S*_2_(*t*_2_), with the strongest deviations typically in the late-life region where shared frailty amplifies concordance. The bivariate survival function is a classical object in survival analysis (Clayton, 1978; Hougaard, 2000; Oakes, 1989); its use as a diagnostic for calibration misspecification in Shenhar’s specific framework is our contribution.

#### Empirical estimator

Given *n* twin pairs with lifespans 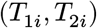 (after cutoff filtering; all simulated lifespans are exact event times with no censoring), we estimated

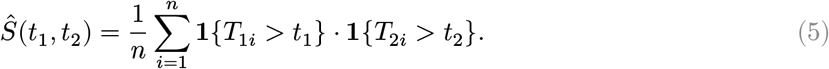

On a uniform grid 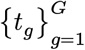 with spacing Δ*t*, this is computed vectorially via indicator matrices *I*^(*k*)^ ∈ {0, 1}^*n*×*G*^ with 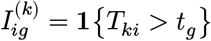, yielding *Ŝ* = (*I*^(1)^) *I*^(2)^ /*n* in a single matrix multiply. Marginal survival estimates are *Ŝ* _*k*_(*t*) = (1/*n*) ∑_*i*_ **1**{*T*_*ki*_ > *t*}. We used Δ*t* = 2 years over [20, 95] (*G* = 38).

#### Surface-based goodness-of-fit metrics

To quantify misfit between a fitted model’s surface and the true DGP, we used three complementary metrics. **(i) Integrated squared error (ISE):** ISE = *∬* (*S*_true_ − *S*_fit_)^2^*dt*_1_ *dt*_2_, approximated by a Riemann sum 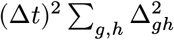. **(ii) Supremum deviation:** 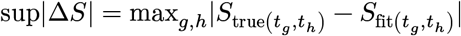, sensitive to localized ridges of misfit. **(iii) Cramér–von Mises type:** 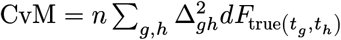, where 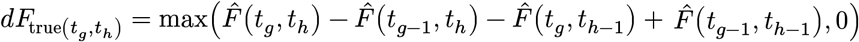 are bivariate CDF rectangle increments (clipped at zero to handle finite-sample noise) that down-weight sparse tail regions. The clipping is negligible: empirically ∑ *dF* < 1 by less than 0.1%.

#### Concordance curve

The age-specific concordance probability

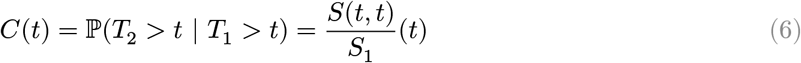

summarizes diagonal dependence while preserving age structure. This is related to the dependence ratio 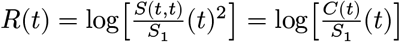 used in §4.5; the two contain equivalent information on different scales. Standard errors are computed via pair bootstrap (200 replicates resampling twin pairs, not individuals) to correctly account for within-pair correlation.

#### Cross-ratio (Oakes 1989)

The cross-ratio function (Oakes, 1989)

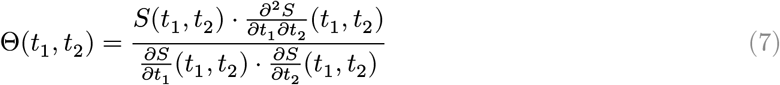

characterizes *local* dependence. (We use capital Θ for the cross-ratio to distinguish it from the individual frailty parameter *θ*_*i*_.) Here *∂*^2^*S*/*∂t*_1_*∂t*_2_ = *f*(*t*_1_, *t*_2_) ≥ 0 is the bivariate density, while *∂S*/*∂t*_*k*_ are the partial derivatives of the *joint* survival function — not the marginal densities. (At (*t*_1_, *t*_2_), 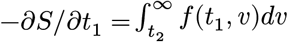 is the sub-density of *T*_1_ given *T*_2_ > *t* _2_, which differs from the marginal density *f*_1_ (*t*_1_) whenever dependence is present.) Under independence Θ = 1; under positive dependence Θ > 1. We evaluated Θ(*t, t*) on the diagonal via central finite differences of the empirical joint survival surface on a 2-year grid (interior points only; grid edges are set to NA). All simulated lifespans are exact event times (no censoring); grid points where *Ŝ* (*t, t*) < 0.01 or the estimated partial derivatives fall below 10^−6^ in absolute value are masked to avoid numerical instability. Under independence, *S*(*t, t*) = *S*_1_(*t*)^2^ and all dependence measures equal their null values (*R*(*t*) = 0, Θ(*t, t*) = 1); we verified this calibration property holds in uncorrelated control simulations. In all comparisons, *S*_true_ denotes the empirical surface from a 50,000-pair simulation under the true DGP (Monte Carlo truth), not an analytic expression.

#### Age-specific conditional correlation

As an additional summary of age-varying association, we computed cor(*T*_1_, *T*_2_ | *T*_1_ > *t, T*_2_ > *t*) — the Pearson and Spearman correlations of lifespans among pairs surviving past age *t*. This statistic is not copula-invariant (unlike the cross-ratio or Kendall’s *τ*) but is directly interpretable and shows how the misspecified model distorts the age-pattern of twin concordance. Minimum pair counts of 100 were required for stable estimates.

### 3.6 Reproducibility

All simulation code, pipeline targets, and data needed to reproduce the analyses in this paper are available at https://github.com/biostochastics/extrinsic-frailty-heritability/.

## 4 Results

The misspecification bias has a rich internal structure. We organize the evidence into three tiers, corresponding to the cumulative logic of a structured methodological critique: each tier addresses a distinct question about the mechanism threatening Shenhar et al.‘s 50% intrinsic heritability estimate.

### Tier 1 — Core argument (§4.1–4.4)

What is inflated, by how much, and is it recoverable? We lead with the primary estimand (*σ*_*θ*_ inflation), demonstrate the two-component bias decomposition and recovery, trace the variance-absorption mechanism, and establish independence from Hamilton’s concordant-survivor conditioning.

### Tier 2 — Structural fingerprints (§4.5–4.7)

Does the absorption leave observable traces beyond the scalar parameter? We show that the misspecified model fails to reproduce the bivariate twin survival surface, produces systematic age-specific distortions in conditional twin correlations, and that switching from Falconer *h*^2^ to ACE structural equation modeling does not restore identification.

### Tier 3 — Scope, specificity, and falsifiability (§4.8–4.13)

How general is the mechanism, what are its necessary conditions, and can it be falsified? We establish that the bias requires all three ingredients (genetic concordance, mortality relevance, extrinsic hazard magnitude), demonstrate model-generality across mortality families and extrinsic frailty constructions, quantify the dose-response and anchored magnitude, confirm a sign-reversal prediction under negative pleiotropy, and show that the bias is partly structural and partly estimator-specific (ML vs. moment calibration).

#### Epistemic hierarchy

The logical structure of the argument is: Proposition B1 (Section B) establishes inflation of the calibrated intrinsic frailty parameter 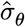 under first-order approximation; Corollary B3 shows that this upstream distortion propagates to *any* downstream dependence summary monotone in *σ*_*θ*_ — Falconer *h*^2^ being one such summary (Proposition B2), but not privileged. The bivariate survival surface and age-specific conditional correlations (Tier 2) are *diagnostic consequences* of the parameter inflation: they show that the absorption leaves structured, observable traces in the joint twin survival structure, but they do not independently prove the mechanism. The ACE decomposition (§4.7) is a *sensitivity and translation layer*: it demonstrates that switching the summary vocabulary from Falconer to structural equation modeling does not restore identification. The cause of the bias is calibration-stage variance absorption (Tier 1); diagnostics and ACE are downstream evidence of its scope.

All estimates are Monte Carlo averages with *t*-based 95% CIs. Table 3 maps each claim to its supporting evidence.

**Table 3.** Exhibit index. Each row maps a claim to the sections, figures, and tables that substantiate it, along with the evidence type. Tier 1 (§4.1–4.4) establishes the core argument; Tier 2 (§4.5–4.7) identifies structural fingerprints; Tier 3 (§4.8–4.13) delineates scope, specificity, and falsifiability.

| Claim | Evidence (§ / Fig / Table) | Type |
| --- | --- | --- |
| $\sigma_\theta$ inflated +22.1% under mis-specification | §4.1; Table 4; Figure 2 | Simulation |
| Inflation decomposes into attenuation + absorption; recovery eliminates inflation | §4.2; Table 5; Figure 1 | Simulation + positive control |
| Absorption tracks omitted-variable prediction ($R^2 = 0.988$) | §4.3; Figure 3 | Simulation + formal |
| Bias persists without concordant-survivor restriction | §4.4 | Simulation |
| Bivariate surface misfit ($48\times$ ISE ratio) | §4.5; Figure 4; Figure 5; Figure 6 | Diagnostic |
| Age-specific conditional correlation excess ($\Delta r = 0.048$ at age 80) | §4.6; Figure 7 | Diagnostic |
| ACE yields $C \approx 0$; bias loads onto $A$, not $C$ | §4.7; Figure 8 | Diagnostic |
| Bias requires genetic concordance + mortality pathway + hazard signal | §4.8; Table 5 | Simulation |
| Bias is model-general (GM, MGG, SR) and construction-robust | §4.9; Figure 9; Table 7; Figure 17 | Simulation |
| Monotonic dose-response to $m_{\text{ex}}$ and heritable fraction | §4.10; Figure 10 | Sensitivity |
| Anchored regime: +2.8 to +18.9 pp | §4.11; Figure 11; Figure 12 | Sensitivity |
| Negative $\rho$ reverses bias sign in all 3 model families | §4.12; Figure 13 | Formal + simulation |
| ML still inflates $\sigma_\theta$ (+3%); moment matching amplifies $\approx 7\times$ | §4.13; Figure 14 | Robustness |

**Tier 1: Core argument** — What is inflated, by how much, and is it recoverable?

### 4.1 Primary estimand: intrinsic frailty parameter inflation

Our central claim concerns variance absorption under survival-model misspecification rather than the validity of Falconer’s summary statistic per se. Accordingly, we report inflation of the fitted intrinsic frailty parameter *σ*_*θ*_ as the primary estimand, with Falconer *h*^2^ as a secondary comparability metric to Shenhar et al..

The estimand ladder is: calibration under misspecification → pseudo-true 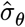 (primary) → counterfactual intrinsic lifespan distribution → reported *h*^2^ (secondary). The misspecification enters at the first step; everything downstream inherits the inflation.

We define three scalar estimands for the inflation. Let *σ*_*θ*,0_ denote the true intrinsic dispersion under the data-generating process, and 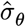 the pseudo-true value to which the calibrated parameter converges under misspecification:

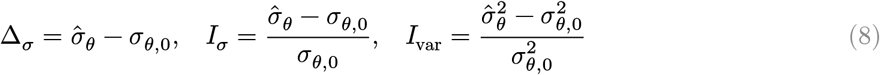

Δ_*σ*_ is the absolute shift in the frailty scale; *I*_*σ*_ is the relative inflation (reported as a percentage throughout this paper); *I*_var_ is the variance-scale inflation, which is the relevant scale for downstream summaries that depend on 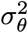. Note *I*_var_ = (1 + *I*_*σ*_)^2^ − 1: a scale inflation of +22% (*I*_*σ*_ = 0.22) corresponds to a variance inflation of +49% (*I*_var_ = 0.49), which explains why a seemingly moderate *σ*_*θ*_ shift produces a large *h*^2^ bias. Under the equal-noise approximation (Proposition B1, Section B; note that Appendix B uses the per-component scale 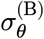 with 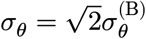, but the sign conclusion is invariant to this reparameterization), *I*_*σ*_ > 0 whenever *ρ* ≥ 0 and *σ*_*γ*_ > 0; by Corollary B3, any dependence summary monotone in *σ*_*θ*_ inherits the bias direction.

Table 4 reports *σ*_*θ*_ inflation across all three mortality model families. Under the default DGP (*σ*_*γ*_ = 0.40, *ρ* = 0.4), the fitted intrinsic dispersion is inflated by +22.1% (GM), +17.9% (MGG), and +14.1% (SR) — translating to variance inflation (*I*_var_ = (1 + *I*_*σ*_)^2^ − 1) of +49%, +39%, and +30% respectively. The correctly specified two-component recovery model restores *σ*_*θ*_ to within −3.2% of the true value, confirming that the inflation is specifically attributable to the omitted extrinsic component.

**Table 4.** Intrinsic frailty parameter inflation (primary estimand) across mortality model families. *σ* inflation = 100 × (*σ*_fit_/*σ*_true_ − 1)%. Falconer *h*^2^ bias is shown for comparability with Shenhar et al. All values are single-seed (MASTER_SEED = 66829); see text for 20-seed Monte Carlo uncertainty.

| Model | $\sigma_{\text{true}}$ | $\sigma_{\text{fit}}$ | $\Delta\sigma_{\theta}$ | $\sigma$ inflation | Falconer bias |
| --- | --- | --- | --- | --- | --- |
| GM (ref.) | 1.259 | 1.537 | +0.278 | **+22.1%** | +7.6 pp |
| MGG | 0.270 | 0.318 | +0.048 | **+17.9%** | +9.2 pp |
| SR | 3.570 | 4.073 | +0.503 | **+14.1%** | +7.8 pp |
| Recovery (GM) | 1.259 | 1.219 | −0.040 | −3.2% | −3.3 pp |

Monte Carlo uncertainty (20 independent seeds) yields *σ*_*θ*_ inflation of +22.1 ± 0.3% SE (95% CI: 21.5 to 22.7%), confirming the effect is precisely estimated and replicable.

Because *h*^2^ is a monotone function of *σ*_*θ*_ at the extrapolation point (Proposition B2, Section B), this upstream parameter distortion propagates directly to Shenhar’s headline estimate: the 50% “intrinsic” heritability is not purely intrinsic if the calibrated *σ*_*θ*_ includes absorbed extrinsic structure. The remaining sections trace the mechanism (§4.2–4.3), demonstrate independence from survival conditioning (§4.4), identify structural fingerprints (§4.5–4.7), and establish scope and falsifiability (§4.8–4.12).

### 4.2 Two-component bias decomposition

The bias decomposes into two competing forces operating in opposite directions. Under correct specification (Baseline: no extrinsic heterogeneity), the calibrate-then-extrapolate procedure slightly **underestimates** intrinsic heritability: observed *h*^2^ = 0.472 versus oracle *h*^2^ = 0.51, a gap of −3.7 pp (Figure 1). This attenuation reflects the fact that extrinsic mortality compresses twin correlations even when the model is correctly specified — an estimand shift, not a pipeline error. The Baseline shows the calibrate-then-extrapolate logic is internally consistent under correct specification.

**Figure 1:**
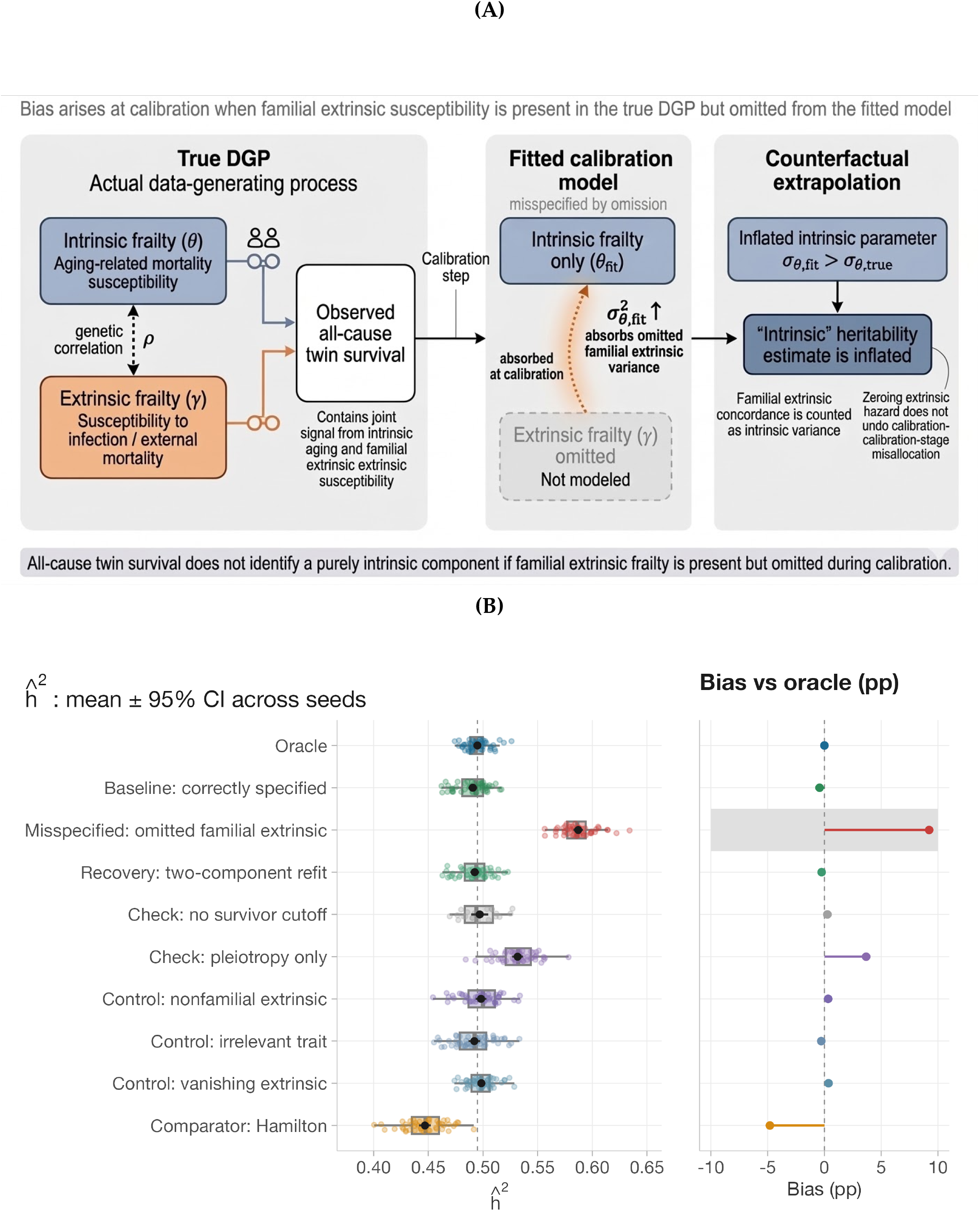
Mechanism and main results. **(A)** Schematic of the bias mechanism: when heritable extrinsic frailty (*γ*) exists in the true DGP but is omitted from the fitted model, the calibration step absorbs familial extrinsic concordance into the intrinsic parameter *σ*_*θ*_, inflating the extrapolated “intrinsic” heritability estimate. **(B)** Falconer *h*^2^ across all simulation conditions (Gompertz–Makeham, 50-seed means ± 95% CI; Check: no survivor cutoff uses 20 seeds). Dashed line: oracle intrinsic *h*^2^. Misspecified shows net upward bias of +9.2 pp. Correctly specified conditions (Baseline, Recovery) cluster near the oracle. Negative controls confirm specificity (§4.8); Check: no survivor cutoff shows bias persists without survival conditioning.

**Figure 2:**
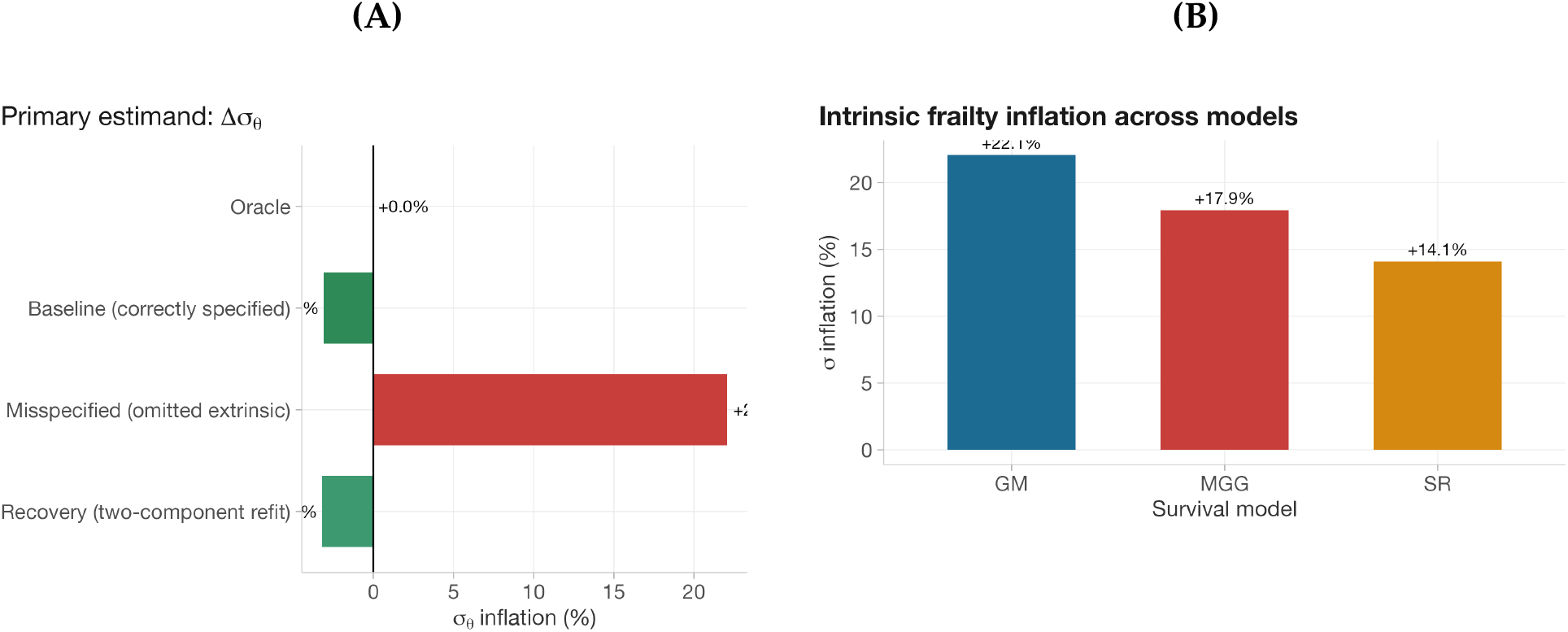
Primary estimand. **(A)** *σ*_*θ*_ inflation across simulation arms (GM model): the misspecified model inflates *σ*_*θ*_ by +22.1%; the recovery model restores it to within −3.2%. **(B)** Cross-model comparison: *σ*_*θ*_ inflation under misspecification is +22.1% (GM), +17.9% (MGG), +14.1% (SR) — the mechanism is model-general.

Under misspecification (Misspecified: heritable extrinsic susceptibility with *σ*_*γ*_ = 0.40, *ρ* = 0.4), the calibrated *σ*_*θ*_ inflates by 22.1% — from 1.2588 to 1.537 — producing *h*^2^ = 0.586, an inflation of +11.4 pp relative to the Baseline. This more than offsets the attenuation, yielding a net upward bias of +7.6 pp (50-seed mean: +9.2 pp, 95% CI: 8.7 to 9.7 pp). Throughout, we refer to the downward extrinsic-noise term as the **attenuation component** and the upward variance-absorption term as the **inflation component**; their sum is the **net bias**. The 11.4 pp inflation component is the methodological vulnerability.

The most decisive causal evidence comes from a recovery analysis. If the bias is genuinely caused by omitting *γ* from the model, then a **correctly specified** model — one that includes extrinsic frailty with its true parameters — should recover the true *σ*_*θ*_ and eliminate the inflation component. We test this by holding (*σ*_*γ*_, *ρ*) at their true DGP values and calibrating only *σ*_*θ*_ to match the observed *r*_MZ_ from the misspecified data. The recovered *σ*_*θ*_ = 1.219 (−3.2% from the true value), yielding *h*^2^ = 0.476 (−3.3 pp bias). The residual bias matches the Baseline’s attenuation (−3.7 pp): both are correctly specified models subject to the same estimand shift from extrinsic noise. Crucially, the **inflation component** is eliminated *σ*_*θ*_ is recovered to within −3.2% of its true value. This confirms that the net upward bias under misspecification is specifically attributable to the omitted extrinsic frailty component and is resolvable by correctly specifying the model.

#### Notation

All bias estimates are Monte Carlo averages over independent seeds: 50 seeds for Figure 1(B) headline estimates, 20 seeds per grid point for sensitivity sweeps, across all three mortality models (GM, MGG, SR). All 95% CIs throughout are *t*-based 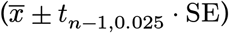. Table 5 reports single-seed values for the pedagogical decomposition (MASTER_SEED = 66829).

**Table 5.** Single-seed decomposition (Gompertz–Makeham, MASTER_SEED = 66829) showing the fitted intrinsic parameter *σ*_*θ*,fit_ alongside *h*^2^ and bias. The *σ*_*θ*,fit_ column traces the variance-absorption mechanism: misspecification inflates *σ*_*θ*,fit_ from 1.259 to 1.537, driving the net upward bias. Multi-seed distributional summaries appear in Figure 1(B).

| Condition | $\sigma_{\theta, \text{fit}}$ | $h^2$ | Bias (pp) |
| --- | --- | --- | --- |
| Oracle | 1.259 | 0.51 | — |
| Baseline: correctly specified | 1.22 | 0.472 | −3.7 |
| **Misspecified: omitted familial extrinsic ($\rho = 0.4$, $\sigma_\gamma = 0.40$)** | **1.537** | **0.586** | **+7.6** |
| Recovery: two-component refit | 1.219 | 0.476 | −3.3 |
| Check: no survivor cutoff | 1.286 | 0.497 | +0.3 |
| Check: pleiotropy only ($\rho = 0$) | 1.376 | 0.528 | +1.9 |
| Control: nonfamilial extrinsic | 1.259 | 0.478 | −3.1 |
| Control: irrelevant trait ($\zeta$) | 1.264 | 0.502 | −0.7 |
| Control: vanishing extrinsic ($m_{\text{ex}} = 0.0002$) | 1.267 | 0.5 | −0.9 |
| Comparator: Hamilton | N/A | 0.427 | −8.2 |

### 4.3 Variance absorption diagnostic

To confirm that the inflation reflects genuine variance absorption — rather than some other artifact — we compare the calibrated 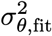against the prediction from the omitted-variable heuristic (Equation (3)) across all 81 cells of the *ρ* × *σ*_*γ*_ sensitivity grid. If the mechanism is variance absorption, then the fitted parameter should track the predicted value 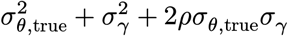.

It does. The *α* = *β* = 1 approximation yields a mean absolute residual of 8.6% (Figure 3). An OLS fit estimates *α* = 1.66, *β* = 1.25 with *R*^2^ = 0.988, confirming that the calibrated parameter is not arbitrarily wrong — it converges to a pseudo-true value in White’s sense (White, 1982) that systematically incorporates both the omitted extrinsic frailty variance and its covariance with intrinsic frailty. The empirical coefficients exceed unity (*α* > 1, *β* > 1) for three reasons. First, the first-order linearization underlying Equation (3) omits higher-order terms in the delta expansion; since both *θ* and *γ* enter through exponential terms (exp(*θ*) and exp(*σ*_*γ*_*γ*)), Jensen-type convexity effects inflate the effective variance absorbed by the calibration. Second, the calibration functional 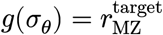 involves implicit age-aggregation: the MZ correlation is an average over the full lifespan distribution, and the sensitivity weights *κ*(*t*) (see Section B, §B.2) vary with age, creating a weighted average that differs from the local linearization. Third, lognormal mixing of extrinsic frailty produces heavier tails than a Gaussian approximation, further amplifying the empirical absorption coefficients. The heuristic (Equation (3)) provides the correct structural form (variance + cross-term), while the empirical *α, β* quantify the effective magnitude under the specific hazard structure and calibration criterion used here. Appendix B (Section B, Proposition B1) establishes the first-order direction of inflation under simplifying assumptions; Equation (3) summarizes a higher-order empirical approximation to the same pseudo-true mapping. Table 6 links the proof’s theoretical objects to the simulation measurements.

**Table 6.** Mapping between the formal proof’s theoretical objects and the simulation’s reported quantities. The proof establishes the sign and mechanism; the simulation quantifies the exact magnitude beyond the first-order approximation.

| Theoretical object (Appendix B) | Simulation measurement |
| --- | --- |
| $\hat{\sigma}_\theta$ from calibration Equation (25) | Fitted $\sigma_\theta$ returned by bisection matching $r_{\text{MZ}}$ |
| $\hat{\sigma}_\theta > \sigma_\theta$ (Proposition B1) | Inflation: $\sigma_{\text{infl}} = 100 \times (\sigma_{\text{fit}}/\sigma_{\text{true}} - 1) \%$ |
| $h^2(\hat{\sigma}_\theta) > h^2(\sigma_\theta)$ (Proposition B2) | Falconer $h^2$ at $m_{\text{ex}} = 0$: Misspecified minus Oracle |
| $\kappa_\theta, \kappa_\gamma$ (sensitivity coefficients) | Implicit in empirical $\alpha, \beta$ from Equation (3) regression |

**Figure 3:**
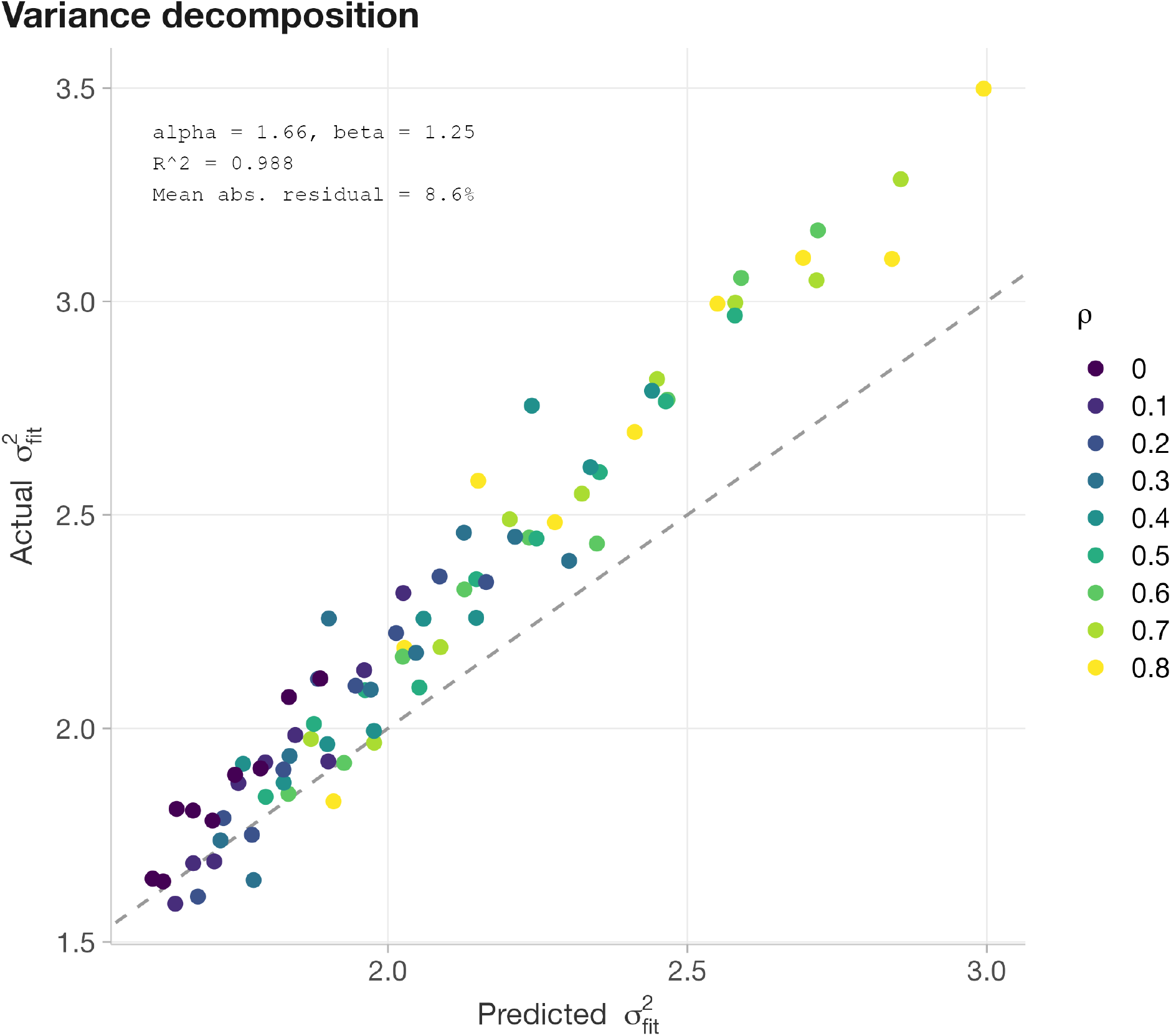
Pseudo-true variance prediction tracks fitted variance. Each point is one (*ρ, σ*_*γ*_) cell. Points cluster near the identity line (mean absolute residual = 8.6%). Empirical coefficients: *α* = 1.66, *β* = 1.25, *R*^2^ = 0.988.

The *R*^2^ = 0.988 predictive accuracy confirms that the calibrated *σ*_*θ*_ systematically incorporates omitted extrinsic structure — not random Monte Carlo error — undermining the interpretation of Shenhar’s fitted intrinsic parameter as purely intrinsic.

### 4.4 Bias without concordant-survivor restriction

A critical distinction from Hamilton’s collider bias is that our mechanism does not require removing any individuals from the analysis. When we repeat the full Misspecified analysis with cutoff = 0 — all twin pairs included regardless of age at death, including infant deaths — the misspecification bias is +0.3 pp (20-seed mean; 95% CI: −0.5 to 1 pp), reduced relative to the +9.2 pp observed with the standard age-15 cutoff; the point estimate remains positive, though the 95% CI spans zero, so the no-cutoff bias alone is not statistically distinguished from null at conventional levels. The key inferential point is that the variance-absorption mechanism does not require concordant-survivor restriction by construction — it operates at the calibration step regardless of whether the sample is restricted. The additional bias from the age-15 cutoff reflects genuine survivor-selection amplification: pairs where both twins survive past 15 are genetically non-representative when extrinsic mortality is heritable. Without such amplification, the net effect is smaller and estimated with lower precision, but the mechanism is structurally present the key distinction from Hamilton’s mechanism, which requires literal removal of individuals. This establishes that the bias mechanism threatening Shenhar’s estimate is analytically distinct from concordant-survivor conditioning and does not depend on sample restriction.

**Tier 2: Structural fingerprints** — Does the absorption leave observable traces beyond the scalar parameter?

### 4.5 Bivariate dependence diagnostics reveal structured misfit

A one-component calibration could in principle inflate a variance parameter yet still provide an adequate reduced-form approximation to the dependence structure. To test whether that is the case here, we compared age-specific bivariate dependence under the true two-component DGP, the misspecified one-component fit calibrated to match *r*_MZ_, and the correctly specified two-component fit. Diagnostics were computed on held-out simulations (≈ 44000 pairs per zygosity; Appendix, Section A.14) to avoid reusing calibration realizations. We report four complementary views: the 1D dependence ratio *R*(*t*), the full 2D residual surface Δ*R*(*t*_1_, *t*_2_), the bivariate survival surface misfit Δ*S*(*t*_1_, *t*_2_) with formal goodness-of-fit metrics, and the age-specific concordance curve *C*(*t*).

#### 1D dependence curve and 2D residual surface

We summarize dependence using *R*(*t*) = log{*P* (*T*_1_ > *t, T*_2_ > *t*)/*P* (*T* > *t*)^2^} and its full bivariate extension *R*(*t*_1_, *t*_2_). Despite matching the calibration target, the misspecified model does not recover the dependence structure (Figure 4). Instead, it systematically overpredicts late-life concordance and modestly underpredicts earlier concordance in both MZ and DZ twins, with peak deviation near age 81. The integrated absolute error for MZ twins is 0.0135 under the misspecified fit versus 0.0029 under the correctly specified two-component model. This directional residual pattern is exactly what the variance-absorption mechanism predicts: when familial extrinsic susceptibility is omitted, the fitted intrinsic parameter is inflated, shifting concordance toward ages where intrinsic hazard contributes most strongly.

**Figure 4:**
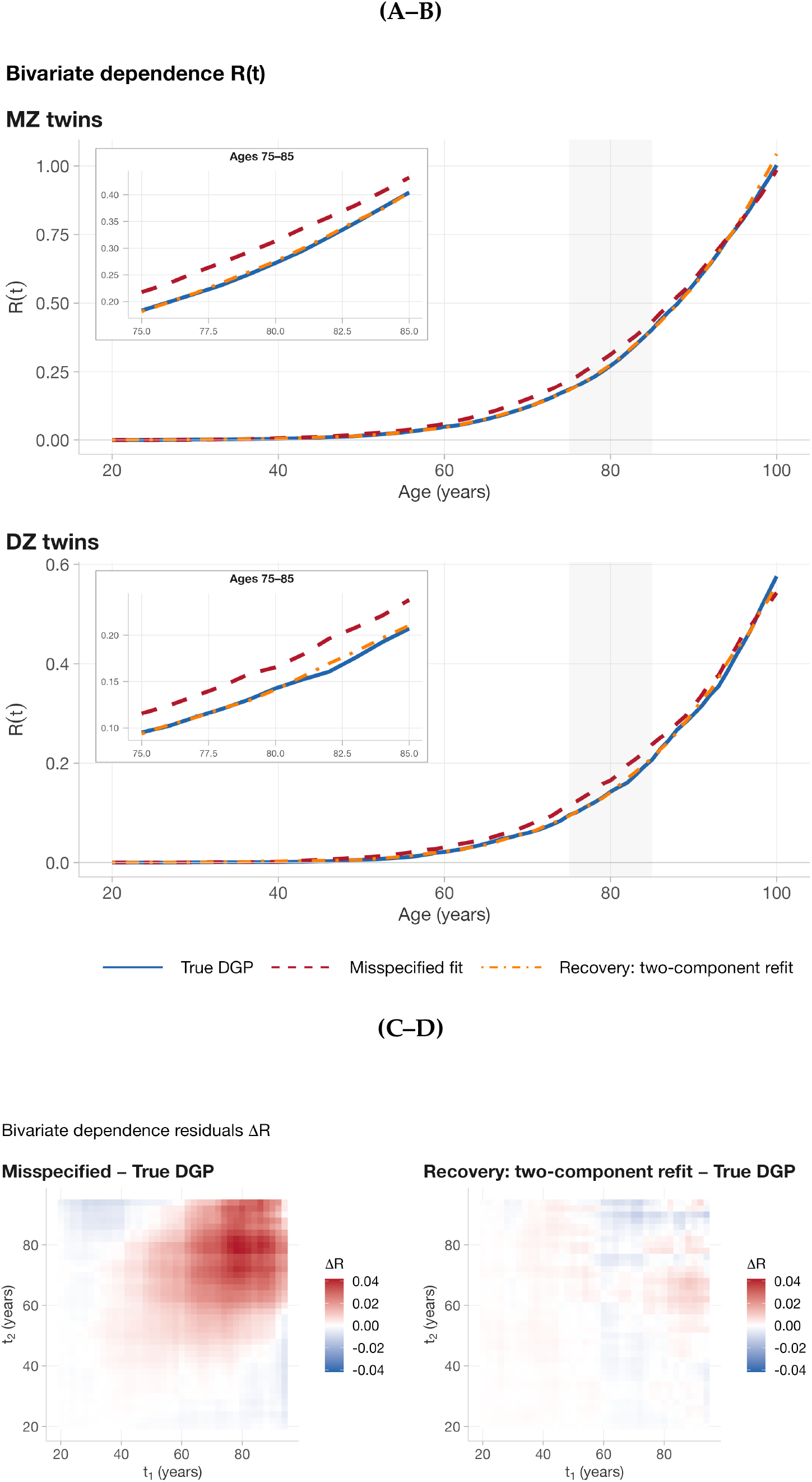
Bivariate dependence diagnostic. **(A–B)** Dependence ratio *R*(*t*) for MZ (top) and DZ (bottom) twins under three DGPs: true two-component model (blue, solid), misspecified one-component fit (red, dashed), and correctly specified recovery refit (orange, dot-dash). Despite matching *r*_MZ_ by construction, the misspecified fit overpredicts late-life concordance (peak deviation at age 81) and slightly underpredicts early-life concordance. IAE (MZ): misspecified = 0.0135, recovery = 0.0029 (4.6× reduction). **(C–D)** 2D bivariate surface residuals Δ*R*(*t*_1_, *t*_2_) for MZ twins. The misspecified fit (C) shows a warm region in the upper-right quadrant (both twins aged 60–90), consistent with excess late-life concordance from inflated *σ*_*θ*_; the recovery refit (D) eliminates this pattern.

The full residual surface shows the same pattern spatially (Figure 4, panels C–D). Relative to the true DGP, the misspecified fit produces a concentrated region of excess concordance in the upper-right quadrant of the (*t*_1_, *t*_2_) surface, corresponding to joint survival at older ages; this structure is largely absent under the correctly specified two-component model. Thus, the misspecification is not limited to scalar bias in *σ*_*θ*_: it leaves a recognizable age-structured footprint in the observable bivariate survival surface.

#### Bivariate survival surface misfit

The bivariate survival surface *S*(*t*_1_, *t*_2_) complements the dependence ratio by capturing both marginal and dependence effects in a single quantity. Figure 5 shows 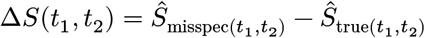 for MZ twins. The surface is predominantly negative: the mis-specified model underpredicts joint survival across young and middle ages because it lacks the extrinsic concordance channel that operates at all ages through the constant *m*_ex_ hazard term. A small positive region at extreme old ages (both twins > 85) confirms that the inflated *σ*_*θ*_ overcompensates through the exponentially growing Gompertz term. The overall pattern reflects concordance redistribution: the single-component model shifts predicted twin similarity from younger ages (where extrinsic frailty acts) toward older ages (where intrinsic aging dominates). This contrasts with the dependence-ratio residual Δ*R* (Figure 4, panels C–D), which isolates excess late-life dependence; Δ*S* additionally reflects marginal survival differences, producing the dominant negative region.

**Figure 5:**
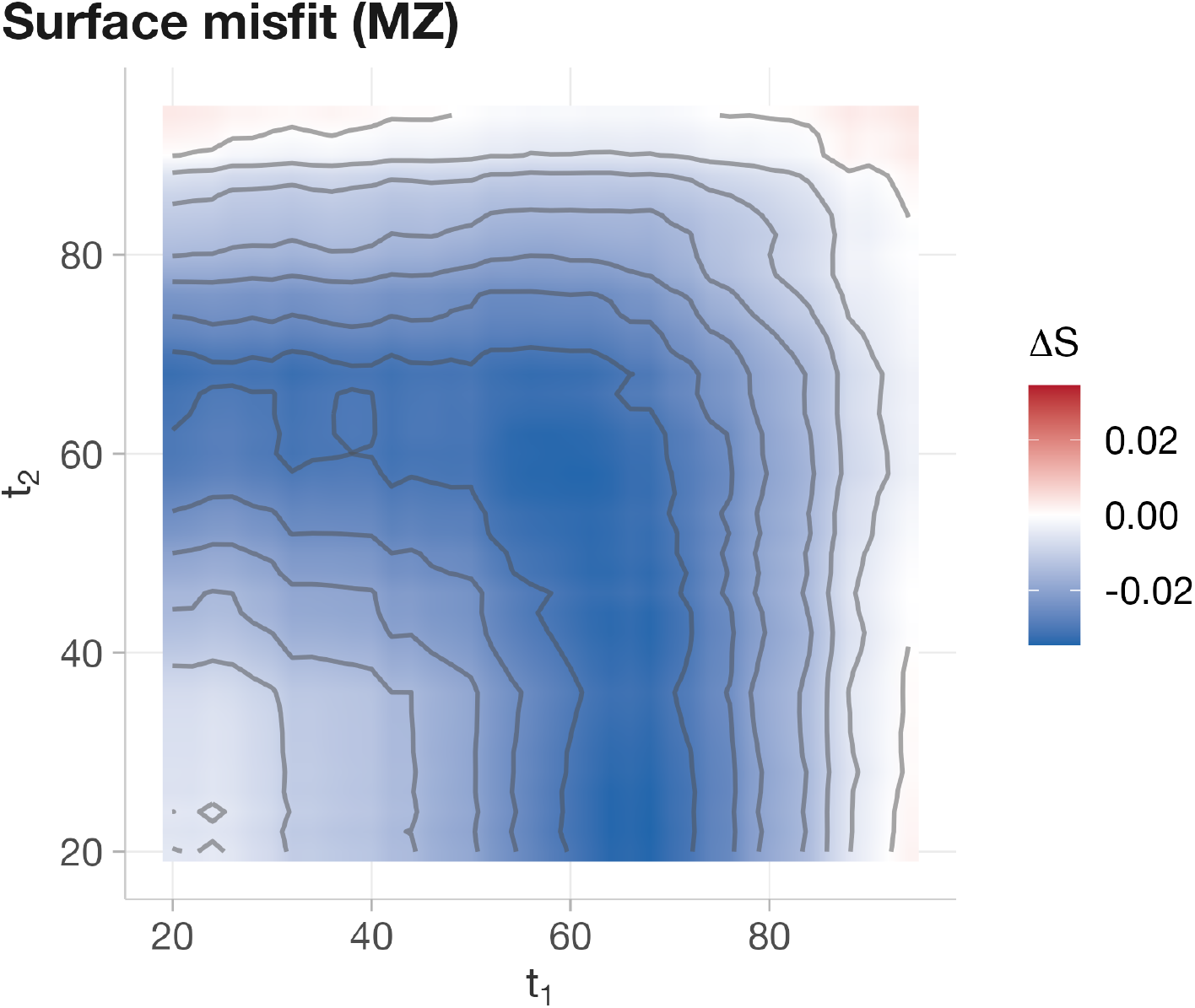
Bivariate survival surface misfit (MZ). Difference Δ*S* =*Ŝ*_misspec_ − *Ŝ*_true_ on the (*t*_1_, *t*_2_) grid. The predominantly negative (blue) surface indicates the misspecified model underpredicts joint survival at young and middle ages, where the missing extrinsic concordance channel contributes most. A faint positive region at extreme old ages (> 85) reflects overcompensation from inflated *σ*_*θ*_ through the Gompertz term — concordance is redistributed from younger to older ages. ISE ratio (misspecified/recovery) = 48.4×; sup|Δ*S*| = 0.035.

The integrated squared error (ISE) is 2.54 for the misspecified model versus 0.052 for the recovery model an ISE ratio of 48.4×. The supremum deviation is sup|Δ*S*| = 0.035, driven by the negative region at middle ages where the missing extrinsic channel contributes most. These formal surface-based metrics confirm that the misspecified model does not just shift a scalar summary — it fails to reproduce the entire joint survival structure by a factor of nearly 50.

#### Concordance curve

The concordance curve *C*(*t*) = ℙ(*T*_2_ > *t* | *T*_1_ > *t*) (Figure 6) shows the age pattern more directly. The misspecified model slightly underpredicts concordance at younger ages (where extrinsic frailty contributes most to twin similarity) and converges toward the true DGP at older ages where intrinsic Gompertz aging dominates. The recovery model tracks the truth closely at all ages. Standard errors are computed via pair bootstrap (200 replicates resampling twin pairs).

**Figure 6:**
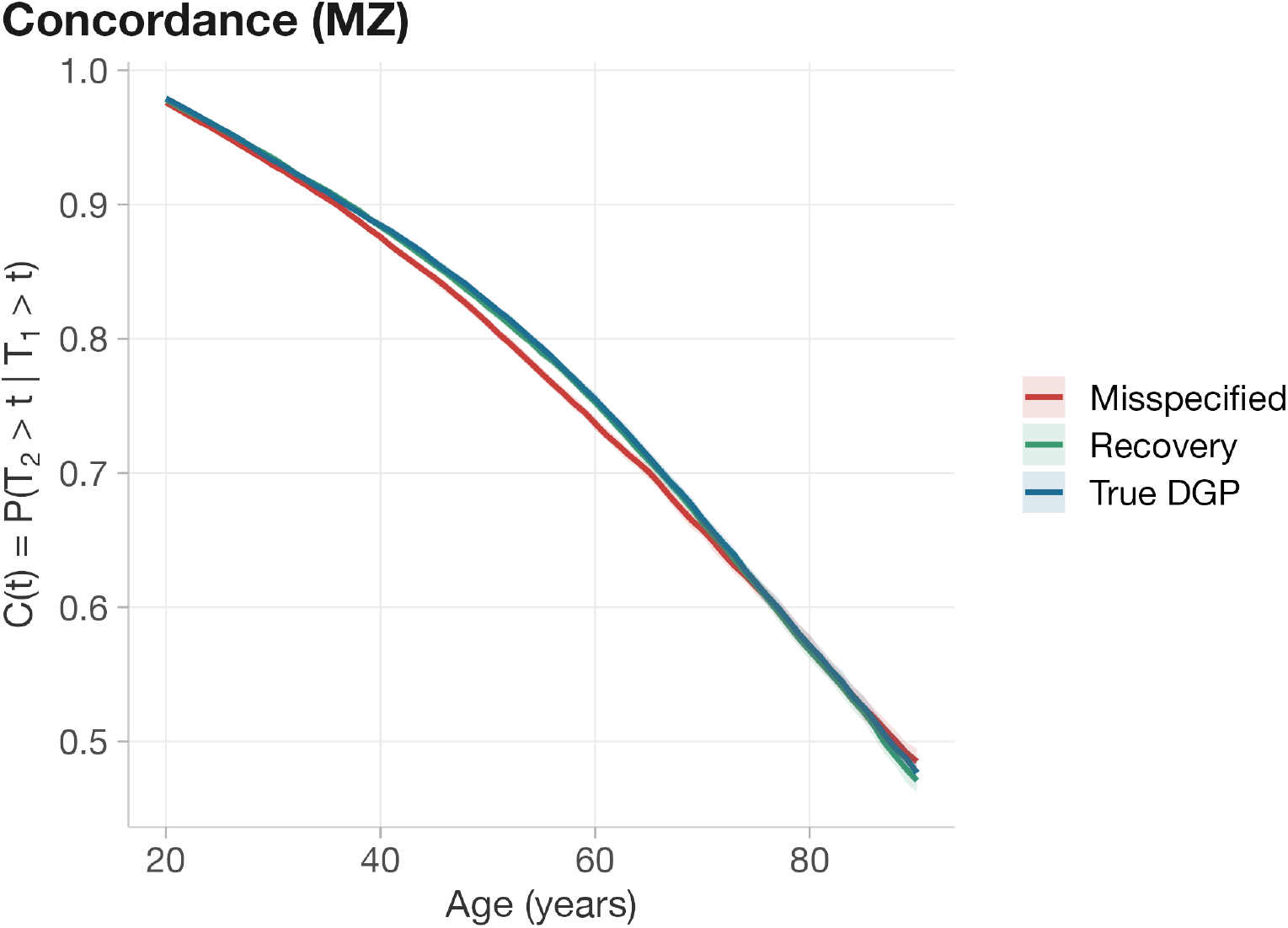
Age-specific concordance *C*(*t*) = ℙ(*T*_2_ > *t* | *T*_1_ > *t*) for MZ twins under three DGPs. Shaded bands: 95% CI from pair bootstrap (200 replicates). The misspecified model underpredicts concordance at younger ages; the recovery model closely matches the oracle.

These structural fingerprints would be detectable in Shenhar’s cohort data via posterior predictive checks on MZ/DZ bivariate survival surfaces: if the misspecified model produces the characteristic early-deficit/late-excess residual pattern predicted by the variance-absorption mechanism, the single-component assumption is empirically falsified.

### 4.6 Age-specific conditional correlation

The conditional Pearson correlation cor(*T*_1_, *T*_2_ | *T*_1_ > *t, T*_2_ > *t*) provides an intuitive summary of how the misspecified model distorts twin dependence across the age range (Figure 7). Under the misspecified fit, MZ twins show excess conditional correlation at all ages from 20 to 90, with a maximum discrepancy of Δ*r* = 0.048 at age 80 (Figure 7, panel A). For DZ twins, the pattern is similar but smaller in magnitude (Δ*r* = 0.045 at age 90 panel B). The recovery model closely matches the oracle for both zygosities.

**Figure 7:**
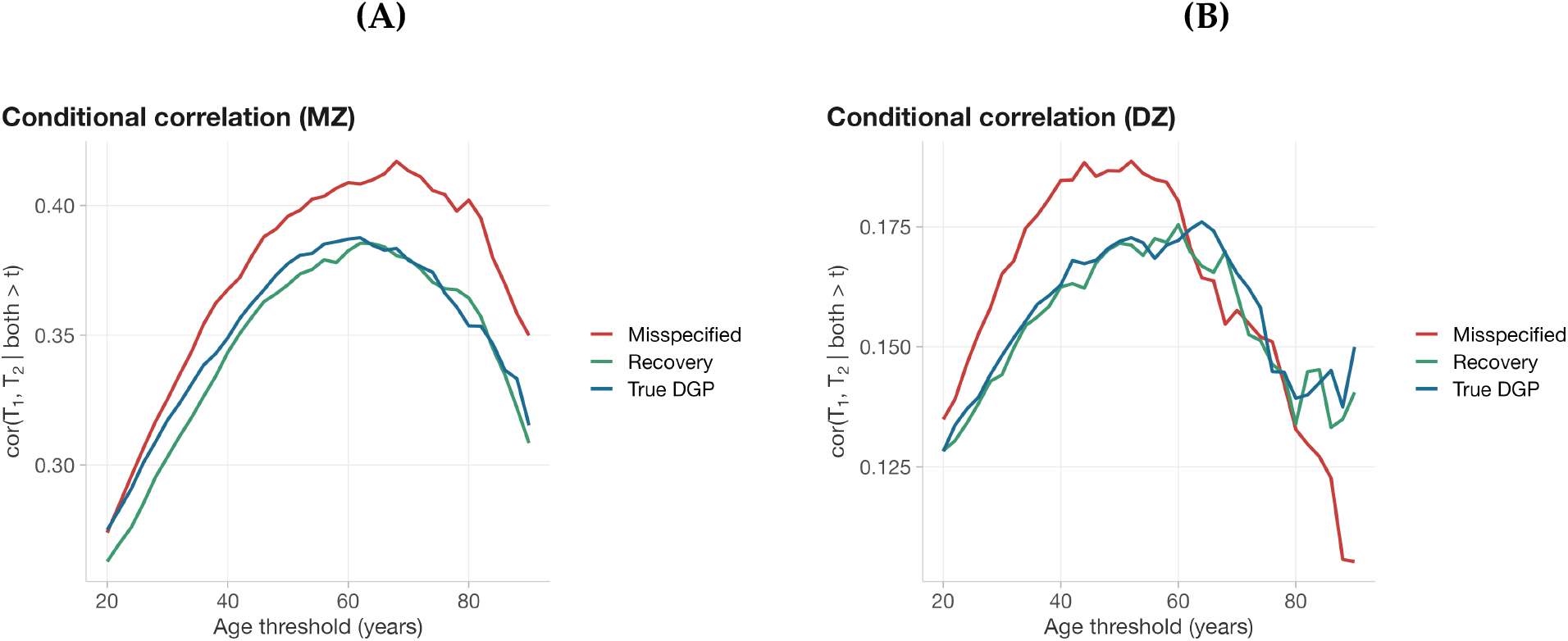
Age-specific conditional correlation. **(A)** MZ twins: cor(*T*_1_, *T*_2_ | *T*_1_ > *t, T*_2_ > *t*) as a function of age threshold *t*. The misspecified model (red) shows excess dependence at all ages, peaking at age 80 (Δ*r* = 0.048). (B) DZ twins: same pattern with smaller magnitude (Δ*r* = 0.045 at age 90).

This is the expected structural signature of *σ*_*θ*_ inflation. Intuitively, inflating *σ*_*θ*_ increases the amount of shared latent heterogeneity among surviving pairs, which raises conditional dependence at each conditioning age *t*. By forcing all twin concordance through the intrinsic aging parameter, the misspecified model produces systematically higher conditional correlations than the true DGP, where concordance arises from both intrinsic **and** extrinsic sources. The mean absolute discrepancy across ages is 0.022 for MZ and 0.014 for DZ twins. If Shenhar’s estimate were free of absorption bias, the conditional correlation pattern under their fitted model should match the observed age-dependence structure; the systematic excess predicted here is a testable diagnostic.

### 4.7 ACE twin decomposition: switching summary statistics does not restore identification

A natural objection is that the bias may be specific to Falconer’s formula *h*^2^ = 2(*r*_MZ_ − *r*_DZ_) rather than reflecting a genuine identification failure. To test this, we fitted classical ACE structural equation models via OpenMx to rank-normalized simulated lifespans under each DGP (Figure 8). ACE decomposes twin lifespan variance into A (additive genetic), C (shared environment), and E (unique environment), with the expected covariance structure: Cov_MZ_ = *A* + *C*, Cov_DZ_ = 0.5*A* + *C*, Var = *A* + *C* + *E*. Under the oracle (*m*_ex_ = 0), ACE correctly recovers *A* = 0.5 × 100%, *C* ≈ 0%, *E* = 0.5 × 100% — matching the DGP’s *h*^2^ = 0.50 as expected. Under the true DGP with heritable extrinsic frailty (*σ*_*γ*_ = 0.40, *ρ* = 0.4, *m*_ex_ = 0.004), the ACE-estimated *A* drops to 0.3 × 100%, reflecting the dilution of genetic signal by extrinsic mortality — but crucially, *C* ≈ 0. The heritable extrinsic frailty does **not** manifest as a “shared environment” component in ACE. Instead, because MZ twins share *γ* identically and DZ twins share it at 0.5 — the same pattern as additive genetics — the extrinsic concordance is absorbed into *A* rather than *C*.

**Figure 8:**
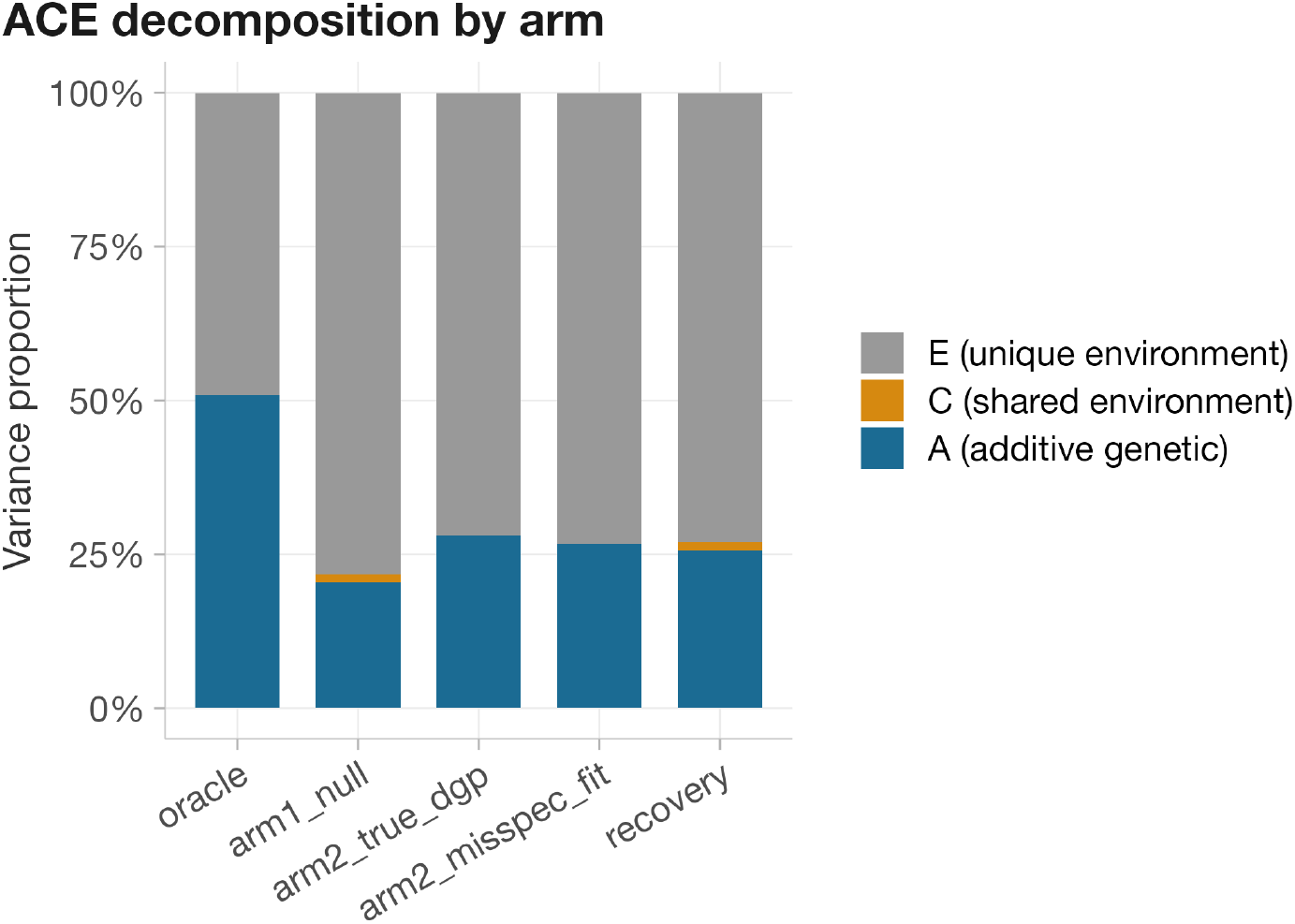
ACE decomposition (rank-normalized lifespans, OpenMx). A = additive genetic, C = shared environment, E = unique environment. Under all conditions, *C* ≈ 0 — the heritable extrinsic frailty is absorbed into *A* (genetic), not *C* (shared environment). ACE is equally vulnerable to the variance absorption mechanism; switching summary statistics does not resolve the identification failure.

This means ACE is **vulnerable in the same way under this omission pattern**: switching from Falconer to a structural equation model does not resolve the identification failure. This does not imply ACE is generally invalid; rather, when the omitted extrinsic component induces the same 1.0/0.5 sharing pattern as additive genetics, ACE cannot distinguish it from *A* without additional information. The bias is in the calibration step, not in the choice of heritability summary statistic — and Shenhar’s 50% estimate would not be rescued by adopting a structural equation model framework instead of Falconer’s formula.

To be precise about the epistemic role of this analysis: the ACE decomposition does not *establish* the calibration-absorption mechanism — that is the content of §4.1–4.3 and Proposition B1 (Section B). Rather, ACE serves as a *sensitivity and translation layer*. It shows that the identification failure persists when the summary vocabulary is changed from Falconer’s formula to structural equation modeling i.e., that the bias is not an artifact of any particular heritability summary but reflects the upstream inflation of 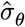, which propagates to any monotone downstream summary (Corollary B3). The cause is variance absorption at calibration; ACE confirms that its consequences are summary-independent.

**Tier 3: Scope, specificity, and falsifiability** — How general is the mechanism, what are its necessary conditions, and can it be falsified?

### 4.8 Specificity: negative controls

In our DGP, inflation arises from **heritable** extrinsic susceptibility that affects mortality; pleiotropy (*ρ* > 0) amplifies the effect, but is not required. Additional latent genetic structure alone is not sufficient. Three negative controls and a boundary check establish this.

When extrinsic susceptibility exists but is independent between twins (nonfamilial extrinsic: *γ* not shared), *h*^2^ = 0.478 (−3.1 pp), matching the Baseline: non-genetic noise adds individual variance but not twin concordance. When a heritable trait *ζ* is genetically correlated with *θ* at the same level as *γ* in the Misspecified condition but **does not enter the hazard function**, *h*^2^ = 0.502 (−0.7 pp, within Monte Carlo noise): the mechanism requires a mortality-relevant pathway. And when heritable *γ* is present but *m*_ex_ is negligible (vanishing extrinsic: *m*_ex_ = 0.0002, one-twentieth of historical levels), *h*^2^ = 0.5 (−0.9 pp): the extrinsic channel must carry non-negligible signal.

The inflation component is therefore specific: it requires (a) genetic concordance in extrinsic susceptibility, ^1^ (b) a mortality-relevant pathway, and (c) sufficient extrinsic hazard magnitude. Removing any one of these conditions eliminates the inflation. In the empirically relevant regime (*σ*_*γ*_ = 0.40), pleiotropy (*ρ* > 0) amplifies the effect substantially: heritable extrinsic susceptibility without pleiotropy (*ρ* = 0) changes the calibrated *σ*_*θ*_ by only 9.3%. Notably, even at *ρ* = 0 the net bias is still positive (+1.9 pp): heritable extrinsic frailty alone — without any shared genetic architecture with intrinsic aging produces a small but non-zero inflation because the calibration absorbs extrinsic concordance that the model cannot represent.

The recovery analysis (two-component refit, §4.2) complements these negative controls: a correctly specified two-component model recovers *σ*_*θ*_ to within −3.2% of its true value, eliminating the inflation component while preserving the same attenuation as the Baseline. Together, the negative controls (which rule out alternative explanations) and the recovery analysis (which confirms recoverability) establish that the bias threatening Shenhar’s estimate is specifically attributable to the omitted extrinsic frailty component.

### 4.9 Model and distributional generality

The mechanism is not specific to the Gompertz-Makeham functional form. We reimplemented Shenhar’s two other mortality models — the Saturating-Removal (SR) model and the Makeham-Gamma-Gompertz (MGG) model — using their published parameters, and ran Baseline and Misspecified under each. The SR model simulates a stochastic damage trajectory 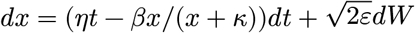, with death at a genetically variable threshold *X*_*c*_ ∼ 𝒩(17, 3.57^2^) and extrinsic mortality as an independent competing risk (Euler-Maruyama at weekly resolution; 50,000 pairs per zygosity). The MGG model uses Strehler-Mildvan coupling: *b*_*i*_ = *q*_*i*_*b*_0_, log *a*_*i*_ = *q*_*i*_ log *a*_0_ (*a*_0_ = 10^−5^, *b*_0_ = 0.115, *q* ∼ 𝒩(1, 0.27^2^)). Under the correctly specified null, both models approximately reproduce Shenhar’s published twin correlations and heritability estimates (Baseline bias < 1 pp for all three). ^2^

All GM estimates are averaged over 50 independent seeds, yielding +9.2 pp (95% CI: 8.7 to 9.7 pp) for the Misspecified condition and −0.4 pp (95% CI: −0.9 to 0.1 pp) for the Baseline — the latter indistinguishable from zero, confirming that the misspecified model, not the pipeline, drives the bias. The same analysis across all three model families at 20 seeds each (Figure 9, Table 7; Appendix, Section A.10) gives mean ± SE entirely positive and mutually overlapping: GM +9.2 ± 0.3 pp, MGG +10.4 ± 0.5 pp, SR +9.2 ± 0.5 pp, with all 95% CIs above +8 pp. The vulnerability is structural: none of the three models attach an individual subscript to *m*_ex_. Late-age mortality deceleration (SR’s saturating removal, MGG’s demographic selection) changes **how much** extrinsic frailty is absorbed but cannot eliminate absorption, because no model has a parameter to represent it. The MGG oracle *h*^2^ (0.464) is lower than GM’s (0.51) because Strehler-Mildvan compensation constrains the spread of intrinsic lifespans. The bias magnitude is statistically indistinguishable across model families (+9–10 pp), indicating that this is a general property of calibrate-then-extrapolate estimation under omitted extrinsic heterogeneity.

**Table 7.** Model-generality validation (20 independent seeds per model; 50,000 pairs per zygosity). Bias = Falconer *h*^2^ minus oracle *h*^2^, in percentage points; values are mean (95% CI). CIs are *t*-based 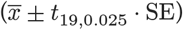. The misspecification bias is robustly positive (+9–10 pp) across all three mortality frameworks, with all 95% CIs above +8 pp. Baseline bias is indistinguishable from zero for all three models. See Figure 9 for the corresponding forest plot.

| Mortality framework | Oracle $h^2$ | Baseline bias (pp) | Misspec. bias (pp) |
| --- | --- | --- | --- |
| Gompertz–Makeham (ref.) | 0.492 | −0.1 (−0.9, +0.7) | **+9.2 (+8.5, +9.9)** |
| MGG (Shenhar) | 0.465 | +0.9 (+0.1, +1.7) | **+10.4 (+9.5, +11.3)** |
| SR (Shenhar) | 0.575 | +0.2 (−0.8, +1.2) | **+9.2 (+8.2, +10.2)** |

**Table 8.** Multi-target vs *r*_MZ_-only calibration under the Misspecified DGP. Both methods produce upward-biased 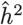; multi-target calibration attenuates but does not eliminate the net upward bias. The *r*_MZ_-only baseline here uses a separate simulation draw from the main analysis (Table 3) to ensure an internally consistent comparison; the 2 pp difference in bias (+10 vs +7.6 pp) reflects Monte Carlo variability across draws. Oracle *h*^2^ ≈ 0.51.

| Calibration criterion | $\sigma_\theta$ fit | Inflation | $\hat{h}^2$ at $m_{\text{ex}} = 0$ | Bias (pp) |
| --- | --- | --- | --- | --- |
| Multi-target (5 statistics) | 1.47 | +16.8% | 0.573 | **+6.4 pp** |
| $r_{\text{MZ}}$-only | 1.582 | +25.7% | 0.609 | **+10 pp** |

**Figure 9:**
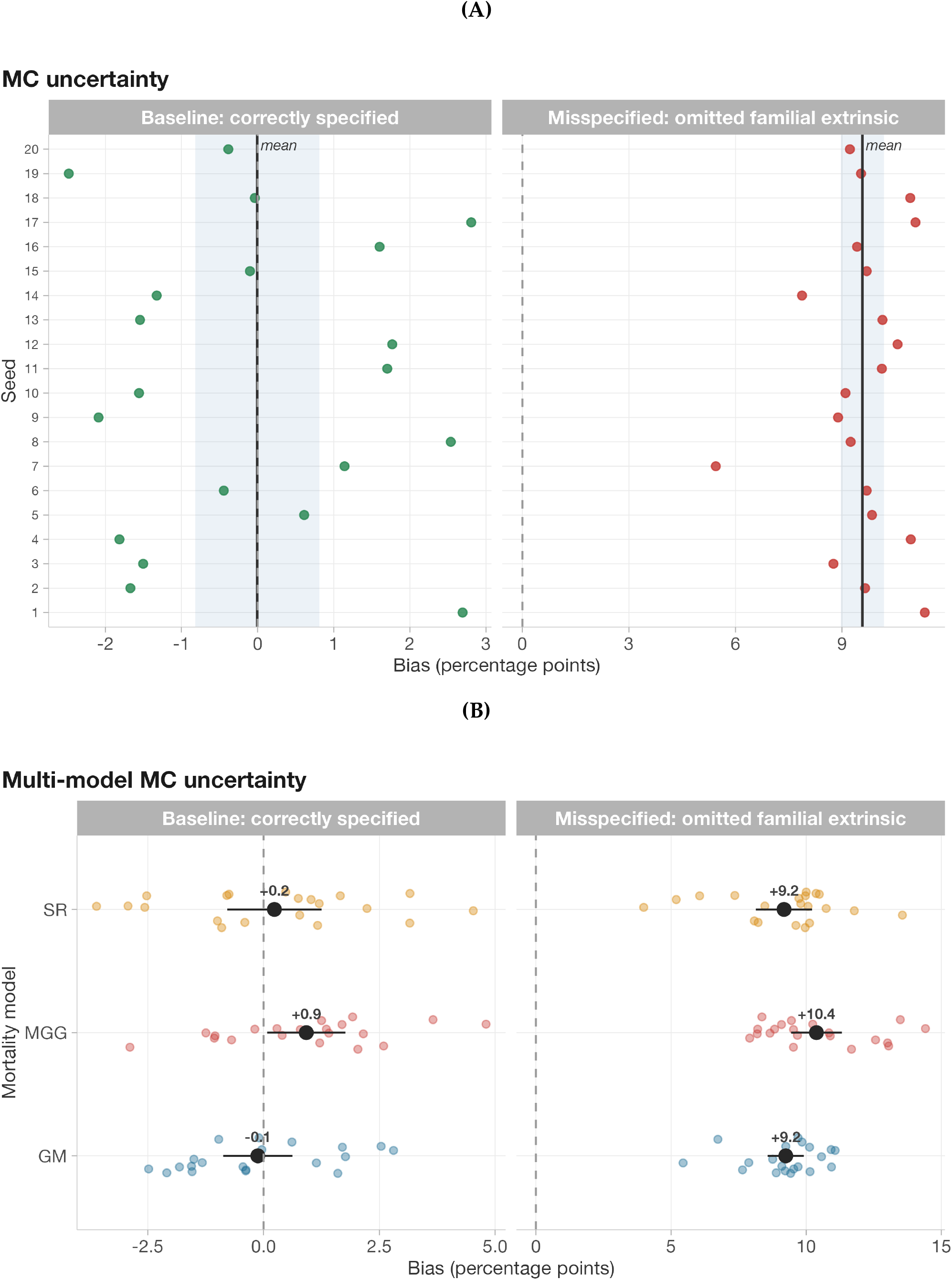
Monte Carlo uncertainty and model generality. **(A)** Per-seed bias estimates for the reference Gompertz–Makeham model (20 independent seeds). Left: Baseline (correctly specified) scatters symmetrically around zero. Right: Misspecified (omitted familial extrinsic) is consistently positive across all seeds, with mean +9.6 pp (95% CI: 9 to 10.2 pp). **(B)** The same analysis across all three mortality model families (20 seeds per model). Black point-range: mean ± 95% CI; colored dots: individual seeds. All three models yield consistent upward bias (+9–10 pp) with 95% CIs entirely above +8 pp. The vulnerability is structural and model-general.

The bias is also robust to the extrinsic frailty construction itself. The default DGP uses multiplicative log-normal frailty; we tested two alternatives — additive perturbation (*c*_*i*_ = *m*_ex_ + *σ*_*γ*_*γ*_*i*_) and gamma-distributed frailty — holding mean extrinsic hazard fixed at *m*_ex_. Across 10 seeds per construction, the bias is +8.1 ± 0.4 pp (log-normal), +8.7 ± 0.4 pp (additive), and +7.6 ± 0.6 pp (gamma), with all 95% CIs overlapping (Figure 17; Appendix, Section A.12). The 1 pp spread across constructions, compared to the ≈ +8 pp mean effect, confirms that the mechanism does not depend on the log-normal parameterization.

Since Shenhar use MGG and SR models in addition to Gompertz–Makeham, the presence of the bias in all three model families establishes that their published estimates are vulnerable regardless of which mortality model is adopted.

### 4.10 Dose-response to extrinsic mortality

Across our sweep, bias increases with the magnitude of extrinsic mortality (Figure 10). It is indistinguishable from zero below *m*_ex_ ≈ 0.001, rises to +7.9 pp at historical levels (*m*_ex_ = 0.004; slightly below the main Misspecified estimate of +7.6 pp because the sweep uses an independent simulation draw), and reaches +30.6 pp at *m*_ex_ = 0.012 (3× historical). This is the expected dose-response: extrinsic hazard must contribute materially to the total hazard over adult ages for genetically-patterned susceptibility to generate appreciable twin concordance. It also implies an empirical prediction: **the net bias should be larger in cohorts where infection mortality constitutes a larger share of total deaths** — directly testable across Shenhar’s Danish cohorts born 1870–1900.

**Figure 10:**
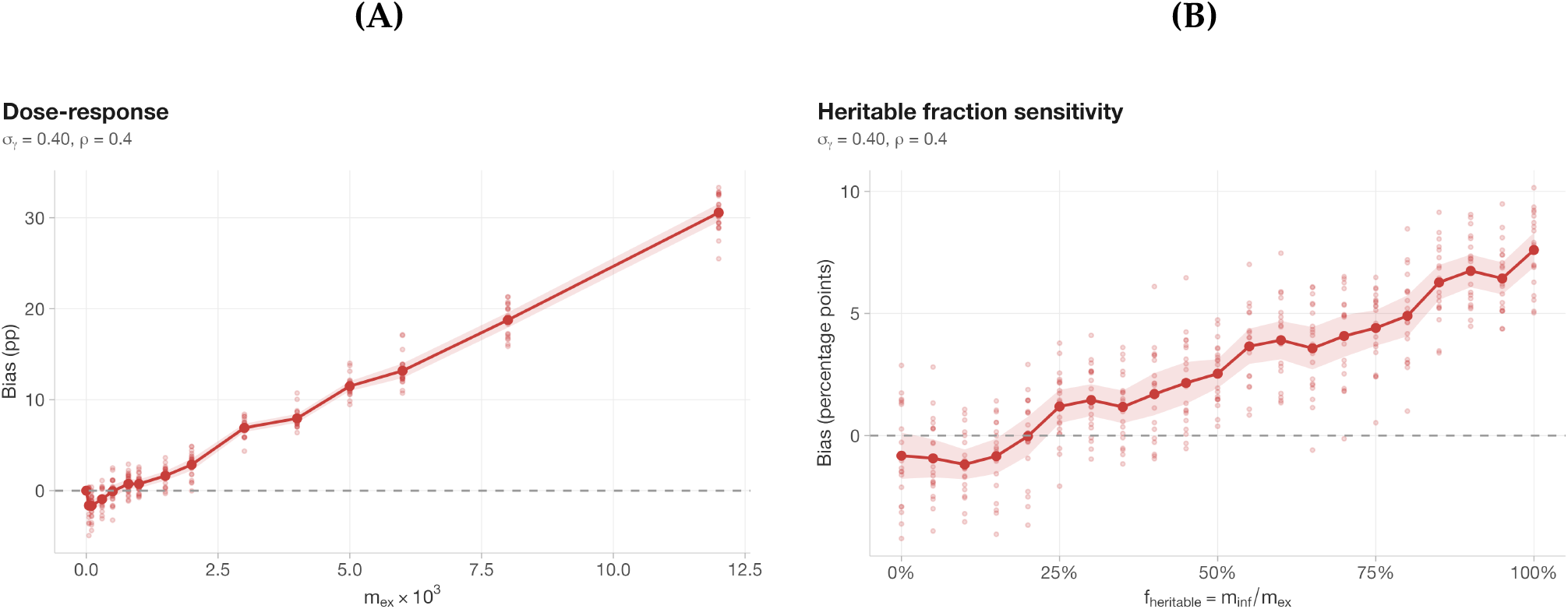
Dose-response structure. **(A)** Bias vs extrinsic mortality magnitude *m*_ex_ (20 seeds per level; ribbon = 95% CI; faint dots = individual seeds): bias rises from negligible at *m*_ex_ < 0.001 to +30.6 pp at 3× historical levels. **(B)** Bias vs heritable fraction of extrinsic mortality (*m*_ex_ = 0.004; 20 seeds per level; ribbon = 95% CI; faint dots = individual seeds). Bias increases monotonically with the heritable share. Together, the two panels show that bias requires both material extrinsic hazard magnitude and a heritable component.

A natural objection is that not all extrinsic mortality is heritable — accidents, war, and famine are largely non-genetic. We test this by decomposing *m*_ex_ = *m*_inf_ + *m*_other_, where only the infectious fraction *m*_inf_ receives genetic modulation while *m*_other_ remains a population constant. Figure 10 shows that bias increases monotonically with the heritable share (10 seeds per level): at 0% heritable, bias is −0.8 pp (indistinguishable from zero); at 50%, 2.5 pp; and at 100%, 7.6 pp. The monotonic relationship implies that if even a moderate minority of historical extrinsic deaths were influenced by heritable frailty, the distortion is positive (Appendix, Section A.9).

### 4.11 Anchored quantification

Within the externally informed restricted sensitivity regime (*σ*_*γ*_ ∈ [0.30, 0.65], *ρ* ∈ [0.20, 0.50]), the bias ranges from +2.8 to +18.9 pp, with a mean of 9 pp (Figure 11). At the upper-bound point estimate (*σ*_*γ*_ = 0.65, *ρ* = 0.35), the bias is +15.7 pp. Across the full exploratory sweep, the bias reaches +18.4 pp at *ρ* = 0.8, *σ*_*γ*_ = 0.55 (*h*^2^ = 0.69).

**Figure 11:**
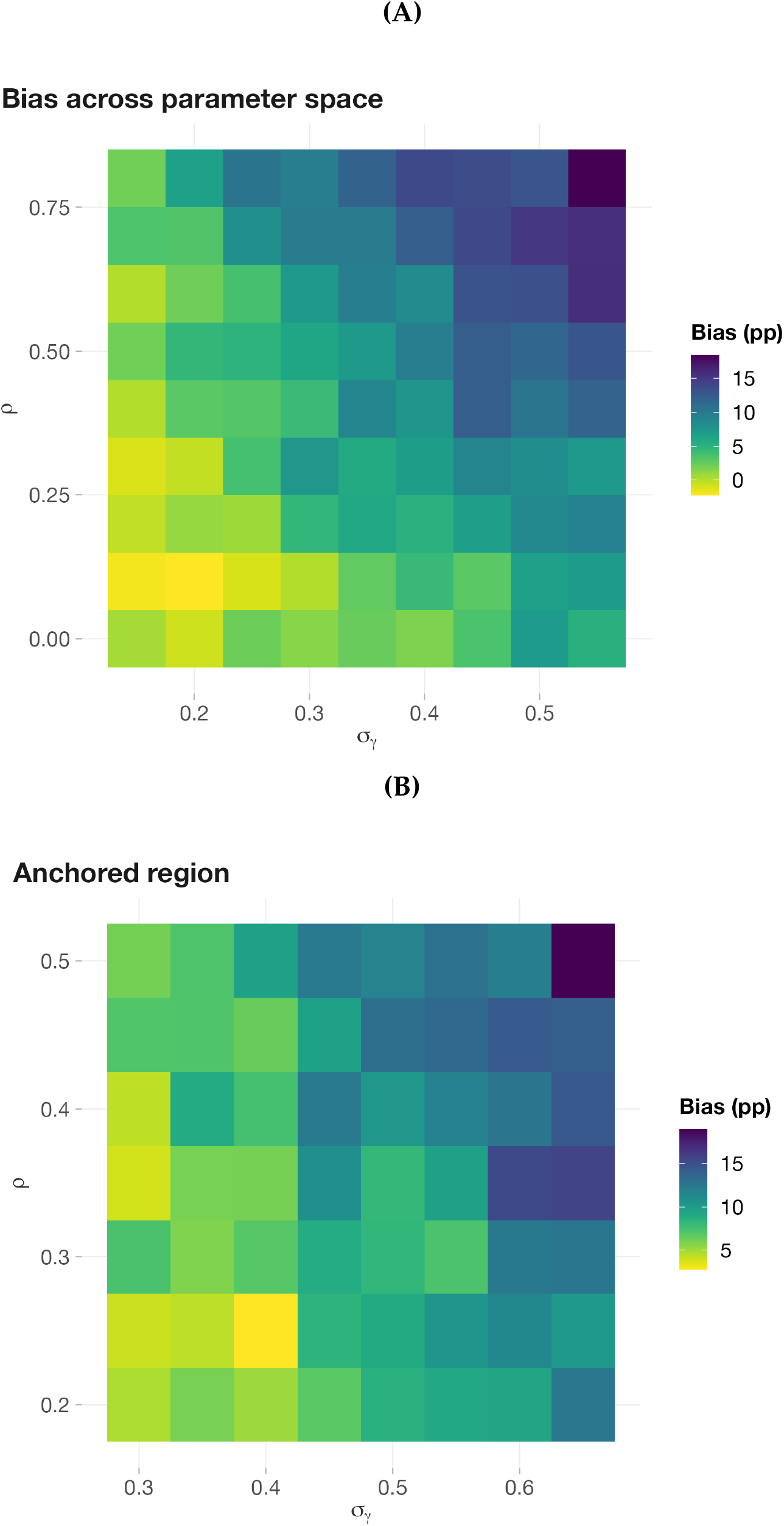
Sensitivity sweeps. **(A)** Full exploratory sweep: bias increases with both *ρ* and *σ*_*γ*_, reaching +18.4 pp at *ρ* = 0.8, *σ*_*γ*_ = 0.55. **(B)** Anchored regime (*σ*_*γ*_ ∈ [0.30, 0.65], *ρ* ∈ [0.20, 0.50]): bias ranges from +2.8 to +18.9 pp (mean 9 pp); point estimate (*σ*_*γ*_ = 0.65, *ρ* = 0.35): +15.7 pp.

**Figure 12:**
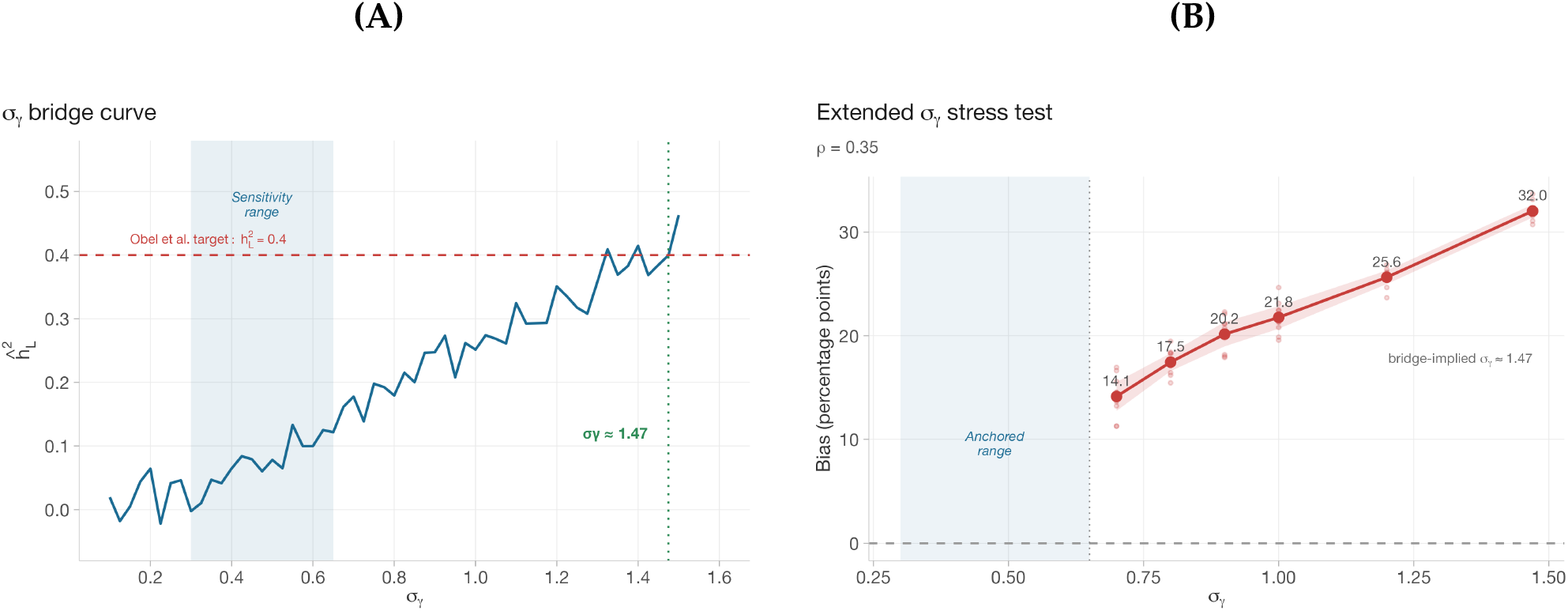
Conservative anchoring. **(A)** Bridging *σ*_*γ*_ to empirical infection-death heritability: matching Obel et al.‘s 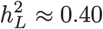 (dashed red) requires *σ*_*γ*_ ≈ 1.47 (dotted green) — more than double the sensitivity ceiling of 0.65 (shaded band). **(B)** Stress test beyond the anchored ceiling (*ρ* = 0.35; 10 seeds per level; ribbon = 95% CI). Bias continues to increase monotonically, reaching +32 pp at the bridge-implied *σ*_*γ*_. Together, the panels confirm that the anchored ceiling (*σ*_*γ*_ = 0.65) sits well below empirically implied levels, and that bias continues to increase beyond it.

These results do not resolve the true intrinsic heritability, but they establish that Shenhar’s estimate is biased upward by a quantitatively meaningful amount — sufficient to materially reduce their 50% estimate (by +15.7 pp at the upper-bound point estimate, and by 9 pp on average across the anchored regime). The analysis yields two falsifiable predictions: (1) a two-component reanalysis that separately models extrinsic frailty should reduce intrinsic *h*^2^ by a comparable amount; and (2) the net bias should be larger in cohorts with higher infection-mortality share. A third falsifiable prediction — sign reversal under negative genetic correlation — is confirmed in §4.12.

As a stress test, we extended the sensitivity sweep beyond the anchored ceiling to *σ*_*γ*_ = 1.47, the value implied by the tetrachoric bridge to liability-scale infection-mortality heritability (Figure 12, panel B; Appendix, Section A.11). Bias continues to increase monotonically, reaching +32 ± 0.6 pp (106.7% inflation of *σ*_*θ*_) at the bridge-implied value. While such extreme hazard heterogeneity may exceed what is biologically plausible under the stylized DGP, this confirms that bias increases monotonically beyond the anchored ceiling: if the true extrinsic heterogeneity lies closer to the liability-scale implication, the bias is strictly larger than the anchored range reports.

### 4.12 Sign reversal: a falsifiable prediction

The bias mechanism makes a strong, falsifiable prediction: if the genetic correlation between intrinsic aging and extrinsic susceptibility is negative (*ρ* < 0), the omitted extrinsic component should **deflate** rather than inflate the calibrated *σ*_*θ*_ — reversing the direction of bias. This follows from Corollary B4 in Section B: negative pleiotropy causes the omitted extrinsic concordance to oppose the intrinsic concordance, pulling the calibrated *σ*_*θ*_ below its true value.

Figure 13 confirms this prediction across all three mortality model families. Bias transitions smoothly from negative (downward) to positive (upward) as *ρ* crosses zero, with the sign reversal occurring near *ρ* = 0 in all three models. The monotone relationship between *ρ* and bias direction shows that the genetic correlation between immune function and intrinsic aging is not just a quantitative modifier — it determines whether Shenhar’s estimate is biased upward (inflated, if *ρ* > 0) or downward (deflated, if *ρ* < 0).

**Figure 13:**
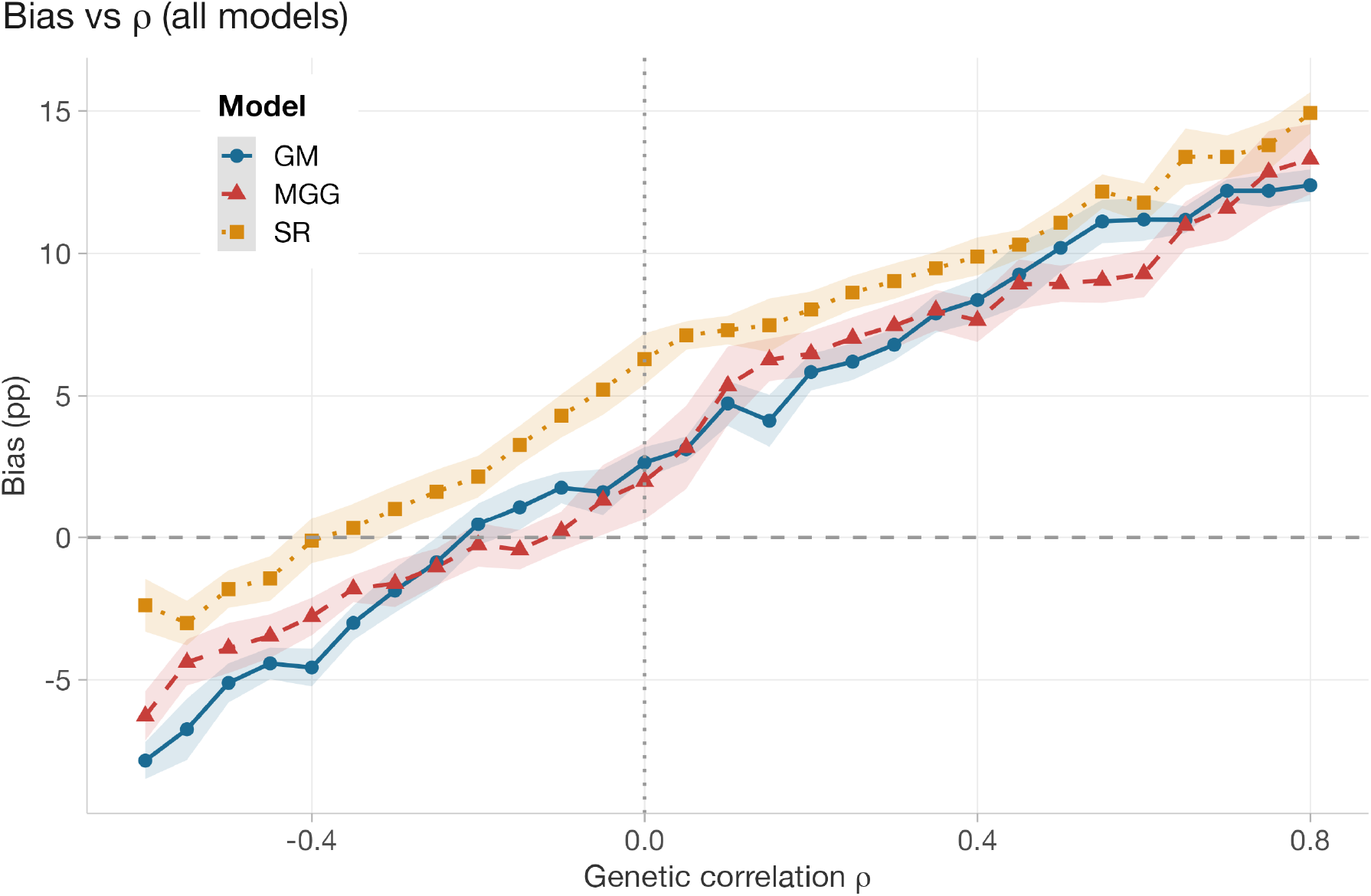
Sign reversal across all three mortality models. Bias vs genetic correlation *ρ* at *σ*_*γ*_ = 0.40 (20 seeds per grid point; ribbon = 95% CI). Negative *ρ* reverses the net bias sign — a falsifiable prediction confirmed in GM, MGG, and SR. The monotone relationship shows that the direction of the genetic correlation determines whether the omitted extrinsic frailty inflates or deflates the intrinsic estimate. The empirically motivated regime (*ρ* > 0, shaded) corresponds to upward bias.

For this prediction to refute our mechanism, independent evidence would need to establish *ρ* < 0 — i.e., that individuals genetically predisposed to extrinsic mortality are genetically protected against intrinsic aging, or vice versa. Current evidence points in the opposite direction: genetic correlations between COVID-19 severity and longevity are negative (*r*_*g*_ ≈ −0.35; (Qiu, 2025)), implying *ρ* > 0 in our notation (§3.4). The sign reversal therefore serves as an internal consistency check: the prediction is confirmed across GM, MGG, and SR, and the empirically motivated parameter regime (*ρ* > 0) places Shenhar’s estimate in the inflation zone.

### 4.13 Estimation method: ML vs moment calibration

A natural robustness question is whether the variance-absorption mechanism is mainly an artifact of calibrating the misspecified one-component model to a single dependence summary, *r*_MZ_, rather than to the full bivariate survival likelihood. To separate **model-class** from **estimator-choice** effects, we implemented a bivariate MZ twin log-likelihood for the same misspecified model, integrating over the shared frailty *θ* by Gauss-Hermite quadrature (64 nodes), with proper left-truncation at age 15 and right-censoring corrections. As *σ*_*θ*_ varies, 𝔼[exp(*θ*)] is constrained to its oracle value so that the marginal hazard level is preserved, and the likelihood is optimized by Brent with *m*_ex_ and *b* fixed at their true values, matching the nuisance information available to the moment estimator. The point of the exercise is not whether ML is preferable in the abstract, but whether upgrading the objective from dependence matching to full likelihood removes the bias or only changes how strongly it is expressed.

It does not remove it (Figure 14). Under correct specification (*σ*_*γ*_ = 0), ML recovers the true *σ*_*θ*_ to within +0.0 ± 0.1% (10 seeds, *n* = 50000 pairs), validating the implementation. Under misspecification (*σ*_*γ*_ = 0.40, *ρ* = 0.4), the ML pseudo-true value still inflates *σ*_*θ*_, but only by +3.0 ± 0.1%, whereas *r*_MZ_ matching inflates it by +21.3 ± 0.4%. At heavier heterogeneity (*σ*_*γ*_ = 0.65), the same pattern strengthens: ML +4.8 % versus moment +41.8%. Even with no pleiotropy (*ρ* = 0), both criteria remain positively biased — ML +0.3% and moment +8.8% — showing that the sign of the effect is structural, while its magnitude depends strongly on the fitting criterion.

**Figure 14:**
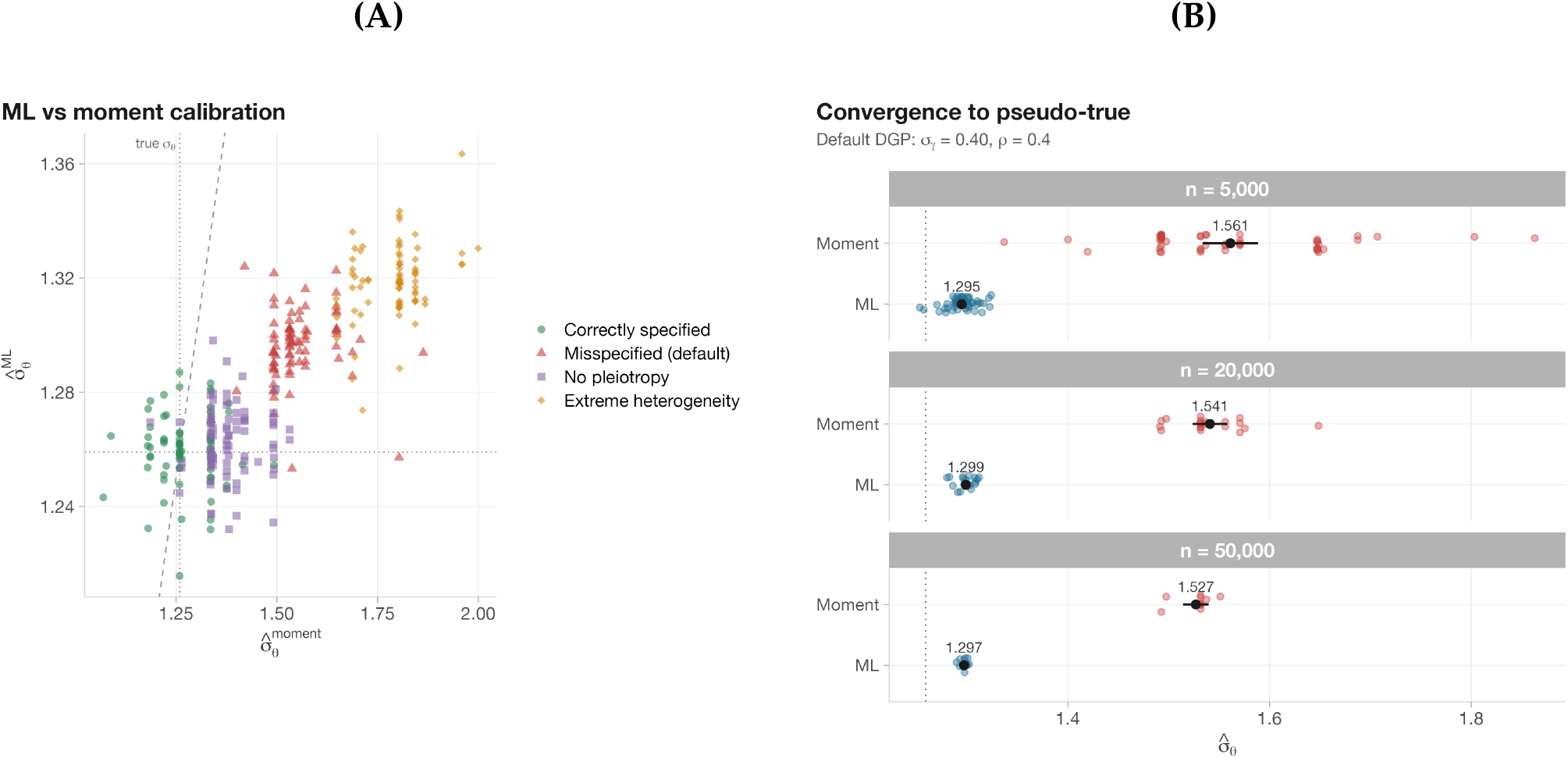
Maximum likelihood vs moment-based calibration. **(A)** 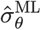 vs 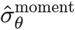 across 4 DGP conditions and 3 sample sizes (50/20/10 seeds at *n* = 5*k*/20*k*/50*k*). Under correct specification (green), both cluster near the identity line. Under misspecification, points migrate rightward (moment inflates more) while ML remains close to the true value — the two estimators converge to different pseudo-true parameters. **(B)** Convergence under the default DGP (*σ*_*γ*_ = 0.40, *ρ* = 0.4): both estimators narrow with increasing *n*, but toward different limits. ML (orange) converges near the true *σ*_*θ*_ (dotted line) while moment (teal) converges to a substantially inflated pseudotrue value.

This is exactly the criterion-dependence emphasized by (White, 1982): under misspecification, the pseudo-true parameter depends on the objective. For ML, it is the parameter that minimizes the Kullback-Leibler discrepancy between the true and fitted joint survival laws; for moment calibration, it is the parameter that reproduces the chosen summary, here *r*_MZ_. These targets need not coincide, and in our setting they do not. The omitted *γ* pathway loads directly into twin dependence, so any one-component approximation must express some of that signal through *σ*_*θ*_; but a dependence-only criterion cannot distinguish whether concordance arises from intrinsic frailty or omitted extrinsic heterogeneity, and therefore asks the model to explain the discrepancy through the single summary most sensitive to absorbed variance.

The reason ML is less inflated is not that it escapes misspecification, but that it is partially regularized by the marginal survival structure. Raising *σ*_*θ*_ improves the fit to excess within-pair dependence, yet it also distorts the marginal age-at-death distribution and the overall joint survival surface, which restrains how much omitted *γ* can be loaded into *σ*_*θ*_. The *r*_MZ_ calibrator has no such counterweight: because it sees only dependence, it channels much more omitted variation into the shared frailty estimate. The robustness check therefore reveals a two-layer vulnerability: **misspecification bias survives objective upgrading**, confirming a model-class problem, while **objective choice changes how much of that bias is expressed as** *σ*_*θ*_ **inflation**, with *r*_MZ_ matching amplifying the effect by about 7 ×.

## 5 Discussion

### 5.1 What is threatened in Shenhar’s estimate (and what is not)

Shenhar’s **mathematical framework** is valid: under its stated assumptions, setting *m*_ex_ = 0 genuinely reveals higher intrinsic heritability. The vulnerability is that calibration is not intrinsically robust to omitted heritable extrinsic structure. If extrinsic mortality susceptibility is heritable and pleiotropic with intrinsic aging — conditions supported by immunogenetic evidence — then the single-parameter calibration conflates two distinct genetic contributions, so the extrapolated “intrinsic” estimate reflects intrinsic aging variance plus whatever heritable extrinsic susceptibility was absorbed into the intrinsic parameter during calibration, rather than intrinsic aging alone — a standard consequence of unmodeled heterogeneity in frailty-type mortality models (Vaupel et al., 1979). The ML-versus-moment comparison (§4.13) sharpens this interpretation: the upward pull is partly **structural**, because omitted *γ* shifts the one-component model toward an inflated pseudo-true *σ*_*θ*_, and partly **estimator-specific**, because calibrating only *r*_MZ_ magnifies that shift by reading the omitted pathway almost entirely as twin dependence.

Extensive simulations strongly suggest this is a quantitatively characterized and structured mechanism with specific boundary conditions, dose-response behavior, and model-general expression. The primary evidence is inflation of the fitted intrinsic frailty parameter: *σ*_*θ*_ is inflated by +22.1% under misspecification (§4.1), translating to +49% variance inflation. This absorption is not limited to a scalar shift — it leaves a structural fingerprint in the bivariate twin survival surface (48× ISE ratio; §4.5) and in age-specific conditional twin correlations (excess at all ages, peaking at age 80 §4.6). A classical ACE structural equation model shows *C* ≈ 0 under all conditions (§4.7), confirming that the heritable extrinsic frailty is absorbed into the additive genetic component rather than shared environment — so the identification failure is not specific to Falconer’s formula. All residual patterns vanish under correct specification. Conceptually, *σ*_*θ*_ inflation is the first-order effect; dependence surfaces, age-conditional correlations, and *h*^2^ are diagnostic projections of that parameter shift.

An apparent objection arises from Shenhar et al.’s own supplementary analysis (their Fig S2), which shows that “variation in extrinsic mortality alone fails to account for the observed correlations” — varying *m*_ex_ as the sole source of individual heterogeneity cannot reproduce observed *r*_MZ_. A reader might interpret this as evidence against heritable extrinsic susceptibility driving twin concordance. In fact, their finding is *consistent with* — and a prerequisite for — the bias mechanism we identify. Shenhar’s Fig S2 tests whether extrinsic heterogeneity *alone* (without intrinsic heterogeneity) can produce the observed correlations; the answer is no. Our mechanism asks a different question: what happens when both *θ* and *γ* exist in the true DGP but the fitted model omits *γ* and estimates only *σ*_*θ*_? Because extrinsic heterogeneity alone is insufficient to explain the correlations (as Shenhar confirm), the calibrated model has no channel through which to represent the extrinsic component of twin concordance — which is precisely why the one-parameter calibration absorbs it into *σ*_*θ*_. Shenhar’s finding establishes a necessary condition for the absorption mechanism: if *γ* variation *could* explain the correlations on its own, the model might not need to absorb it into the intrinsic parameter.

### 5.2 How large is the bias in plausible regimes?

The bias is established at three distinct epistemic levels: an analytic sign result under first-order approximation (Proposition B1, Section B), simulation-based quantitative magnitude across model families and DGP variants (§4), and empirically anchored sensitivity to externally informed parameter values (§4.11). Several bias magnitudes appear throughout this paper; they are not competing estimates but successive refinements across these levels:

1. **Default demonstration** (+7.6 pp net). At the default parameters (*σ*_*γ*_ = 0.40, *ρ* = 0.4), the misspecification produces +11.4 pp of upward inflation offset by 3.7 pp of downward attenuation from extrinsic noise (§4.2). This is the worked example; it establishes the mechanism but makes no claim about the empirically correct parameter values.
2. **Anchored sensitivity regime** (+2.8 to +18.9 pp, mean 9 pp). Within the externally informed restricted range (*σ*_*γ*_ ∈ [0.30, 0.65], *ρ* ∈ [0.20, 0.50]), the bias varies by roughly an order of magnitude (§4.11). At the upper-bound point estimate (*σ*_*γ*_ = 0.65, *ρ* = 0.35), it reaches +15.7 pp. This range is our primary quantitative result.
3. **Alternative calibration robustness** (still positive, magnitude changes). Multi-target calibration fitting five summary statistics yields +6.4 pp, versus +10 pp under *r*_MZ_-only calibration on the same simulation draw (a dedicated draw distinct from the canonical seed; see Section A.6). The direction is robust; only the magnitude shifts with the fitting criterion.

Under even the most cautious assumptions within this range, the bias can materially reduce Shenhar’s 50% estimate. The anchored ceiling (*σ*_*γ*_ = 0.65) was intentionally set well below the bridge-implied parameterization (*σ*_*γ*_ ≈ 1.47) to ensure understated rather than overstated conclusions; bias continues to increase monotonically beyond this ceiling, reaching +32 pp at the bridge-implied value (Figure 12, panel B).

Applying the anchored bias range as a conditional correction to Shenhar’s 50% estimate yields an adjusted intrinsic heritability of approximately 0.50 − Δ, where Δ ∈ [2.8 pp, 18.9 pp], i.e., roughly 31–47%. This corrected range assumes full within-MZ extrinsic concordance (*r*_*γ*, MZ_ = 1); if concordance is lower, the implied upward bias is smaller (e.g., ≈ 6.1 pp at *r*_*γ*, MZ_ = 0.5), shifting the corrected *h*^2^ upward by ∼ 2.4 pp. Across the plausible 0.7–0.9 band for pre-antibiotic Danish MZ twins (§5.6), the shift is small (∼ 0.6–1.8 pp), so the headline range is robust. This is not an estimate of the true intrinsic heritability — it is a what-if correction under the assumptions of our stylized DGP, the anchored (*σ*_*γ*_, *ρ*) regime, and the premise that the dominant distortion in Shenhar’s estimate is omitted extrinsic heterogeneity. Nonetheless, even the lower bound of the adjusted range (31%) substantially exceeds classical twin-study estimates (typically < 25%), suggesting that Shenhar’s core qualitative finding —that intrinsic heritability is higher than observed heritability — may survive correction, albeit at reduced magnitude. If assortative mating inflates the twin correlations that Shenhar calibrates to (Ruby et al., 2018), the correction grows. These interact multiplicatively with the anchored parameter range, so the adjusted interval should be read as indicative, not definitive.

Crucially, this upward bias is not an idiosyncrasy of any one mortality architecture. Across 50 seeds for the main GM conditions and 20 seeds per model for the three-model comparison, Misspecified inflation is consistently +9–10 pp across all three model families (§4.9, Figure 9), and all three extrinsic frailty constructions produce overlapping CIs (+7.6 to +8.7 pp; Appendix, Section A.12) — indicating that the bias is a general property of calibrate-then-extrapolate estimation under omitted extrinsic heterogeneity.

### 5.3 What reanalysis would resolve it

All-cause twin survival data alone cannot uniquely identify a decomposition of genetic variance into “intrinsic” versus “extrinsic” pathways without additional structure — distributional signatures can reveal the presence of misspecification (§4.5, Section 4.5), but not the partition itself. To break this identification trap, twin survival models must move beyond scalar calibration targets. Three paths are most promising.

The most immediately actionable diagnostic is the bivariate dependence surface analysis demonstrated in §4.5. Rather than calibrating to a single summary statistic (*r*_MZ_), future estimation pipelines should evaluate model fit against the full age-structured cross-ratio surface *R*(*t*_1_, *t*_2_): if the optimal scalar 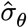 produces a structured residual field — specifically, overpredicting late-life concordance and under-predicting early-life concordance — the single-component assumption is falsified and the resulting heritability estimate should be treated as suspect. We have demonstrated this approach in simulation: the characteristic early-deficit/late-excess pattern vanishes under correct two-component specification (§4.5, Section 4.5). Applying the same diagnostic to Shenhar’s cohort data, via posterior predictive checks on MZ/DZ bivariate survival surfaces, would test whether the fingerprint predicted by our mechanism is present in the empirical data.

When historical registries include cause-of-death information, the identification problem can be addressed more directly through a competing-risks framework. By separating intrinsic and extrinsic death events, a hierarchical competing-risks likelihood can identify the cross-correlation *ρ* from discordant pairs (e.g., where one twin dies of infection and the other of senescent decline), neutralizing the omitted-variable absorption. When cause-of-death data is unavailable, an alternative is to anchor one of the latent frailty distributions using observed genomic data — for example, partitioning extrinsic frailty into an observed polygenic-score component and a residual latent term. By making a fraction of the extrinsic concordance observable, such anchoring breaks the mathematical symmetry that allows the single intrinsic parameter to absorb all familial concordance. These extensions go beyond the scope of the present simulation study; we note them as natural next steps for the field.

The true intrinsic heritability could plausibly be lower than Shenhar’s 50% if extrinsic susceptibility is heritable and unmodeled. Ruby et al.‘s estimates (< 7%) highlight that multiple, distinct biases can operate in the same downward direction: they identify assortative mating as inflating *observed* twin correlations, while we identify omitted extrinsic heterogeneity as inflating the *calibrated* intrinsic parameter. These two corrections are analytically independent — ours addresses the calibrate-then-extrapolate step, theirs addresses the input data — and are potentially additive. Applying our correction alone reduces Shenhar’s 50% estimate by +2.8 to +18.9 pp depending on parameter assumptions; if assortative mating further inflates the twin correlations that Shenhar calibrates to, the combined effect could push the estimate substantially lower. However, Ruby et al. use genealogical (non-twin) data with different methodology, so direct combination requires care.

### 5.4 Low modern extrinsic mortality does not resolve calibration-stage misspecification

Modern extrinsic mortality rates in contemporary cohorts (*m*_ex_ ≈ 3 × 10^−4^; B. Shenhar, personal communication) are far below historical levels. Our dose-response analysis agrees that, *conditional on a correctly identified intrinsic parameter*, the additional distortion from evaluating the model at a modern *m*_ex_ is small. But this concerns the **evaluation stage**, not the **calibration stage** at which the key parameter is fitted.

Shenhar’s published estimate is obtained by calibrating a one-component frailty model to historical cohorts with substantially larger extrinsic mortality (*m*_ex_ ≈ 0.004) and then extrapolating that fit to the modern regime. If the historical data-generating process contains omitted familial extrinsic heterogeneity, then the fitted one-component variance converges to a pseudo-true value 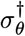, not in general to the intrinsic variance *σ*_*θ*_. Evaluating the extrapolated model in the limit *m*_ex_ → 0 therefore yields 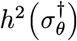 rather than *h*^2^(*σ*_*θ*_). Low modern extrinsic mortality can make the issue second-order for present-day *evaluation*, while remaining first-order for interpretation of the published *σ*_*θ*_ as an intrinsic biological parameter. The modern asymptotic regime simplifies downstream evaluation; it does not retroactively identify a historically misspecified fit.

Even granting that extrinsic mortality is much lower in modern cohorts, that does not rescue the published estimate if the inflation entered earlier at calibration. In our analyses, omitted familial extrinsic susceptibility materially shifts the published intrinsic frailty parameter itself, inflating the headline *σ*_*θ*_ by roughly +22% (§4.1) — not a negligible perturbation but a material distortion of the specific reported estimate. The magnitude of Shenhar’s own correction — from observed *h*^2^ ≈ 0.20 to intrinsic *h*^2^ ≈ 0.50 — demonstrates that extrinsic mortality is not negligible in the identifying cohorts. The relevant contrast is therefore not high versus low modern *m*_ex_, but correctly specified versus misspecified historical calibration.

A related objection is that modern twin studies such as SATSA already report substantial late-life *h*^2^ without invoking the historical correction machinery considered here. We do not dispute that modern twin data may support high heritability. Our claim is narrower: such evidence bears on the magnitude of familial resemblance in contemporary cohorts, not on whether a historically calibrated one-component frailty variance isolates a purely intrinsic biological component. SATSA and Shenhar et al. involve different designs, different estimands, and different identification assumptions. There is no contradiction between (a) high late-life heritability in modern cohorts, (b) negligible direct correction at contemporary *m*_ex_, and (c) contamination of a historically calibrated one-component parameter by omitted familial extrinsic structure.

### 5.5 Scope of the claim

To avoid overstatement, it is useful to distinguish three questions: (i) whether the misspecification mechanism exists in principle, (ii) whether it changes the estimand identified by the historical calibration, and (iii) how large the induced discrepancy is under plausible regimes. Our cross-model sign-reversal results speak to (i), the pseudo-true-parameter argument speaks to (ii), and the anchored sensitivity analysis speaks to (iii). Low modern *m*_ex_ bears mainly on (iii) at the evaluation stage; it does not by itself answer (ii).

Our claim should therefore be read narrowly. We do not argue that contemporary cohorts require large extrinsic correction, nor that late-life heritability in modern twin data must be low. Rather, we argue that a one-component frailty variance calibrated on historical cohorts cannot be interpreted as a purely intrinsic parameter unless omitted familial extrinsic structure is either modeled explicitly or shown to be negligible in the calibration data. Our strongest claim is therefore interpretive: the published one-component estimate should be treated as a model-dependent historical-fit parameter unless omitted familial extrinsic structure is shown to be negligible in the calibration cohorts.

### 5.6 Limitations

Our primary calibration uses stochastic bisection on *r*_MZ_ because it reproduces Shenhar’s estimating target, but we now benchmark it against full bivariate maximum likelihood within the same misspecified one-component model. This comparison shows that the positive bias is not an artifact of using moments: omitted *γ* still induces an inflated pseudo-true *σ*_*θ*_ under ML, although dependence-only matching expresses substantially more of that misspecification as *σ*_*θ*_ inflation. To further verify robustness to the fitting criterion, we re-estimated *σ*_*θ*_ using a multi-target minimum-distance calibration that simultaneously fits five summary statistics (*r*_MZ_, mean and SD of age-at-death, *q*_25_, *q*_75_): the bias persists at +6.4 pp, versus +10 pp under *r*_MZ_-only calibration on the same draw (Appendix, Section A.6; the small difference from Table 5′s +7.6 pp reflects Monte Carlo variability across independent simulation draws). Moreover, the misspecified model’s best-compromise fit leaves detectable residual misfit (e.g., *q*_25_ underpredicted by 2.2 years) — consistent with the broader frailty-survival literature showing that model misspecification can bias both absolute risk and heterogeneity measures (Gasparini et al., 2019) confirming that the presence of omitted extrinsic frailty is in principle detectable from distributional features beyond twin correlations — though detecting misspecification is not the same as uniquely identifying the intrinsic/extrinsic decomposition, which requires the additional structure discussed in §5.3. Similarly, the age-structured residual in bivariate dependence (§4.5) is a mechanism-consistent diagnostic — its directionality matches the omitted-variable prediction — but is not a unique signature; alternative misspecifications could also distort joint dependence, underscoring the need for the external anchoring and negative controls provided elsewhere in the paper.

The anchoring of *σ*_*γ*_ and *ρ* to published liability-scale and GWAS estimands is approximate, and the sensitivity range (*σ*_*γ*_ ∈ [0.30, 0.65], *ρ* ∈ [0.20, 0.50]) is restricted below empirically motivated levels rather than calibrated to them. We evaluate robustness by sweeping both parameters across wide ranges, and the bias is positive throughout the upper-left triangle of the parameter space (where pleiotropy and extrinsic heterogeneity are both non-negligible). COVID-19 severity provides an existence proof that genetic correlation between infection susceptibility and aging pathways is non-negligible, but it represents a single novel pathogen with specific immunological characteristics (ACE2-mediated entry, cytokine storm). Pre-antibiotic-era infection mortality in Danish cohorts born 1870–1900 was dominated by tuberculosis, pneumonia, diphtheria, and enteric infections — pathogens engaging different arms of the immune system. GWAS evidence strengthens the pleiotropic case: signals in the HLA class II region (HLA-DQA1/DRB1) reach genome-wide significance for both TB susceptibility (Schurz et al., 2024) and parental lifespan (Joshi et al., 2017), consistent with shared immunogenetic architecture, while Nordic-specific TB associations are documented in FinnGen (Tervi et al., 2023). The *ρ* range we adopt is informed by the convergence of COVID-19, HLA-longevity overlap, and Sørensen adoption-study evidence (Petersen et al., 2010; Sørensen et al., 1988), not by quantitative transfer from any single pathogen to historical cohorts.

Our simulation is framed around heritable extrinsic frailty — the case most directly motivated by immunogenetic evidence — but the calibration-absorption mechanism operates on any source of within-pair concordance in mortality-relevant extrinsic susceptibility, whether genetic or shared-environmental (§4.8, footnote). The genetic case is our primary target; the broader familial generalization is a secondary implication that widens the scope of the vulnerability but does not change its structure. The DGP assumes MZ twins share extrinsic susceptibility identically (correlation 1) and DZ twins at 0.5, paralleling the standard ACE model for intrinsic frailty. In reality, within-pair concordance in extrinsic susceptibility reflects both genetic sharing and individual-specific exposure noise (e.g., different infection contacts), so the effective within-MZ-pair correlation may be below 1. To quantify this sensitivity, we swept the within-MZ extrinsic concordance *r*_*γ*,MZ_ from 0 to 1 (DZ concordance scales as 0.5 ⋅ *r*_*γ*,MZ_; 20 seeds per level; Appendix, Section A.13). The bias rises monotonically from 0.1 pp at *r*_*γ*,MZ_ = 0 (matching the nonfamilial control) to 8.5 pp at *r*_*γ*,MZ_ = 1 (matching the Misspecified condition), remaining positive across the entire range (Figure 15). Crucially, even at *r*_*γ*,MZ_ = 0.5 the net bias is already 6.1 pp — within 2.4 pp of the perfect-concordance benchmark. Once extrinsic concordance is moderate (*r*_*γ*,MZ_ > 0.4), additional concordance adds little; conclusions are not sensitive to whether *r*_*γ*,MZ_ is 0.7 or 1.0. For MZ twins raised in the same household in pre-antibiotic Denmark (born 1870–1900), the effective concordance is likely 0.7–0.9, driven by both shared genetic susceptibility and shared household infection exposure; at *r*_*γ*,MZ_ = 0.7, the bias is 6.7 pp, and at 0.85 it is 7.9 pp. The bridge analysis (§3.4) assumes *r*_*γ*,MZ_ = 1; the concordance sweep shows that relaxing this to the plausible 0.7–0.9 range reduces the bias by only ∼ 0.6–1.8 pp. Our reported estimates correspond to the *r*_*γ*,MZ_ = 1 case and should be read as upper bounds, but the sensitivity analysis shows these bounds are tight: the plausible range of concordance produces bias within 1–2 pp of the maximum.

**Figure 15:**
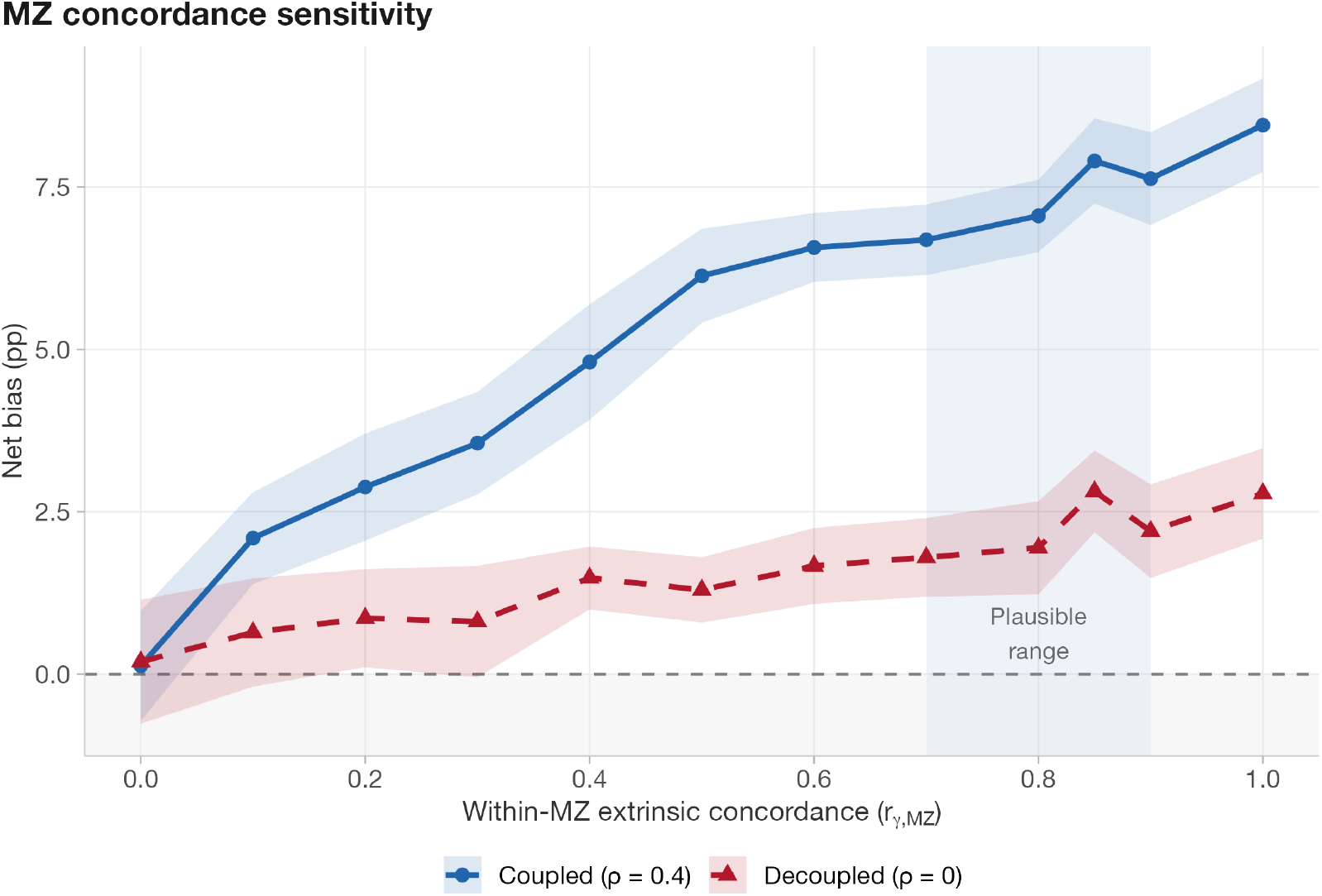
Sensitivity of net bias to within-MZ extrinsic concordance (*r*_*γ*,MZ_). Blue solid: coupled sweep (*ρ* = 0.4; varying concordance also varies effective pleiotropy). Red dashed: decoupled sweep (*ρ* = 0; isolates pure concordance effect). Ribbon: 95% CI from 20 seeds per level. Blue band: plausible range (0.7–0.9) for pre-antibiotic Danish MZ twins. The coupled bias rises to 8.5 pp at full concordance and is already within 2.4 pp of its maximum by *r*_*γ*,MZ_ = 0.5. The decoupled trace shows that extrinsic concordance alone produces positive bias even without pleiotropy; the gap between the two traces quantifies the amplifying role of *ρ* > 0. To disentangle these two effects, we ran a second concordance sweep at *ρ* = 0 (red dashed trace in Figure 15), isolating the pure extrinsic concordance effect from pleiotropy amplification. At *ρ* = 0, the bias from concordance alone is smaller (e.g., 2.8 pp at *r*_*γ*,MZ_ = 1) but still positive, confirming that extrinsic concordance contributes independently of pleiotropy; the difference between the coupled and decoupled traces quantifies the amplifying role of *ρ* > 0.

The extrinsic frailty DGP uses multiplicative log-normal heterogeneity 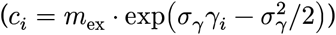. To verify that the bias is not driven by this distributional choice, we tested two alternative constructions: additive perturbation (*c*_*i*_ = *m*_ex_ + *σ*_*γ*_*γ*_*i*_, truncated at zero) and gamma-distributed frailty, each with mean extrinsic hazard held fixed at *m*_ex_. Across 10 seeds per construction, the bias is +8.1 ± 0.4 pp (log-normal), +8.7 ± 0.4 pp (additive), and +7.6 ± 0.6 pp (gamma) — all with overlapping 95% CIs (Appendix, Section A.12). The 1 pp spread across constructions, relative to a 8 pp mean effect, confirms that the qualitative conclusion does not depend on the parametric form of extrinsic heterogeneity.

## 6 Conclusion

Taken together, the analyses above establish that omitting heritable extrinsic frailty from a single-parameter calibration does not just shift a point estimate — it produces a structured, directional, and model-general distortion of the fitted latent variance *σ*_*θ*_, with diagnostic consequences visible in the joint twin survival surface and the age-pattern of twin dependence. The following points summarize the principal findings.

1. Shenhar’s calibrate-then-extrapolate framework is mathematically coherent under its assumptions, but its “intrinsic” estimate is not empirically identified as purely intrinsic if heritable extrinsic frailty exists and is omitted at calibration.
2. The primary estimand is inflation of the fitted intrinsic frailty parameter (*I*_*σ*_): *σ*_*θ*_ is inflated by +22.1 % (95% CI: 21.5 to 22.7%; 20 seeds). Because *I*_var_ = (1 + *I*_*σ*_)^2^ − 1, this corresponds to +49% variance inflation — the upstream distortion from which downstream biases in any monotone summary follow (Corollary B3). The downstream Falconer *h*^2^ bias is +9.2 pp (95% CI: 8.7 to 9.7 pp; 50 seeds).
3. The inflation is model-general: +22% (GM), +18% (MGG), +14% (SR) dispersion inflation across three mortality model families; robust to alternative extrinsic frailty constructions (log-normal, additive, gamma).
4. The misspecified model’s joint twin survival surface is 48× worse (ISE) at reproducing the true bivariate dependence structure than the recovery model. Age-specific conditional correlation shows excess dependence at all ages, peaking at age 80 (Δ*r* = 0.048). These structural fingerprints vanish under correct specification.
5. Classical ACE twin decomposition yields *C* ≈ 0 under all conditions — the heritable extrinsic frailty is absorbed into the additive genetic component *A*, not shared environment *C*. Switching from Falconer to a structural equation model does not resolve the identification failure.
6. The bias scales monotonically with the heritable fraction of extrinsic mortality; the anchored sensitivity range (+2.8 to +18.9 pp) uses a ceiling deliberately set below bridge-implied levels.
7. A correctly specified two-component model recovers *σ*_*θ*_ to within −3.2% of its true value, eliminating both the scalar inflation and the structured age-specific misfit — confirming that the distortion is specifically attributable to omitting extrinsic frailty.
8. Negative genetic correlation (*ρ* < 0) reverses the bias sign — a falsifiable prediction confirmed in all three model families.
9. The ML robustness check reveals a two-layer vulnerability: omitted *γ* induces a structurally inflated pseudo-true *σ*_*θ*_ even under full ML, while matching only *r*_MZ_ amplifies that same misspecification by about 7 × because the omitted pathway loads directly into twin dependence.

## 7 Acknowledgements

GPT-5.2-Pro (OpenAI) and Gemini-3-Pro (Google) were used during development to review the simulation design, verify that the calibration procedure parallels Shenhar et al.‘s methodology, and provide structural feedback on manuscript organization. MiniMax-M2.5 (MiniMax), GLM-5 (Zhipu AI), GPT-5.2-Pro, and Gemini 3 Pro were used to review and assist in deriving the formal proofs in Section B. Claude Opus 4.6 (Anthropic) was used for code generation, drafting assistance, and proof typesetting. All code, analytical results, and final text were reviewed and revised by the author.

APPENDICES

## A Appendix: Computational implementation

### A.1 Pipeline architecture

All simulations use the `{targets}` reproducible pipeline framework (Landau 2021) with `{crew}` for parallel execution on 10 local workers (Apple M4 Pro, 14 cores, 24 GB RAM). The pipeline defines 207 targets organized in a directed acyclic graph (DAG), with static branching (`tar_map`) for the 81-cell sensitivity sweep and 56-cell anchored sweep. A single master seed (66829) propagates deterministic sub-seeds to every target, ensuring full reproducibility.

R 4.4.2 is used throughout. Key packages: `MASS` (multivariate normal twin parameters), `lamW` (Lambert W closed-form GM solver), `mvtnorm`, `Rcpp`.

### A.2 Gompertz–Makeham solver

Lifespan inversion for the Gompertz–Makeham model uses the Lambert W function. The cumulative hazard 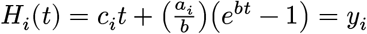 (where *y*_*i*_ = − log(*V*_*i*_), *V*_*i*_ ∼ Uniform(0, 1), *a*_*i*_ = exp(*θ*_*i*_), and *c*_i_ is the individual extrinsic hazard) has the closed-form solution:

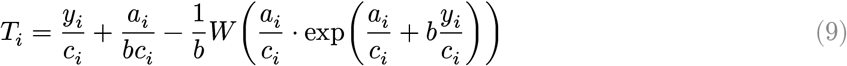

For pure Gompertz 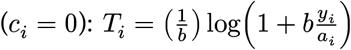. When log *W*_arg_ > 500, the Lambert W function is computed in log space via Newton iteration (*w* + log *w* = *z*) to avoid floating-point overflow. A bisection fallback handles remaining edge cases.

### A.3 SR Euler–Maruyama solver (Rcpp)

The saturating-removal (SR) damage model requires numerical integration of the stochastic differential equation:

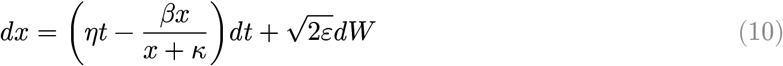

with death occurring when damage *x* exceeds the individual threshold *X*_*c,i*_ or an independent Poisson extrinsic event occurs. We integrate via Euler–Maruyama with timestep Δ*t* = 1/52 (weekly).

The inner simulation loop is compiled to C++ via `Rcpp` for performance. The C++ implementation uses a Xoroshiro128+ pseudorandom number generator for the per-individual noise and extrinsic-event draws, avoiding the overhead of R’s scalar RNG interface. A compact alive-index array skips deceased individuals at each timestep. This yields a measured ≈4.6× speedup over the equivalent vectorized R loop (0.54 s vs 2.5 s per 10,000-pair simulation). Each `{crew}` worker lazy-compiles the C++ source on first SR invocation; the compilation is cached for subsequent calls within the same worker process.

### A.4 MGG solver and Strehler–Mildvan parameterization

The Makeham–Gamma–Gompertz model uses the closed-form cumulative hazard from Shenhar et al.‘s Supplementary Materials. The Strehler–Mildvan coupling sets *b*_*i*_ = *q*_*i*_*b*_0_; the remaining question is how *a*_*i*_ depends on *q*_*i*_. Shenhar et al.‘s bioRxiv preprint (v1) writes the compensatory form in exponential notation: 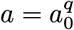. Taking logs gives the equivalent log *a* = *q* ⋅ log *a*_0_ ; in the published *Science* supplement, however, the *q* appears in the denominator — log *a* = (log *a*_0_)/*q* — yielding 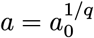. This is the kind of slip that can occur when converting between exponential and log-linear forms during manuscript preparation, and the preprint’s form is clearly intended: only 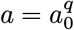 produces Strehler–Mildvan compensation (higher *q* raises *b* but *lowers a*, because *a*_0_ = 10^−5^ < 1), yields hazard-curve crossing, and reproduces Shenhar et al.‘s reported twin correlations and heritability estimates.

Under 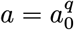, higher *q* raises *b* but lowers *a*, producing the characteristic hazard-curve crossing at 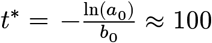 years — the hallmark of Strehler–Mildvan compensation. Figure 16 illustrates the consequences. Panel A confirms that 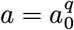 produces converging hazard curves while 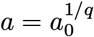 produces diverging ones. Panel B shows cohort survival: the alternative form creates a bimodal mortality pattern in which high-*q* individuals die prematurely (no compensation to restrain their hazard). Panel C confirms the identification: with identical dispersion (*σ*_*q*_ = 0.27) and no extrinsic mortality, 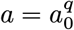 yields *r*_MZ_ = 0.43 and Falconer *h*^2^ = 0.47 — consistent with Shenhar et al.’s published twin correlations — whereas 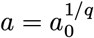 yields *r*_MZ_ = 0.92 and *h*^2^ = 1.01, exceeding unity.

**Figure 16:**
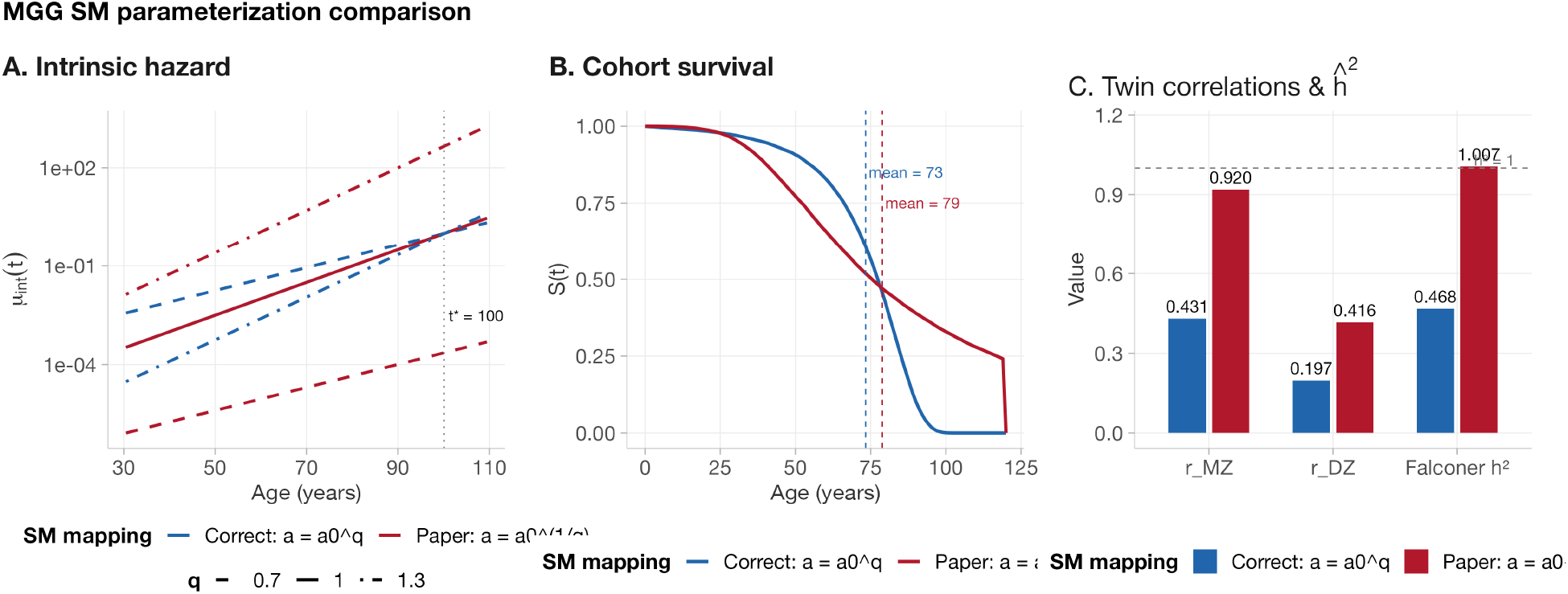
Strehler–Mildvan parameterization comparison. (A) Intrinsic hazard curves for *q* ∈ {0.7, 1.0, 1.3}: the compensatory form 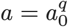 (blue) produces convergence at *t*^∗^; the literal supplement formula 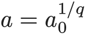 (red) produces divergence. (B) Cohort survival (*n* = 20000, *m*_ex_ = 0): the literal formula creates early mortality in high-*q* individuals. (C) Oracle twin correlations and Falconer *h*^2^ (*σ*_*q*_ = 0.27, *m*_ex_ = 0): 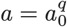 yields *h*^2^ = 0.47, consistent with Shenhar et al.’s published values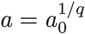 yields *h*^2^ = 1.01.

**Figure 17:**
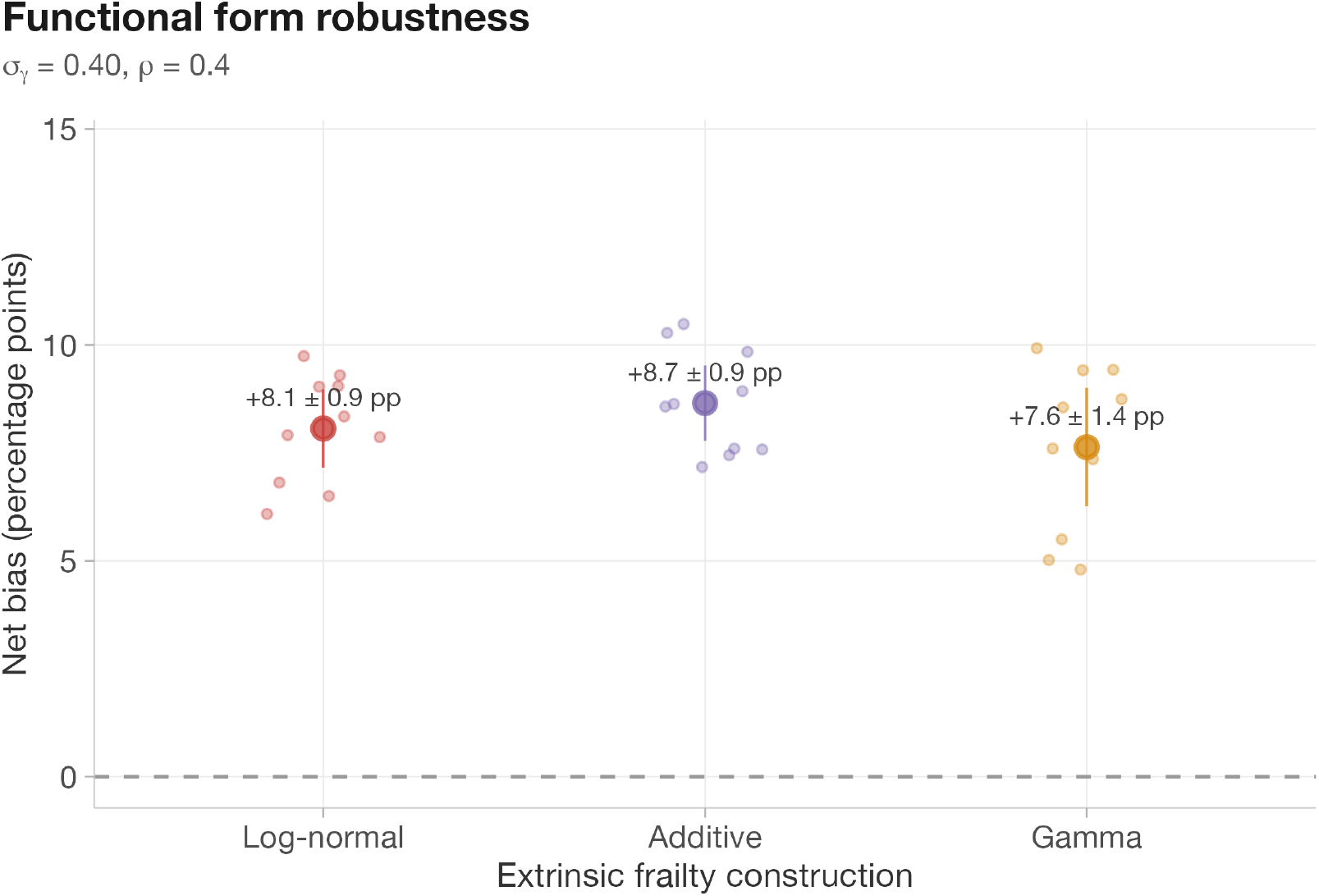
Functional-form robustness (GM model, 10 seeds per construction). Log-normal (+8.1 ± 0.4 pp), additive (+8.7 ± 0.4 pp), and gamma (+7.6 ± 0.6 pp) extrinsic frailty constructions produce overlapping CIs, confirming that the bias does not depend on the parametric form of extrinsic heterogeneity.

We adopt 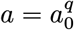 throughout. The inverse CDF is solved via hybrid bisection–Newton iteration with 40 steps.

### A.5 Calibration

All calibration uses 40-iteration bisection on the respective dispersion parameter (*σ*_*θ*_ for GM, *σ*_*q*_ for MGG, 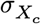 for SR) to match a target *r*_MZ_. The stochastic bisection uses common random numbers (CRN): the same RNG seed is used for each candidate parameter value within a bisection run, reducing variance in the comparison.

### A.6 Multi-target calibration robustness

A potential concern is that the upward bias under misspecification could be an artifact of calibrating the misspecified intrinsic-only model by matching only *r*_MZ_. To test this, we re-estimated *σ*_*θ*_ using a multitarget calibration that simultaneously fits five summary statistics computed from concordant-survivor MZ pairs: *r*_MZ_, the mean and standard deviation of age-at-death, and the 25th and 75th percentiles. The misspecified model’s sole free parameter *σ*_*θ*_ was chosen by one-dimensional optimization (Brent method) minimizing a weighted sum of squared errors across all five targets, with common random numbers ensuring a deterministic objective and fair comparison to the *r*_MZ_-only calibration.

Because the misspecified model has only one degree of freedom and five fitting targets, the system is overidentified — the optimizer finds a best-compromise *σ*_*θ*_ that cannot match all targets simultaneously.

Under multi-target calibration, the inferred intrinsic-only heritability remained biased upward 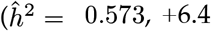 pp relative to the oracle *h*^2^ ≈ 0.51), although less than under *r*_MZ_-only calibration 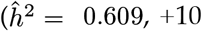 pp). The best-compromise fit leaves detectable residual misfit: the 25th percentile of age-at-death is underpredicted by 2.2 years, and the goodness-of-fit RMSE across all targets is 1.13 years. These residuals confirm that the omitted extrinsic frailty is in principle identifiable from distributional features, and that the upward heritability bias is not specific to the choice of *r*_MZ_ as the sole calibration target.

### A.7 Grid specifications

Sensitivity sweeps used the grids specified in Table 9.

**Table 9.** Grid specifications for all parameter sweeps and robustness checks.

| Grid | Specification |
| --- | --- |
| Full sweep | $\rho \in \{0, 0.1, \dots, 0.8\} \times \sigma_\gamma \in \{0.15, 0.20, \dots, 0.55\}$ (81 cells) |
| Anchored sweep | $\rho \in \{0.20, 0.25, \dots, 0.50\} \times \sigma_\gamma \in \{0.30, 0.35, \dots, 0.65\}$ (56 cells) |
| Dose-response | 15 $m_{\text{ex}}$ values from 0 to 0.012 |
| Negative $\rho$ | $\rho \in \{-0.60, -0.55, \dots, 0.80\}$ by 0.05 (29 points) |
| $m_{\text{ex}}$ split | 21 heritable fractions from 0 to 1 by 0.05 |
| $\sigma_\gamma$ bridge | 57 values from 0.1 to 1.5 by 0.025 |
| Extended $\sigma_\gamma$ | 7 values from 0.70 to 1.47 ($\rho = 0.35$; 10 reps each) |
| Functional form | 3 constructions (log-normal, additive, gamma; 10 reps each) |
| Multi-model MC | 20 seeds × 3 models (GM, MGG, SR) |
| Heritable fraction (hi-res) | 21 levels from 0 to 1 by 0.05 (20 reps each) |

**Table 10.**
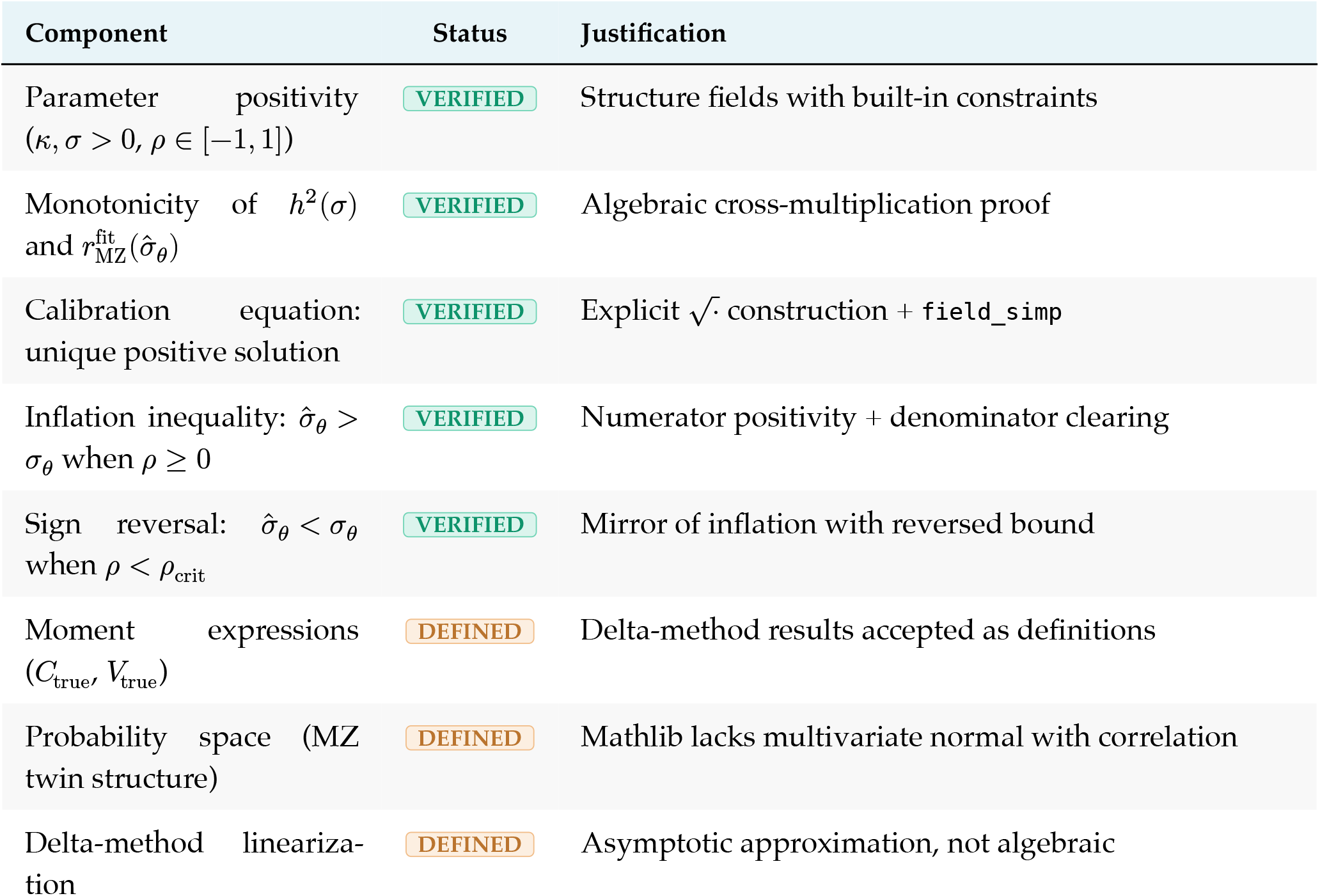
Verification status of each component. **VERIFIED** indicates a machine-checked proof; **DEFINED** indicates a mathematical definition accepted without derivation from measure theory.

| Component | Status | Justification |
| --- | --- | --- |
| Parameter positivity<br>($\kappa, \sigma > 0, \rho \in [-1, 1]$) | **VERIFIED** | Structure fields with built-in constraints |
| Monotonicity of $h^2(\sigma)$<br>and $r_{\text{MZ}}^{\text{fit}}(\hat{\sigma}_\theta)$ | **VERIFIED** | Algebraic cross-multiplication proof |
| Calibration equation:<br>unique positive solution | **VERIFIED** | Explicit $\sqrt{\cdot}$ construction + field_simp |
| Inflation inequality: $\hat{\sigma}_\theta > \sigma_\theta$<br>when $\rho \geq 0$ | **VERIFIED** | Numerator positivity + denominator clearing |
| Sign reversal: $\hat{\sigma}_\theta < \sigma_\theta$<br>when $\rho < \rho_{\text{crit}}$ | **VERIFIED** | Mirror of inflation with reversed bound |
| Moment expressions<br>($C_{\text{true}}, V_{\text{true}}$) | **DEFINED** | Delta-method results accepted as definitions |
| Probability space (MZ<br>twin structure) | **DEFINED** | Mathlib lacks multivariate normal with correlation |
| Delta-method lineariza-<br>tion | **DEFINED** | Asymptotic approximation, not algebraic |

All models use 50,000 pairs per zygosity. The SR Euler–Maruyama solver is *O*(*n* ⋅ *T* /Δ*t*); with the Rcpp implementation, each 50,000-pair SR simulation completes in ≈2.5 s.

### A.8 Hardware and runtime

Simulations were run on an Apple M4 Pro (14 cores, 24 GB unified memory) under macOS 26 (Tahoe). The `{crew}` controller dispatches up to 10 parallel workers. The pipeline defines 207 targets with static branching for sweeps and per-model MC targets dispatched as independent crew jobs. Inner parallelism (`mclapply` with min(*n*_seeds_, 4) cores) is used within long-running MC targets to exploit the remaining cores on each worker. All parameter sweeps (negative *ρ*, dose-response, pleiotropy isolation) use 20 independent seeds per grid point across all three mortality models (GM, MGG, SR), producing averaged curves with *t*-based 95% CIs. The SR Euler–Maruyama sweeps are the computational bottleneck; the full pipeline — including all multi-seed sweeps, 20-seed multi-model MC uncertainty quantification, and robustness analyses — completes in approximately 20 hours wall-clock time.

### A.9 Heritable-fraction sensitivity

The heritable-fraction sensitivity analysis (§4.10, Figure 10) evaluates bias at 21 fraction levels (0 to 1 by 0.05) with 20 independent seeds per level. Each seed uses an independent RNG offset from the master seed, ensuring distinct twin draws. The 95% CIs shown in Figure 10 are *t*-based intervals 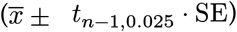 across the 20 seeds at each level.

### A.10 Multi-model Monte Carlo uncertainty

To quantify Monte Carlo uncertainty across mortality model families, we ran 20 independent seeds for each of the three models (GM, MGG, SR). Each seed defines a complete Baseline/Misspecified pipeline: fresh twin-pair generation, calibration (40-iteration bisection with CRN), and evaluation at *m*_ex_ = 0. Oracle *h*^2^ is computed from a separate pure-intrinsic simulation. The three per-model targets (`mc_unc_gm`, `mc_unc_mgg`, `mc_unc_sr`) are dispatched as independent `{crew}` jobs for full parallelism. Within each target, the 20-seed loop uses `mclapply` with 4 cores to further exploit available hardware. Reported CIs are *t*-based intervals 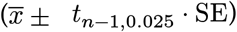 across the 20 seeds. For the SR model, the Rcpp Euler–Maruyama solver is used throughout, with each seed receiving a distinct fast-seed derived from the R RNG state.

### A.11 Extended *σ*_*γ*_ stress test

The extended *σ*_*γ*_ sweep (§4.11, Figure 12 (panel B)) evaluates bias at 7 grid points from *σ*_*γ*_ = 0.70 to *σ*_*γ*_ = 1.47 (the bridge-implied value), with *ρ* fixed at 0.35 and 10 seeds per grid point. This complements the main anchored sweep (which caps at *σ*_*γ*_ = 0.65) by characterizing the bias trajectory beyond the intentionally conservative sensitivity ceiling. CIs are *t*-based intervals (*t*_9,0.025_ = 2.26) across the 10 seeds.

### A.12 Functional form variants

The functional-form robustness check (§4.9, §5.6) tests three constructions for extrinsic frailty heterogeneity, each with mean extrinsic hazard held fixed at *m*_ex_ = 0.004:

1. **Log-normal** (baseline): 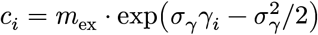, with *γ*_*i*_ ∼ 𝒩(0, 1). This is the default DGP used throughout the paper.
2. **Additive**: *c*_*i*_ = *m*_ex_ + *σ*_*γ*_*γ*_*i*_, truncated at zero. The additive construction models heterogeneity as a shift in the extrinsic hazard rather than a multiplicative perturbation.
3. **Gamma**: *c*_*i*_ ∼ Gamma(*k, k*/*m*_ex_), where *k* is chosen to match the coefficient of variation of the log-normal construction at the same *σ*_*γ*_. This ensures that the first two moments of the extrinsic hazard distribution are matched across constructions.

Each construction is evaluated with 10 independent seeds at the default parameters (*σ*_*γ*_ = 0.40, *ρ* = 0.4) under the GM mortality model. The 95% CIs are *t*-based intervals (*t*_9,0.025_ = 2.26) across seeds. All three constructions use the same calibration machinery (40-iteration bisection on *r*_MZ_) and the same evaluation protocol (Falconer *h*^2^ at *m*_ex_ = 0).

### A.13 Within-MZ extrinsic concordance sensitivity

The main analysis assumes MZ twins share extrinsic susceptibility identically (within-pair correlation *r*_*γ*,MZ_ = 1). To quantify sensitivity to this assumption, we swept *r*_*γ*,MZ_ from 0 to 1 in steps of 0.1 (plus 0.85), generating 50,000 MZ and DZ pairs at each level under the default DGP (*σ*_*γ*_ = 0.40, *ρ* = 0.4), with 20 independent seeds per level. For each *r*_*γ*,MZ_, we decomposed the extrinsic frailty into a shared genetic component and an individual-specific component: 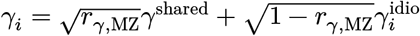, where *γ*^shared^ is common to both twins in an MZ pair and 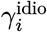 is independent. DZ pairs use 0.5 ⋅ *r*_*γ*,MZ_ for the shared component. Reducing *r*_*γ*,MZ_ also reduces the effective within-person pleiotropy: 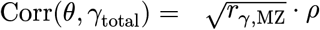, since only the genetic component of extrinsic susceptibility is pleiotropic with intrinsic frailty. At each level, we recalibrated 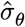 and computed the misspecification bias relative to the oracle. The resulting bias profile rises monotonically from 0.1 pp (matching the nonfamilial control) to 8.5 pp (matching the Misspecified condition), with near-saturation above *r*_*γ*,MZ_ ≈ 0.4.

### A.14 Bivariate dependence diagnostic

The bivariate dependence diagnostic (§4.5, Figure 4) compares the joint survival structure of MZ and DZ twin pairs under three DGPs: (i) the true two-component model (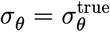, *σ*_*γ*_ = 0.40, *ρ* = 0.4, heritable extrinsic frailty with individual-level heterogeneity); (ii) the misspecified one-component fit (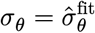, *σ*_*γ*_ = 0), where 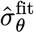 was calibrated to match *r*_MZ_ of the true DGP; and (iii) a recovery refit (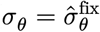, *σ*_*γ*_ = 0.40, *ρ* = 0.4), where 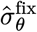 was calibrated under the correctly specified two-component model (§3.3).

All three DGPs are evaluated on a held-out seed block (master seed +200000), with independent seeds offset by 1000 between DGPs, to prevent information leakage from the calibration stage. Each DGP generates *n* = 50000 twin pairs (MZ and DZ separately) under the Gompertz–Makeham closed-form solver. Pairs where either twin dies before age 15 are excluded, consistent with the concordant-survivor conditioning used throughout the paper.

The 1D dependence ratio *R*(*t*) is computed on a grid from age 20 to 100 (1-year spacing) as 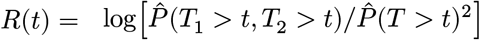, where the numerator is the empirical joint survival fraction and the denominator is the squared marginal survival fraction (pooling both twins). Grid points are masked (set to `NA`) if the joint count falls below 30 or the marginal survival probability falls below 3%, to prevent ratio instability in the tails.

The 2D bivariate surface *R*(*t*_1_, *t*_2_) is computed on a coarser grid (age 20 to 95, 2-year spacing) for computational tractability. The same masking thresholds apply. Residuals Δ*R* = *R*_model_ − *R*_true_ are displayed as diverging heatmaps centered at zero.

Two scalar summaries are reported: the integrated absolute error (IAE), defined as the mean absolute deviation 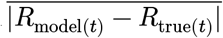 over non-masked grid points; and the peak deviation age, the age at which |*R*_misspec(*t*)_ − *R*_true(*t*)_| is maximized. The IAE ratio (misspecified/recovery) provides a scale-free measure of the improvement from correct specification.

## B Appendix: Formal proof of calibration inflation

This appendix provides a first-order (delta-method) proof that omitting heritable extrinsic frailty from the fitted model inflates the calibrated intrinsic dispersion parameter. The qualitative predictions — direction and mechanism of bias — are corroborated by exact simulation throughout the main text. **Scope note:** the equal-noise approximation underlying Proposition B1 is quantitatively reliable in the anchored sensitivity regime (*σ*_*γ*_ ∈ [0.30, 0.65]) but becomes unreliable at large extrinsic frailty (e.g., *σ*_*γ*_ ≈ 1.47, the bridge-implied value). In such regimes, the formal result establishes the sign and mechanism; the exact simulation results constitute the operative quantitative statement. See the detailed scope discussion following Proposition B1.

### B.1 Setup: true DGP and fitted model

#### True data-generating process

Let *b* > 0 and *m*_ex_ > 0 be fixed. For individual *i*, define intrinsic frailty *θ*_*i*_ = *σ*_*θ*_(*G*_*i*_ + *E*_*i*_), where *G*_*i*_ ∼ 𝒩(0, 1) is the additive genetic component and *E*_*i*_ ∼ 𝒩(0, 1) is the non-shared environmental component. (We use *G, E* rather than *α, β* to avoid collision with the heuristic coefficients in Equation (3).) For an MZ pair (1, 2): *G*_1_ = *G*_2_ and *E*_1_ ⊥ *E*_2_. Note 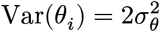; under this parameterization, *σ*_*θ*_ is the per-component scale factor, not the standard deviation of *θ*_*i*_ — note 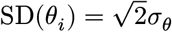. This convention separates the genetic (*G*) and environmental (*E*) components with symmetric unit-variance scaling.

Define extrinsic frailty 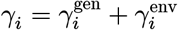 with *γ*^gen^ shared identically in MZ pairs, 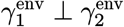, and Var(*γ*^gen^) = Var(*γ*^env^) = 1/2, so Var(*γ*) = 1. Assume Corr(*G, γ*^gen^) = *ρ* ∈ [−1, 1], with all other cross-components independent. The individual extrinsic hazard is *c*_*i*_ ≔ *m*_ex_ exp(*σ*_*γ*_*γ*_*i*_). (For derivational convenience, we absorb the centering constant 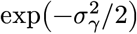 from the main text’s Equation (2) into *m*_ex_ here, so that 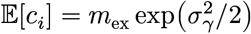 rather than *m*_ex_; this does not affect the inflation result because *m*_ex_ enters only as a fixed constant in the fitted model.)

**True hazard:**

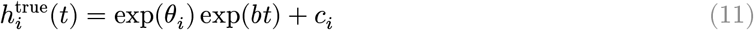

#### Fitted (misspecified) model

Shenhar’s fitted class uses 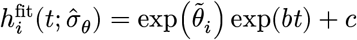 with 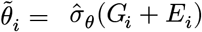 and constant *c* (no individual extrinsic variation). We take *c* to be fixed at the population mean, matching the procedure of calibrating *σ*_*θ*_ while holding *b* and *c* fixed. The calibration adjusts 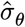 to match the observed MZ twin correlation *r*_MZ_ from the true DGP, then extrapolates by setting *c* = 0.

### B.2 Linearization via delta method

Let *L* = log *T*. The cumulative hazard under the true DGP is:

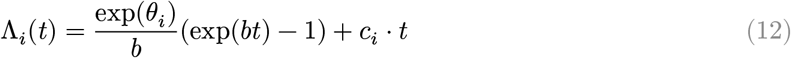

Death occurs at *T*_*i*_ where Λ_*i*_(*T*_*i*_) = *U*_*i*_, *U*_*i*_ ∼ Exp(1). Define:

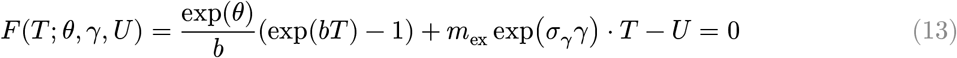

Since *∂F* /*∂T* = *h*^true^(*T*) > 0, the implicit function theorem guarantees *T* = *T* (*θ, γ, U*) is differentiable. Differentiating with respect to a generic parameter *φ* ∈ {*θ, γ*}:

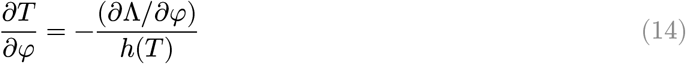

#### Intrinsic slope

With *∂*Λ/*∂θ* = Λ_int_(*T*), define:

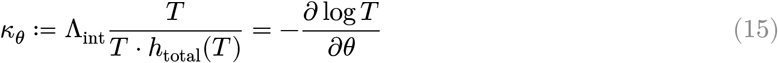

#### Extrinsic slope

By the chain rule (*∂*Λ/*∂γ* = *σ*_*γ*_ ⋅ Λ_ext_(*T*)), define *κ*_*γ*_ as the base extrinsic slope *without* the *σ*_*γ*_ factor:

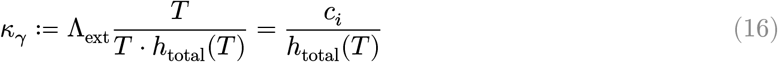

so that −*∂* log *T* /*∂γ* = *σ*_*γ*_*κ*_*γ*_.

#### Slope approximation

The coefficients *κ*_*θ*_ and *κ*_*γ*_ as defined above are functions of (*θ, γ, U*) through the realized death time *T*. In the first-order delta-method framework, we evaluate them at the linearization point (e.g., *θ* = *μ*_*θ*_, *γ* = 0, *U* = 1) and treat them as population-level constants in the subsequent covariance algebra. Under the true model, 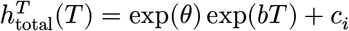; under the fitted model, 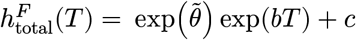. Define 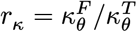; note *r*_*κ*_ → 1 as *m*_ex_/[exp(*θ*) exp(*bT*)] → 0. The analysis treats this difference as higher-order relative to the omitted-variable term in the empirically relevant regime.

#### First-order expansion

True DGP:

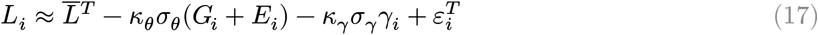

Fitted model:

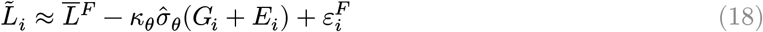

where *ε*^*T*^, *ε*^*F*^ capture higher-order terms and residual stochasticity from *U*.

### B.3 Variance-covariance decomposition for MZ pairs

#### Covariance normalization

Given *G* ∼ 𝒩(0, 1) and *γ*^gen^ ∼ 𝒩(0, 1/2) with Corr(*G, γ*^gen^) = *ρ*:

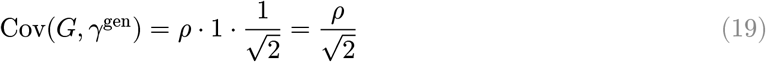

For an MZ pair, the shared components are *G* (identical) and *γ*^gen^ (identical); the non-shared components are *E*_1_ ⊥ *E*_2_ and 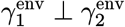.

#### True MZ covariance (*C*_true_)

Only shared components contribute:

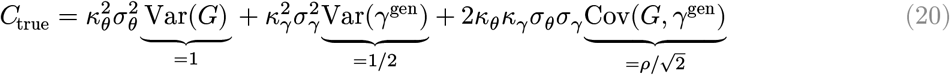

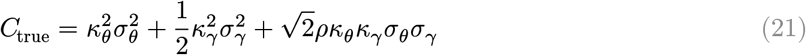

#### True DGP variance (*V*_true_)

All components contribute. Both *G* and *E* contribute to intrinsic variance (each with Var = 1, independent), giving a factor of 2:

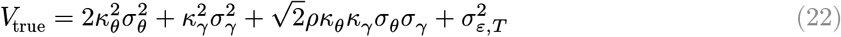

The cross-term 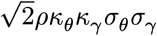 appears once (not doubled) because only *G* (not *E*) is correlated with *γ*^gen^, and *γ*^env^ is independent of everything.

#### Fitted model

With only 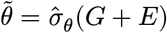 and no extrinsic variation:

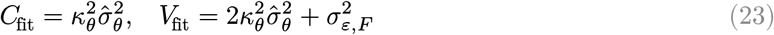

#### Key structural asymmetry

*C*_true_ contains extra terms from shared extrinsic frailty (*γ*^gen^) and pleiotropy (*ρ*) that *C*_fit_ lacks; *V*_true_ contains extra non-shared extrinsic variance that *V*_fit_ lacks. The net effect on *r*_MZ_ = *C*/*V* depends on relative magnitudes.

### B.4 Calibration equation and inflation condition

#### Lemma (existence and uniqueness)

In the fitted model, 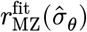 is strictly increasing in 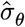 and takes values in (0, 1/2) for 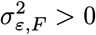. Hence a unique finite 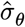 exists if and only if 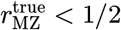, equivalently *V*_true_ − 2*C*_true_ > 0. We assume this throughout.

#### Correlation matching

The calibration sets 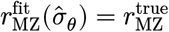:

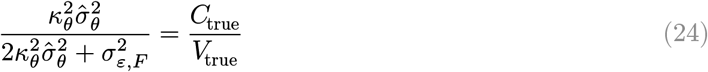

Let 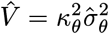. Cross-multiplying (using *V*_true_ − 2*C*_true_ > 0):

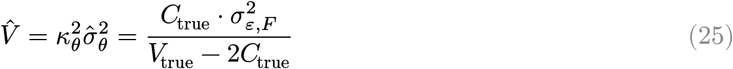

Define shorthands: 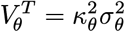 (intrinsic), 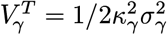 (extrinsic), 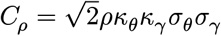 (pleiotropy). Then:

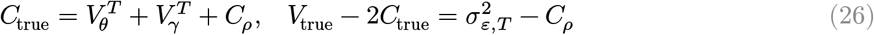

The feasibility condition requires 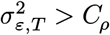. Even at the upper bounds of the anchored regime (*ρ* = 0.50, *κ*_*θ*_ ≈ 0.8, *κ*_*γ*_ ≈ 0.05, *σ*_*θ*_ ≈ 0.27, *σ*_*γ*_ = 0.65), 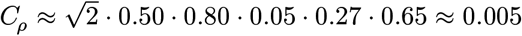, which is one to two orders of magnitude below typical residual log-lifespan variance; the condition is comfortably satisfied throughout the anchored parameter space. Substituting:

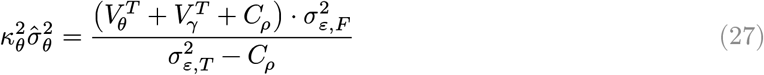

#### Inflation condition

Inflation 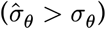 requires 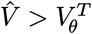. When 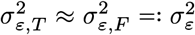 (valid when extrinsic hazard is small relative to intrinsic at typical death ages), this simplifies to:

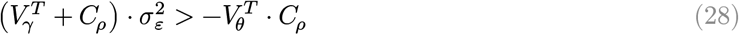

When *ρ* ≥ 0: *C*_*ρ*_ ≥ 0 and 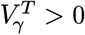, so the LHS is strictly positive and the RHS is ≤ 0. The inequality holds.

##### Proposition B1

**(local inflation under first-order approximation)**. *Under the first-order linearization of log-lifespan and when slope and residual-noise differences between the true and fitted models are second-order, omission of heritable extrinsic frailty (σ*_*γ*_ > 0*) with non-negative pleiotropy (ρ* ≥ 0*) inflates the calibrated intrinsic dispersion:*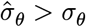.

#### Scope of the equal-noise approximation

The approximation 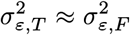 becomes quantitatively unreliable when extrinsic frailty variance is large — for example, at *σ*_*γ*_ ≈ 1.47 (the bridge analysis estimate that matches published infection-mortality heritability). In such regimes, the analytical bound isolates the direction and mechanism of bias, but the exact simulation results — not the first-order approximation — constitute the operative quantitative statement on bias magnitude.

For the case *ρ* < 0, see Corollary B4 below.

### B.5 Remark on concordant survivor truncation

Let *A* = {*T*_1_ > *t*_0_, *T*_2_ > *t*_0_} denote concordant survival to age *t*_0_. Truncation selects against high-frailty pairs. Because shared components (*G, γ*^gen^) are carried by both twins jointly while non-shared components (*E, γ*^env^) are selected independently for each twin, truncation tends to reduce non-shared variance relatively more than shared variance. Empirically, the sign of the bias is preserved and often amplified. Since the bias mechanism operates at the calibration step regardless of sample restriction, truncation is not required for the main result.

### B.6 Monotonicity of extrapolated *h*^2^(*σ*)

#### Proposition B2.

*Under the first-order approximation with fixed local slope κ and frailty-independent residual variance* 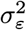, *the extrapolated Falconer heritability h*^2^(*σ*) = 2(*r*_MZ_ − *r*_DZ_) *is strictly increasing in σ for σ* > 0 *at the extrapolation point (c* = 0, *pure Gompertz)*.

**Proof**. Under pure Gompertz with *c* = 0, MZ twins share *G* (correlation 1) and DZ twins share *G* with correlation 0.5. Treating *κ* as a constant evaluated at the linearization point:

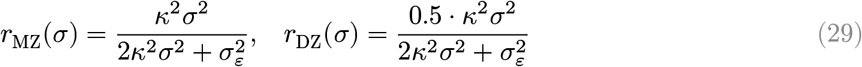

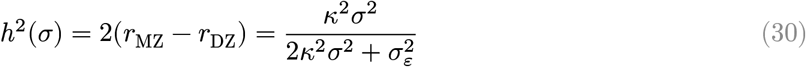

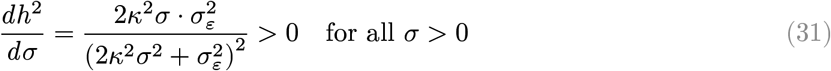

If calibration inflates 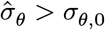 (Proposition B1), it inflates the extrapolated heritability estimate within the same first-order framework.

### B.7 Monotone-summary propagation

#### Corollary B3 (order propagation through monotone summaries)

*Let S* : (0, ∞) → ℝ *be strictly increasing on an interval containing* 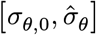. *If Proposition B1 gives* 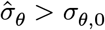, *then* 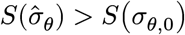.

**Proof**. Immediate from strict monotonicity.

#### Remark (portability and scope)

This corollary establishes a *portability principle*: the calibration-stage inflation of 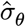 is the single upstream distortion; all downstream summary biases are consequences of it. Proposition B2 verifies the monotonicity condition for the specific case *S*(*σ*) = *h*^2^(*σ*). The same first-order algebra shows that *r*_MZ(*σ*)_ is strictly increasing in *σ* under the linearized one-component model. For concordance functionals (Kendall’s *τ*, Spearman’s *ρ*, concordance probability), monotonicity in the frailty scale is a standard property of Archimedean copulas induced by shared frailty models; see (Oakes, 1989) and (Hougaard, 2000) for the connection between frailty variance and concordance ordering.

For *composite* summaries constructed as differences of increasing functions — such as the ACE *A*-component *A*(*σ*) = 2{*r*_MZ(*σ*)_ − *r*_DZ(*σ*)_} — monotonicity of each component does not in general imply monotonicity of the composite. Corollary B3 applies once monotonicity of the *overall mapping σ* ↦ *S*(*σ*) has been established, analytically or over the relevant parameter range. In our simulations, *A*(*σ*) is increasing over the calibrated range; we treat this as empirically verified rather than as a consequence of B4 alone.

### B.8 Sign reversal under negative pleiotropy

#### Corollary B4

*If ρ* < 0 *and sufficiently negative, the omitted extrinsic component reduces the observed MZ correlation, causing the fitted model to underestimate intrinsic dispersion:* 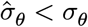.

Under the equal-residual-noise simplification 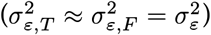, the exact inflation boundary is:

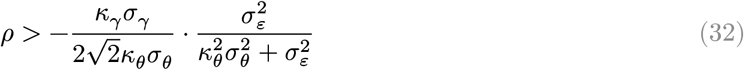

Downward bias occurs when *ρ* falls below this threshold. A sufficient condition for sign reversal (stronger but easier to interpret) that does not depend on 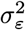 is 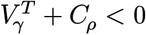:

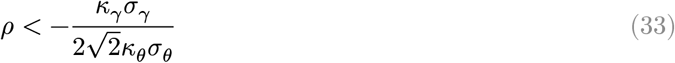

This is a **sufficient but not necessary** condition for sign reversal, corresponding to the high-noise limit 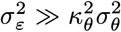 where the residual-dependent factor approaches 1. The exact boundary (first equation above) may be closer to zero than this sufficient condition; indeed, simulation confirms sign reversal near *ρ* ≈ 0 in some parameter regimes, consistent with the exact boundary being less negative than the sufficient condition.

### B.9 Surface-misfit diagnostic

The bivariate survival diagnostics in §4.5 show that the misspecified model’s joint twin survival surface exhibits structured residuals (ISE ratio 48×). The following identity clarifies the connection between parameter inflation and surface-fit deterioration.

#### Proposition B5

**(residual–sensitivity identity)**. *Let* 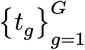 *be a grid on* (*t*_1_, *t*_2_)*-space with weights w*_*ij*_ > 0. *Let R*_*σ*_(*t*_*i*_, *t*_*j*_) *denote the model-implied dependence surface parametrized by σ, and R*_true_(*t*_*i*_, *t*_*j*_) *the true DGP surface. Define the weighted squared-error misfit*

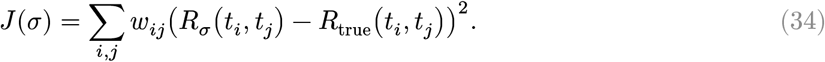

*If R*_*σ*_ *is differentiable in σ, then*

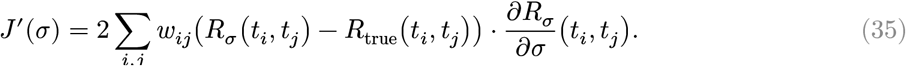

*In particular, J*^′^(*σ*_*θ*,0_) > 0 *if and only if the inner product of the residual field* 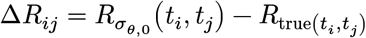 *and the sensitivity field* 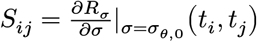 *is positive:* 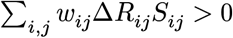.

**Proof**. Differentiate the finite sum term by term.

#### Interpretation

In the one-component Gompertz–Makeham model, increasing *σ* concentrates concordance in the late-life region where intrinsic aging dominates — the sensitivity field *S*_*ij*_ is large and positive at high ages. The omitted extrinsic channel, however, contributes concordance preferentially at *younger* ages (where the constant *m*_ex_ hazard channel is relatively strongest). The misspecified model, lacking this channel, underestimates joint survival at younger ages (Δ*R*_*ij*_ < 0) and slightly overestimates it at the oldest ages (Δ*R*_*ij*_ > 0 in the small upper-right region where inflated *σ*_*θ*_ over-compensates). The inner product in Equation (35) reflects this structured misalignment.

#### Remark (local quadratic approximation)

Linearizing 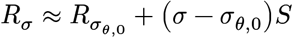 yields the approximate *L*^2^-minimizer

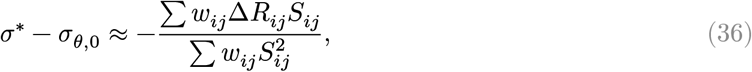

which is the projection of the residual field onto the sensitivity direction. This connects the parameter shift to the geometry of the surface misfit.

#### Caveats

(i) This is a local first-order result; it does not characterize the global behavior of *J*(*σ*). (ii) Our calibrated 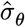 is defined by moment matching (*r*_MZ_ target), not by minimizing *J*; Proposition B5 is therefore used as *diagnostic geometry* — explaining why the ISE worsens under inflation — not as an estimating equation. (iii) The weights *w*_*ij*_ correspond to the grid spacing and any masking thresholds used in §3.5; changing the weighting changes the inner product.

### B.10 Interpretation under empirically motivated parameter regimes

The theoretical analysis is a first-order (delta-method) result. Its qualitative predictions are confirmed by exact simulation across the biologically anchored parameter space. The manuscript’s sensitivity regime uses *σ*_*γ*_ ∈ [0.30, 0.65], *ρ* ∈ [0.20, 0.50], and *m*_ex_ ∈ [0.001, 0.012]. Since *ρ* > 0 throughout, Proposition B1 predicts upward bias in 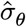 — confirmed quantitatively at 5–14 pp across all three model families. The sign-reversal threshold (Corollary B4) requires negative *ρ*, outside the empirically motivated range. The first-order analysis provides the direction and mechanism; the simulation provides exact magnitudes beyond the linear approximation.

### B.11 Summary of formal results

**Proposition B1 (local inflation)**. Omission of heritable extrinsic frailty (*σ*_*γ*_ > 0) with non-negative pleiotropy (*ρ* ≥ 0) inflates the calibrated intrinsic dispersion under first-order linearization: 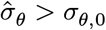. **Proposition B2 (monotonicity of Falconer)**. The extrapolated Falconer *h*^2^(*σ*) is strictly increasing in *σ*. Combined with B1, this implies upward bias in the extrapolated intrinsic heritability.

**Corollary B3 (monotone-summary propagation)**. Any strictly increasing dependence summary *S*(*σ*) inherits the inflation: 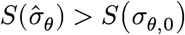. Proposition B2 is the specific case *S* = *h*^2^; the result is not specific to Falconer’s formula.

**Remark (truncation)**. Concordant survivor truncation is not required for the bias but empirically amplifies it.

**Corollary B4 (sign reversal)**. Sufficiently negative pleiotropy reverses the bias direction.

**Proposition B5 (surface-misfit diagnostic)**. The slope of the weighted squared-error surface misfit equals the inner product of the residual and sensitivity fields; positive alignment implies that parameter inflation worsens the bivariate fit.

Simulations verify these qualitative predictions beyond the local approximation, across all three model families and throughout the biologically anchored parameter space.

## C Appendix: Computer-verified algebraic core

This appendix documents the machine-checked formalization of the algebraic core of Section B — the deductive chain from the first-order moment expressions (*C*_true_, *V*_true_, *C*_fit_, *V*_fit_) to the inflation inequality (Proposition B1), the monotonicity result (Proposition B2), and the sign-reversal threshold (Corollary B4). The formalization uses the Lean 4 interactive theorem prover (v4.29.0) with the Mathlib4 mathematical library, and compiles with **zero** `sorry` (unproven assertions) and **zero** `axiom` declarations beyond Lean’s foundational type theory.

### C.1 Scope and methodology

The proofs in Section B rest on two epistemic layers: (i) a first-order (delta-method) linearization of log-lifespan, yielding moment expressions for *C*_true_, *V*_true_, *C*_fit_, and *V*_fit_; and (ii) algebraic manipulation of those moment expressions to establish the inflation inequality. Layer (i) is an asymptotic approximation accepted throughout the survival analysis literature. Layer (ii) is where algebraic errors could silently invalidate the claim.

We machine-verified layer (ii). Every algebraic step from the moment definitions through the final inequality is checked by Lean’s kernel — a small, trusted code base that verifies each deduction.

The **DEFINED** items are the standard delta-method framework. Their correctness is the premise of Section B‘s “first-order approximation” qualifier and is validated by exact simulation throughout the main text. The **VERIFIED** items constitute the algebraic core — the deductive chain from moment definitions to the inflation conclusion — and are checked with zero gaps.

### C.2 Formal statement correspondence

Table 11 maps each manuscript result to its Lean theorem name and source file.

**Table 11.** Correspondence between manuscript propositions and Lean 4 theorem names.

| Manuscript | Lean 4 theorem | File |
| --- | --- | --- |
| Proposition B2 | corrFun_strictMonoOn | Monotonicity.lean |
| Lemma (existence) | lemma_rMZ_strictMono, lemma_rMZ_range | Monotonicity.lean |
| Eq. (calibration) | calibrated_variance_expanded | Calibration.lean |
| Proposition B1 | prop_B1_inflation | Inflation.lean |
| Corollary B4 | corollary_B3_deflation | SignReversal.lean |
| Eq. $V_{\text{true}} - 2C_{\text{true}}$ | Params.V_true_minus_2C | Moments.lean |
| Feasibility | Params.feasible_iff | Moments.lean |

### C.3 Parameter structure

All results are parameterized over a structure encoding the sensitivity coefficients, frailty scales, residual variance, and pleiotropy correlation with built-in positivity and boundedness constraints:

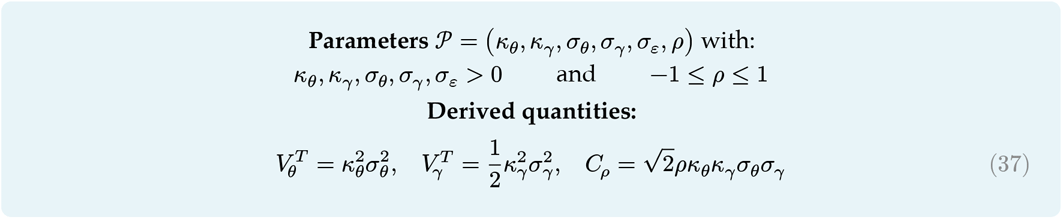

### C.4 Machine-checked results

#### C.4.1 Proposition B2 (monotonicity)

**Formal statement**. For all *κ* > 0, *ε* > 0:

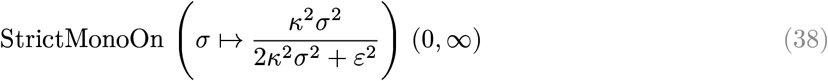

**Proof method**. Algebraic: for 0 < *x* < *y*, cross-multiply positive denominators and reduce to *κ*^2^*ε*^2^*x*^2^ < *κ*^2^*ε*^2^*y*^2^, which follows from *x*^2^ < *y*^2^. No calculus (derivatives, continuity) required.

**Corollary**. The function takes values in (0, 1/2), establishing existence and uniqueness of the calibration solution.

#### C.4.2 Proposition B1 (inflation)

**Formal statement**. Given parameters 𝒫 with *ρ* ≥ 0, feasibility 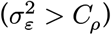, a calibrated value 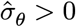 satisfying

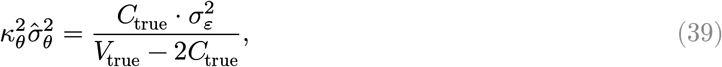

then 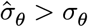.

**Proof method**. Three steps, each machine-checked:

1. **Key identity** (proved by `ring`):

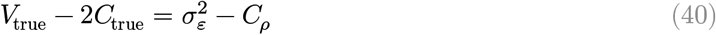
2. **Numerator positivity** (proved by `nlinarith`):

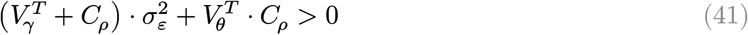

Since 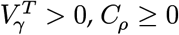 (when *ρ* ≥ 0), and 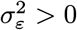.
3. **Denominator clearing** (proved by lt_div_iff_0_ + `nlinarith`):

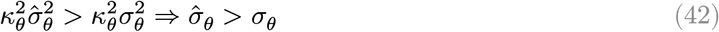

#### C.4.3 Corollary B4 (sign reversal)

**Formal statement**. If *ρ* < *ρ*_crit_ where

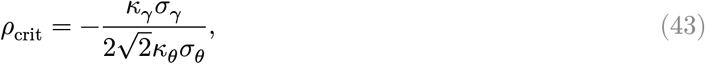

then under feasibility and the same calibration equation, 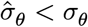.

**Proof method**. The threshold *ρ*_crit_ makes 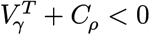, which (combined with *C* < 0) reverses the numerator sign. The proof clears the division in the threshold using lt_div_iff_0_, then multiplies the cleared inequality by *κ*_*γ*_*σ*_*γ*_ > 0 to establish the bound.

### C.5 Axiomatization boundary

The formalization draws a deliberate boundary between what is proved and what is assumed:

**Proved (layer ii)**. Every algebraic manipulation from the moment definitions (*C*_true_, *V*_true_ as functions of 𝒫) through the calibration equation to the final inflation inequality and sign-reversal threshold. This includes all intermediate identities, positivity lemmas, and the denominator-clearing steps.

**Defined (layer i)**. The moment expressions themselves:

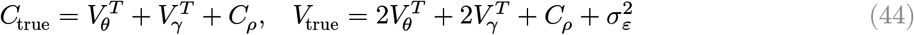

These are the output of the delta-method linearization (Section B, §B.2–B.3). We encode them as Lean definitions rather than deriving them from a measure-theoretic probability space, because Mathlib4 does not yet provide multivariate normal distributions with specified correlation structure. The equalnoise approximation 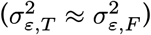 is implicit in the formalization — the parameter *σ*_*ε*_ represents both residual variances.

This axiomatization is conservative: it means a bug in the covariance algebra of §B.3 would not be caught by the formalization, but any error in the **deductive chain from those covariances to the inflation conclusion** would be. The covariance derivation is itself a short, checkable calculation that the simulation confirms numerically.

### C.6 Reproduction instructions

The formalization is self-contained in the `CalibrationInflation/` directory of the companion Lean 4 project, available at github.com/biostochastics/extrinsic-frailty-heritability under the `lean/` subdirectory.

**Prerequisites:** Lean 4.29.0 (via elan), Mathlib4 (fetched automatically by Lake).

#### Build

git clone https://github.com/biostochastics/extrinsic-frailty-heritability.git cd extrinsic-frailty-heritability/lean

`lake build CalibrationInflation # ∼45s; downloads Mathlib on first run`

#### Verify zero sorry

`grep -rn “sorry” CalibrationInflation/ # expect: no output`

`grep -rn “^axiom” CalibrationInflation/ # expect: no output`

#### File manifest

**File manifest:**
|  |  |
| --- | --- |
| **Params.lean** | **Parameter structure, derived quantities, positivity/nonzero lemmas** |
| **Monotonicity.lean** | Prop B2, Lemma: algebraic monotonicity of $\kappa^2\sigma^2/(2\kappa^2\sigma^2 + \varepsilon^2)$ |
| **Moments.lean** | Moment definitions ($C_{\text{true}}, V_{\text{true}}$), feasibility, calibration equation |
| **Calibration.lean** | Explicit solution existence, expanded form of calibrated variance |
| **Inflation.lean** | Prop B1: numerator positivity, variance inflation, $\hat{\sigma}_\theta > \sigma_\theta$ |
| **SignReversal.lean** | Corollary B4: threshold derivation, deflation under negative pleiotropy |

**Lean version:** leanprover/lean4:v4.29.0-rc8 **Mathlib:** latest via lake update

### C.7 Limitations

The formalization verifies the algebraic core under the equal-noise approximation. It does not verify: (a) the delta-method linearization itself, (b) the covariance decomposition from the MZ twin probability space, or (c) the Gompertz-Makeham DGP specification. These are validated by exact simulation in the main text. Extending the formalization to a full measure-theoretic derivation would require Mathlib infrastructure for multivariate normal distributions with specified correlation matrices, which is not yet available.

## Footnotes

1 The mechanism generalizes to any source of familial concordance in extrinsic susceptibility, including shared environment. The Falconer decomposition separates genetic from shared-environmental effects through MZ/DZ differences, but the calibration step conflates any within-pair extrinsic concordance regardless of its source.

2 We adopt the compensatory form 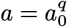, consistent with Shenhar et al.‘s preprint; see Appendix Section A.4 for implementation details.

## Notes

https://github.com/biostochastics/extrinsic-frailty-heritability

### Competing Interest Statement

The authors have declared no competing interest.

## References

Clayton, D. G. (1978). A model for association in bivariate life tables and its application in epidemiological studies of familial tendency in chronic disease incidence. Biometrika, 65(1), 141–151. 10.1093/biomet/65.1.141

Franceschi, C., Bonafè, M., & Valensin, S. (2000). Inflamm-aging: an evolutionary perspective on immunosenescence. Annals of the New York Academy of Sciences, 908, 244–254. 10.1111/j.1749-6632.2000.tb06651.x

Gasparini, A., Clements, M. S., Abrams, K. R., & Crowther, M. J. (2019). Impact of model misspecification in shared frailty survival models. Statistics in Medicine, 38(23), 4477–4502. 10.1002/sim.8309

Hamilton, F. (2026). Misclassification of heritable mortality undermines estimates of intrinsic life span heritability. Medrxiv. 10.64898/2026.02.26.26347172

Hougaard, P. (1995). Frailty models for survival data. Lifetime Data Analysis, 1(3), 255–273. 10.1007/BF00985760

Hougaard, P. (2000). Analysis of Multivariate Survival Data. Springer. 10.1007/978-1-4612-1304-8

Joshi, P. K., Pirastu, N., Kentistou, K. A., Fischer, K., Hofer, E., Schraut, K. E., Clark, D. W., & Wilson, J. F. (2017). Genome-wide meta-analysis associates HLA-DQA1/DRB1 and LPA and lifestyle factors with human longevity. Nature Communications, 8, 910. 10.1038/s41467-017-00934-5

Oakes, D. (1989). Bivariate survival models induced by frailties. Journal of the American Statistical Association, 84(406), 487–493. 10.1080/01621459.1989.10478795

Obel, N., Christensen, K., Petersen, I., Sørensen, T. I. A., & Skytthe, A. (2010). Genetic and environmental influences on risk of death due to infections assessed in Danish twins, 1943–2001. American Journal of Epidemiology, 171(9), 1007–1013. 10.1093/aje/kwq037

Petersen, L., Andersen, P. K., & Sørensen, T. I. A. (2010). Genetic influences on incidence and case-fatality of infectious disease. Plos ONE, 5(5), e10603. 10.1371/journal.pone.0010603

Qiu, S. (2025). COVID-19 infection and longevity: an observational and Mendelian randomization study. Journal of Translational Medicine, 23, 283. 10.1186/s12967-024-05932-y

Rosa, M., Chignon, A., & Li, Z. (2019). A Mendelian randomization study of IL6 signaling in cardiovascular diseases, immune-related disorders and longevity. Npj Genomic Medicine, 4, 23. 10.1038/s41525-019-0097-4

Ruby, J. G., Wright, K. M., & Rand, K. A. (2018). Estimates of the heritability of human longevity are substantially inflated due to assortative mating. Genetics, 210(3), 1109–1124. 10.1534/genetics.118.301613

Schurz, H., Naranbhai, V., Yates, T. A., Gilchrist, J. J., Parks, T., Dodd, P. J., Möller, M., Hoal, E. G., Morris, A. P., & Hill, A. V. S. (2024). Multi-ancestry meta-analysis of host genetic susceptibility to tuberculosis identifies shared genetic architecture. Elife, 13, e84394. 10.7554/eLife.84394

Shenhar, B., Pridham, G. Oliveira, T. Lopes de, Raz, N., Yang, Y., Deelen, J., Hägg, S., & Alon, U. (2026). Heritability of intrinsic human life span is about 50% when confounding factors are addressed. Science, 391(6784), 504–510. 10.1126/science.adz1187

Sørensen, T. I. A., Nielsen, G. G., Andersen, P. K., & Teasdale, T. W. (1988). Genetic and environmental influences on premature death in adult adoptees. New England Journal of Medicine, 318(12), 727–732. 10.1056/NEJM198803243181202

Tervi, A., Junna, N., Broberg, M., Jones, S. E., Strausz, S., Kreivi, H.-R., Heckman, C. A., & Ollila, H. M. (2023). Large registry-based analysis of genetic predisposition to tuberculosis identifies genetic risk factors at HLA. Human Molecular Genetics, 32(1), 161–172. 10.1093/hmg/ddac212

Vaupel, J. W., Manton, K. G., & Stallard, E. (1979). The impact of heterogeneity in individual frailty on the dynamics of mortality. Demography, 16(3), 439–454. 10.2307/2061224

White, H. (1982). Maximum likelihood estimation of misspecified models. Econometrica, 50(1), 1–25. 10.2307/1912526

Wienke, A., Holm, N. V., Christensen, K., Skytthe, A., Vaupel, J. W., & Yashin, A. I. (2003). The heritability of cause-specific mortality: a correlated gamma-frailty model applied to mortality due to respiratory diseases in Danish twins born 1870–1930. Statistics in Medicine, 22(24), 3873–3887. 10.1002/sim.1669

Wienke, A., Holm, N. V., Skytthe, A., & Yashin, A. I. (2001). The heritability of mortality due to heart diseases: a correlated frailty model applied to Danish twins. Twin Research, 4(4), 266–274. 10.1375/1369052012399

Yashin, A. I., & Iachine, I. A. (1995). Genetic analysis of durations: correlated frailty model applied to survival of Danish twins. Genetic Epidemiology, 12(5), 529–538. 10.1002/gepi.1370120510

Yashin, A. I., Iachine, I. A., & Harris, J. R. (1999). Half of the variation in susceptibility to mortality is genetic: findings from Swedish twin survival data. Behavior Genetics, 29(1), 11–19. 10.1023/A:1021481620934

Ying, K., Zhai, R., & Pyrkov, T. V. (2021). Genetic and phenotypic analysis of the causal relationship between aging and COVID-19. Communications Medicine, 1, 35. 10.1038/s43856-021-00033-z

